# PartiDesign: A structure-sequence co-design paradigm for practical all-to-all biomolecules design

**DOI:** 10.64898/2026.11.03.763860

**Authors:** Zichang Jin, Xujun Zhang, Yu Kang, Tingjun Hou

## Abstract

Engineering interactions among biomolecules offers a route to creating new biological functions. Although unified models have advanced biomolecular structure prediction and design, efficient and controllable generation across molecular modalities remains challenging. A key challenge is to couple three-dimensional structure generation with sequence or chemical-identity design across chemically diverse systems. Here, we introduce PartiDesign, a unified structure–sequence co-design model that extends the partially latent framework of La-Proteina to proteins, nucleic acids and small molecules. PartiDesign jointly generates molecular structures and their corresponding sequences or chemical identities, while providing fine-grained control over secondary structure, chemical bonding and amino acid preferences. Across diverse benchmarks for the de novo design of proteins, nucleic acids and small molecules against specified targets, PartiDesign achieved the highest estimated successful-design throughput among the evaluated methods on most benchmarks under the adopted compute-budget protocols. Its configurable constraints further support cyclic peptide and antibody design, allowing generation to be tailored to specific structural and chemical requirements. PartiDesign was developed using 128 Huawei Ascend 910B accelerators, each with 64 GB of memory. The source code is available at https://github.com/MegaGatlingPea/PartiDesign-official, and a freely accessible web server at https://cadd.zju.edu.cn/partidesign/ enables users to generate binding partners without programming expertise. Together, these capabilities provide a versatile and accessible framework for all-to-all biomolecular design, with applications in molecular engineering and drug discovery.

## 1 Introduction

Biological systems depend on interactions among proteins, nucleic acids and small molecules to coordinate processes ranging from signal transduction and metabolism to gene regulation. Despite differences in size, chemical composition and structural organization, these molecular classes interact through shared physicochemical principles, providing a common foundation for structure prediction and rational molecular design. Historically, however, AI-based approaches have largely developed around individual molecular classes or interaction types. Representative examples include RAPiDock for protein–peptide docking ^1^ and KarmaDock for protein–ligand docking ^2^, alongside RFdiffusion for protein generation ^3^, ResGen for protein-conditioned small-molecule generation ^4^ and RhoDesign for structure-conditioned RNA sequence design ^5^. These methods have advanced their respective tasks, while relying on representations and workflows tailored to particular molecular systems.

AlphaFold 3^6^ demonstrated the potential of unified biomolecular modelling by predicting complexes containing proteins, nucleic acids, small molecules and other chemical components within a single framework, with substantial improvements over specialized methods across a broad range of benchmarks. This advance motivates the development of generative models that can apply a similarly unified treatment to molecular design. The transition from prediction to design introduces an additional requirement: molecular identities must be determined together with three-dimensional configurations compatible with a specified target or functional constraint. A general design framework must therefore accommodate both the shared physical basis of molecular interactions and the distinct chemical organization of different molecular classes.

Toward this goal, we previously co-developed ODesign ^7^, a unified generative framework for biomolecular design across molecular modalities and conditioning requirements. ODesign adopts a two-stage pipeline that combines an AlphaFold 3–like structural generator with a multimodal inverse-folding module to assign sequences or atom types to generated scaffolds. Across 11 benchmark tasks spanning proteins, RNA, DNA and small molecules, ODesign outperformed the evaluated modality-specific methods and increased design throughput, demonstrating the feasibility of broad multimodal generation within a common framework.

In parallel, advances in protein design have expanded the range of accessible targets and design objectives. PXDesign ^8^, BoltzGen ^9^ and BindCraft2 ^10^ support increasingly diverse protein and peptide design workflows, with BoltzGen accommodating targets that include proteins, nucleic acids and small molecules. RFdiffusion3 and its nucleic-acid extensions ^11,12^ further broaden atomistic design to include RNA, DNA and hybrid protein– nucleic acid systems. Alongside these developments, La-Proteina ^13^ introduced a partially latent flow-matching framework that jointly generates protein sequence and atomistic structure by explicitly modelling backbone coordinates and encoding sequence identity and remaining atomic details in continuous latent variables. Proteina-Complexa ^14^ extends this approach to conditional protein binder design. These advances suggest that partially latent representations offer a promising basis for integrating molecular identity and geometry in a generative model.

Building on these advances, we identify three priorities for improving the practical utility of unified biomolecular design. First, the computational demands of ODesign’s atomistic, AlphaFold 3–like architecture motivate a more efficient generative representation. Second, its separation of structure generation from sequence or atom-type assignment makes the final output dependent on the capabilities of the downstream design module in each modality. Differences in reconstruction performance can therefore limit consistency across molecular classes. Third, although ODesign supports conditioning at molecular, motif and atomic levels, its control over properties such as secondary-structure composition remains limited. More flexible control over structural features, chemical connectivity and residue preferences would facilitate adaptation to practical tasks, including cyclic peptide and antibody design. Together, these considerations motivate a framework that combines multimodal coverage with joint generation and explicit design constraints.

Here, we introduce PartiDesign, a unified structure–sequence co-design model for practical all-to-all biomolecular design. Inspired by La-Proteina, PartiDesign extends partially latent generation from proteins to nucleic acids and small molecules, jointly modelling three-dimensional structures and their corresponding sequences or chemical identities. This formulation brings structural and chemical generation into a common framework, addressing the separation between these processes in two-stage pipelines. PartiDesign also incorporates fine-grained controls over secondary structure, chemical bonding and amino acid preferences, enabling users to tailor generation to task-specific requirements, including those associated with cyclic peptides and antibodies. We systematically evaluate PartiDesign across diverse benchmarks for the de novo design of proteins, nucleic acids and small molecules against specified targets. PartiDesign achieves the highest estimated successful-design throughput among the evaluated methods on most benchmarks under the adopted compute-budget protocols, supporting the practical value of multimodal co-design. By providing both source code and a freely accessible web server, we make these capabilities available to users with varying levels of computational expertise and establish a flexible platform for biomolecular engineering.

## 2 PartiDesign framework

PartiDesign couples a variational autoencoder with a flow-matching model to generate molecular identity and structure jointly (Fig. 1). The autoencoder learns a local representation of all-atom molecular environments and reconstructs polymer identities, atomic coordinates and ligand chemical graphs. With the autoencoder frozen, the flow matcher learns to transport Gaussian noise to the joint space of token centers and local latents. During training, separate noise levels perturb these two components, allowing the model to learn global arrangement and local molecular content across their respective noise conditions. Given these perturbed representations, the denoiser uses pair-biased attention to integrate information across all design and context tokens. The resulting joint updates couple local identity and geometry to the surrounding molecular arrangement, enabling sequence and structure to be generated together.

**Figure 1.**
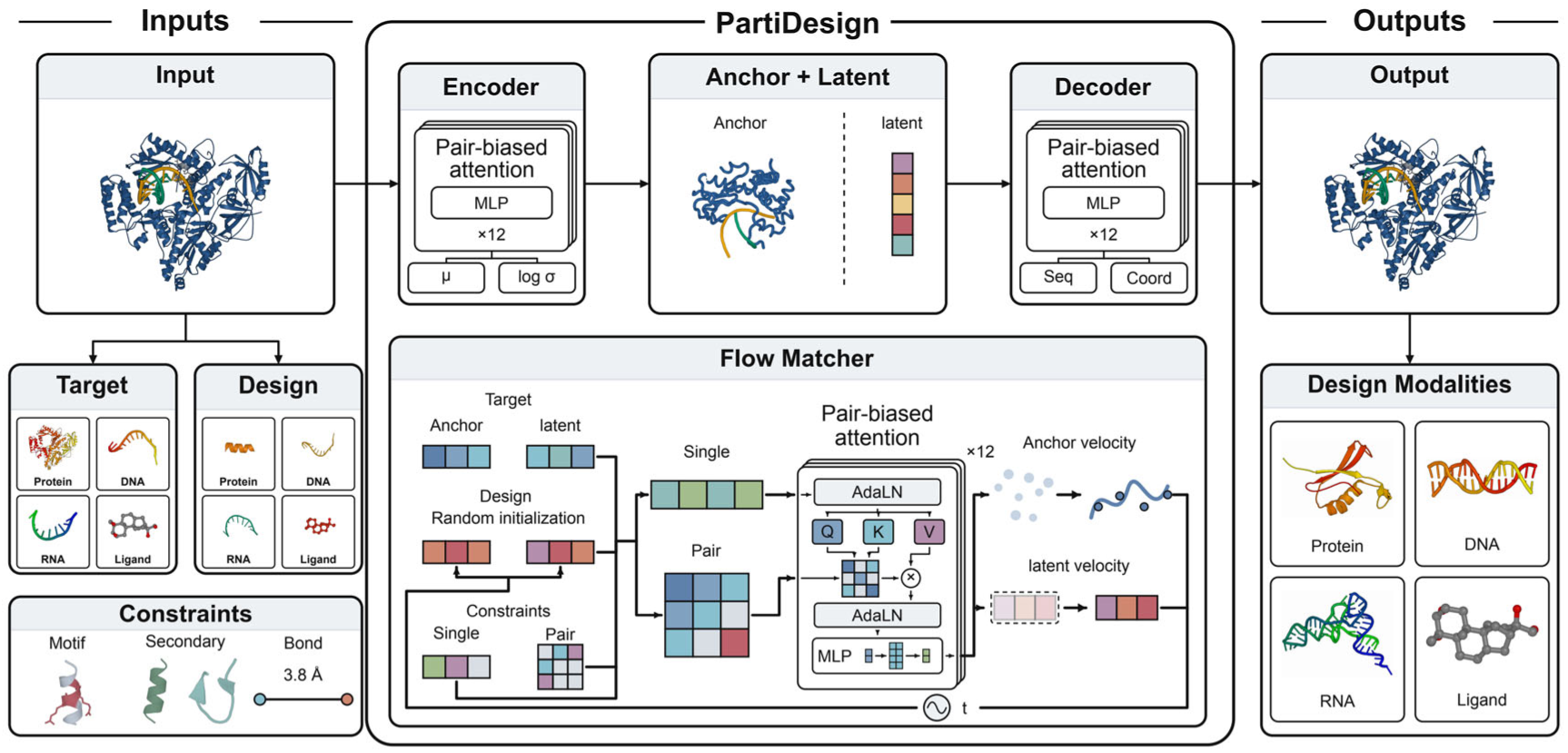
PartiDesign framework. Proteins, RNA and DNA are represented by residue tokens, whereas each ligand heavy atom forms a token. Each token comprises a three-dimensional center and an eight-dimensional local latent. Token centers describe the global molecular arrangement; local latents encode molecular identity and local atomic geometry. A variational autoencoder maps all-atom inputs to local latents and decodes the generated representation into molecular identities and coordinates. The flow matcher jointly generates token centers and latents, conditioned on molecular context, sequence, geometry, hierarchical secondary structure and chemical connectivity. This shared token-center *×* local-latent representation supports de novo generation and motif-based redesign across modalities, with user-specified conditions narrowing the design space.

User-specified context, sequence, structural and chemical conditions narrow the design space for de novo generation and motif-based redesign. A supplied protein, nucleic acid or small molecule provides the context for one or more jointly generated chains, and receptor hotspots focus generation on a chosen interaction site. Within this context, fixed identities retain residues required for recognition or chemistry, while backbone, full-residue and partial-atom motifs retain selected structural features. Contigs connect retained segments through newly designed regions.

The dual-crosslinked peptide in Fig. 2 illustrates how these conditions operate together. A protein receptor supplies the molecular context for an 18-residue peptide. Fixed cysteine and MK8 identities define the crosslink anchors, while chain-level *α* guidance and local helix conditions specify the intended conformation. Named atom endpoints and bond orders then specify a disulfide and a hydrocarbon staple within the same chain. Hydrocarbon crosslinks can increase peptide helicity and resistance to proteolysis^15^, motivating the combined control of sequence, conformation and connectivity in this example.

**Figure 2.**
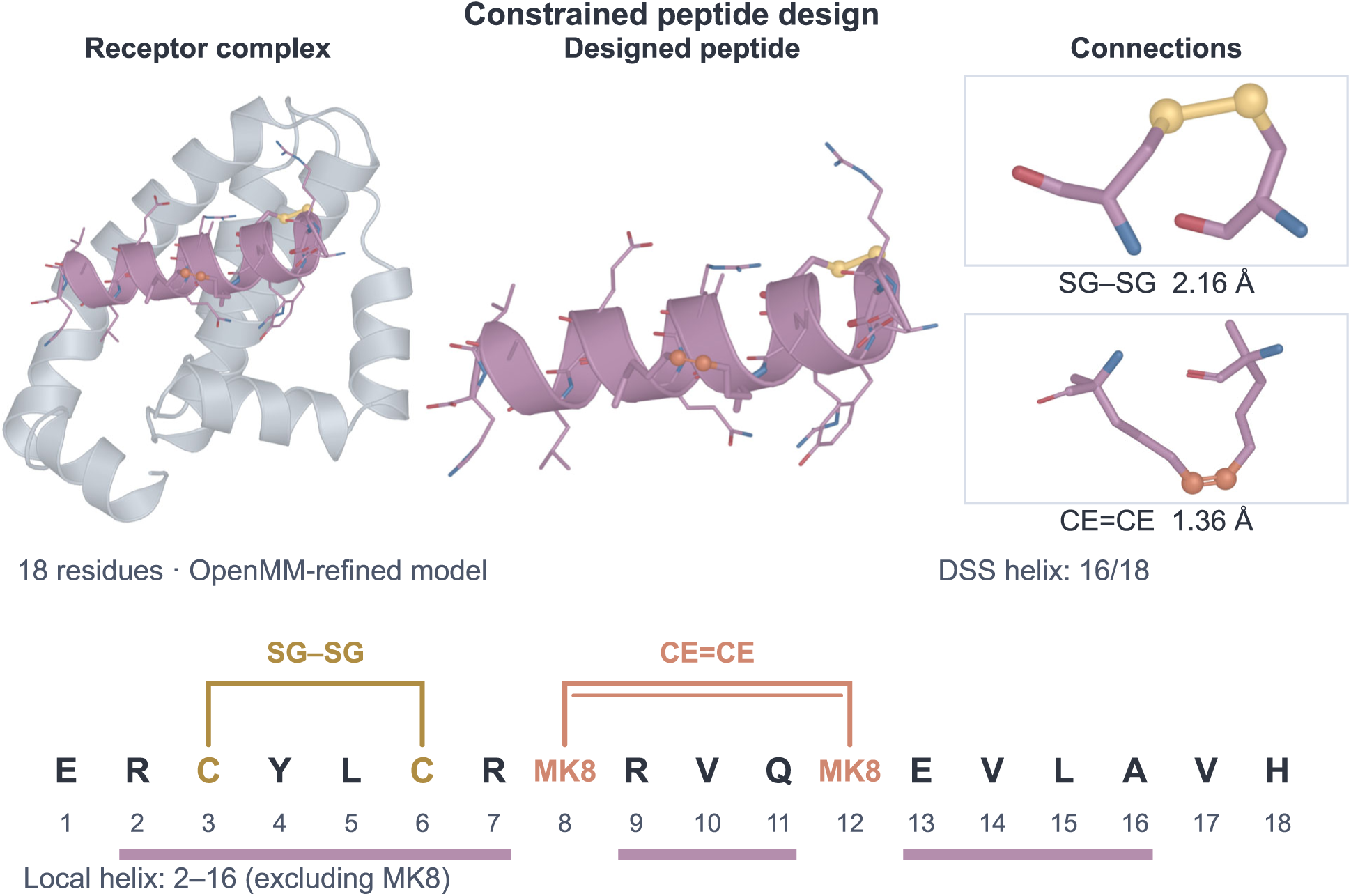
Combining receptor, sequence, structural and chemical conditions. The receptor complex, designed peptide and enlarged connections show the same refined output. The sequence diagram identifies the Cys3–Cys6 disulfide and MK8 CE8=CE12 staple; colored bars mark the locally specified helical segments. Fixed identities, helical geometry and both crosslinks are supplied in one design request.

The same controls extend beyond this example. Chain-level secondary-structure conditions guide proteins toward *α*- or *β*-rich folds and nucleic acids toward paired or unpaired structures; finer conditions specify local structural states, adjacency between protein secondary-structure blocks and explicit nucleotide pairs. Atom-level connections also encode ring closures and covalent links between molecular components. A supplied ligand graph can retain its chemical identity while its pose or surrounding protein pocket is designed, and partial-atom motifs specify functional atoms around which the remaining structure is generated.

Sequence control can also reflect experimental requirements. Token-identity omission excludes selected identities at free polymer positions. For example, excluding free cysteines reduces opportunities for unwanted intermolecular disulfide crosslinking, while cysteines required by a motif or an intended disulfide remain fixed (Fig. 3)^16^. Nucleotide exclusions similarly allow a DNA or RNA library to use a chosen alphabet. These exclusions therefore complement the structural and chemical conditions of a design request.

**Figure 3.**
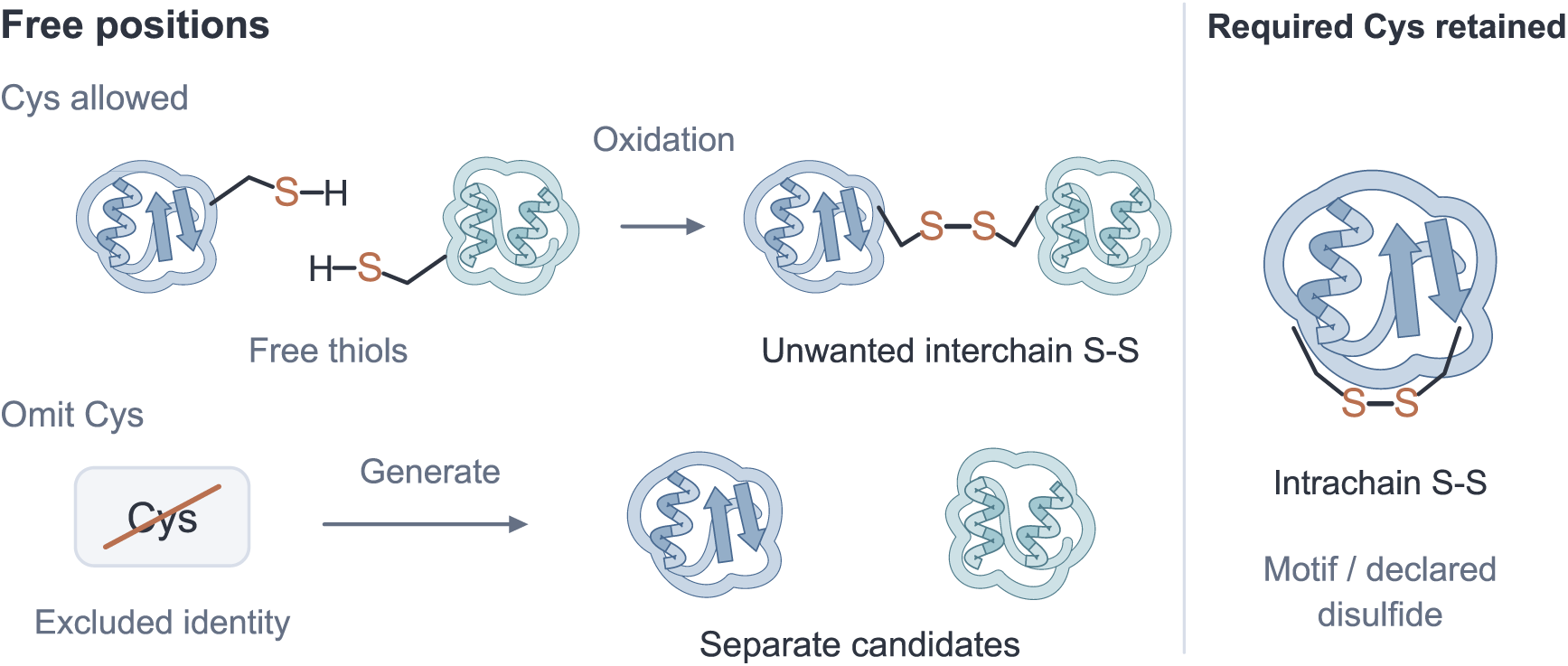
Sequence exclusions retain intended chemistry. The schematic contrasts free cysteines that can form an unwanted interchain disulfide with designs that exclude cysteine at free positions. Residue positions remain available for other identities. Required cysteines in a motif or declared intrachain disulfide are retained.

After generation, optional Best-of-*N* search produces multiple candidates for the same task and length specification, then retains designs according to an application-specific reward. PartiScore uses an MDN network to score molecular interfaces from interatomic contact geometry; refolding rewards instead assess sequence–structure consistency. All-atom relaxation refines generated coordinates before final scoring. Figure 4 illustrates this selection step through PartiScore ranking of refined peptide-docking candidates.

**Figure 4.**
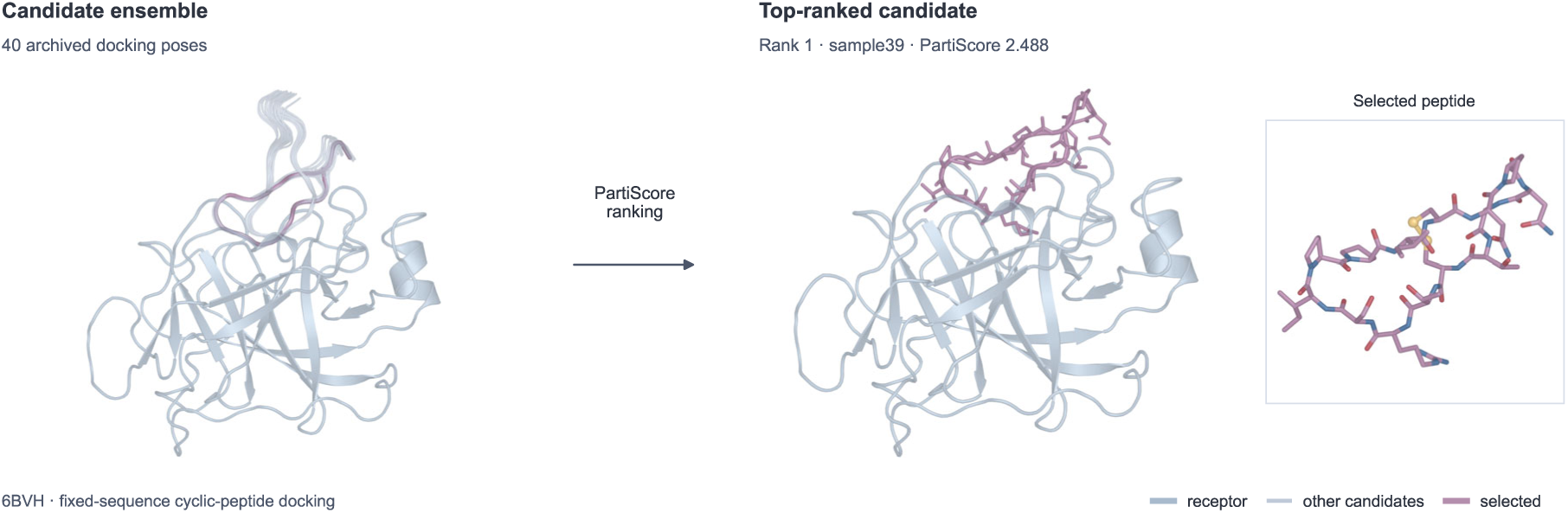
Selecting a refined peptide pose with PartiScore. Forty fixed-sequence cyclic-peptide docking candidates for 6BVH are ranked by PartiScore. The highest-scoring refined candidate is shown in receptor context and enlarged; yellow atoms identify the disulfide.

Model training and the PartiScore MDN are described in Sections A and B.

## 3 Evaluation strategy and computational budget

We combine task-specific passing rates with computational cost to estimate the number of passing candidates within a fixed budget. Tasks are grouped by the designed molecular modality and molecular context, which determine the predictor and filters (Fig. 5).

**Figure 5.**
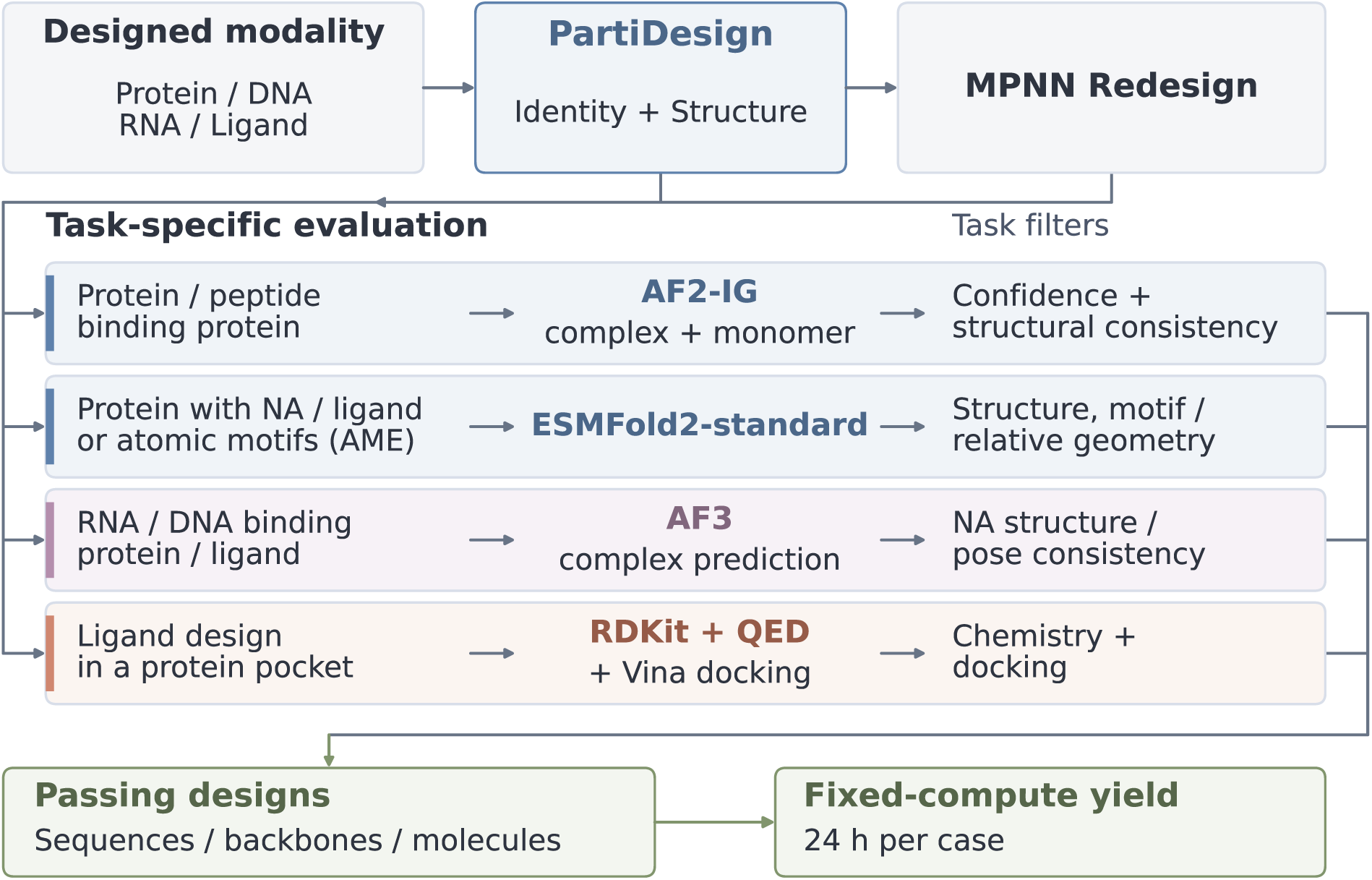
Evaluation follows the designed modality and its molecular context. PartiDesign generates molecular identities and structures, with optional sequence redesign for polymers. Each task then uses its designated structure predictor or chemical screen. Passing sequences, successful backbones and passing molecules are counted according to the task protocol; measured compute costs determine the fixed-budget yield.

### 3.1 Choosing a structural reference for each design modality

Protein–protein and peptide–protein designs are evaluated with AlphaFold2 initial guess (AF2-IG). BindCraft provides experimental precedent for selecting protein binders using AlphaFold-based confidence and structural-consistency measures ^17^. We retain the benchmark-specific thresholds and the respective ProtDBench and peptide protocols for predicting the complex and isolated designed chain. The resulting screens assess interface confidence and agreement between bound and unbound conformations.

For proteins designed around nucleic acids or small molecules, and for atomic-motif enzyme scaffolding, we use ESMFold2-standard. ESMFold2 incorporates ESMC-6B protein representations and predicts multimolecular structures in a single-sequence configuration ^18,19^. We use query-only inputs to screen newly generated sequences without a separate external multiple-sequence-alignment (MSA) search for each candidate. Computing the language-model representations is included in the prediction cost. The standard model also supports MSA conditioning; query-only evaluation is the configuration adopted here.

Nucleic-acid designs are evaluated with AlphaFold3 (AF3), a broadly adopted reference for modeling interactions among proteins, nucleic acids and small molecules ^6^. Context-chain searches are reused through matching MSA caches, while designed chains remain query-only. This protocol uses AF3’s multimolecular modeling capability while sharing context features across candidates. MotifBench retains its official ESMFold-based evaluation, and ligand generation is assessed through chemical validity, QED and Vina docking.

### 3.2 Sequences and successful backbones

PartiDesign directly generates a sequence alongside its structure (Self). We separately evaluate one sequence redesigned with its interface identities fixed (Self-FI) and ordinary MPNN redesigns, where available. These branches distinguish direct sequence generation from the subsequent designability of a backbone. Self and Self-FI each contribute one sequence; the number of ordinary redesigns is task-specific.

For a case with *n* backbones and *K* ordinary redesigns per backbone, the MPNN@1 sequence passing rate is the number of passing redesigns divided by *nK*. It estimates the success probability of one uniformly selected redesign. MPNN@*K* backbone passing is the fraction of backbones with at least one passing redesign.

### 3.3 A standardized 24-hour compute budget

Computational cost determines how many candidate designs can be generated and validated within a fixed GPU budget. We measured generation, sequence-redesign and structure-prediction latencies on a single RTX 4090-48G GPU, using workloads spanning 50–500 tokens to characterize size-dependent costs (Fig. 6).

**Figure 6.**
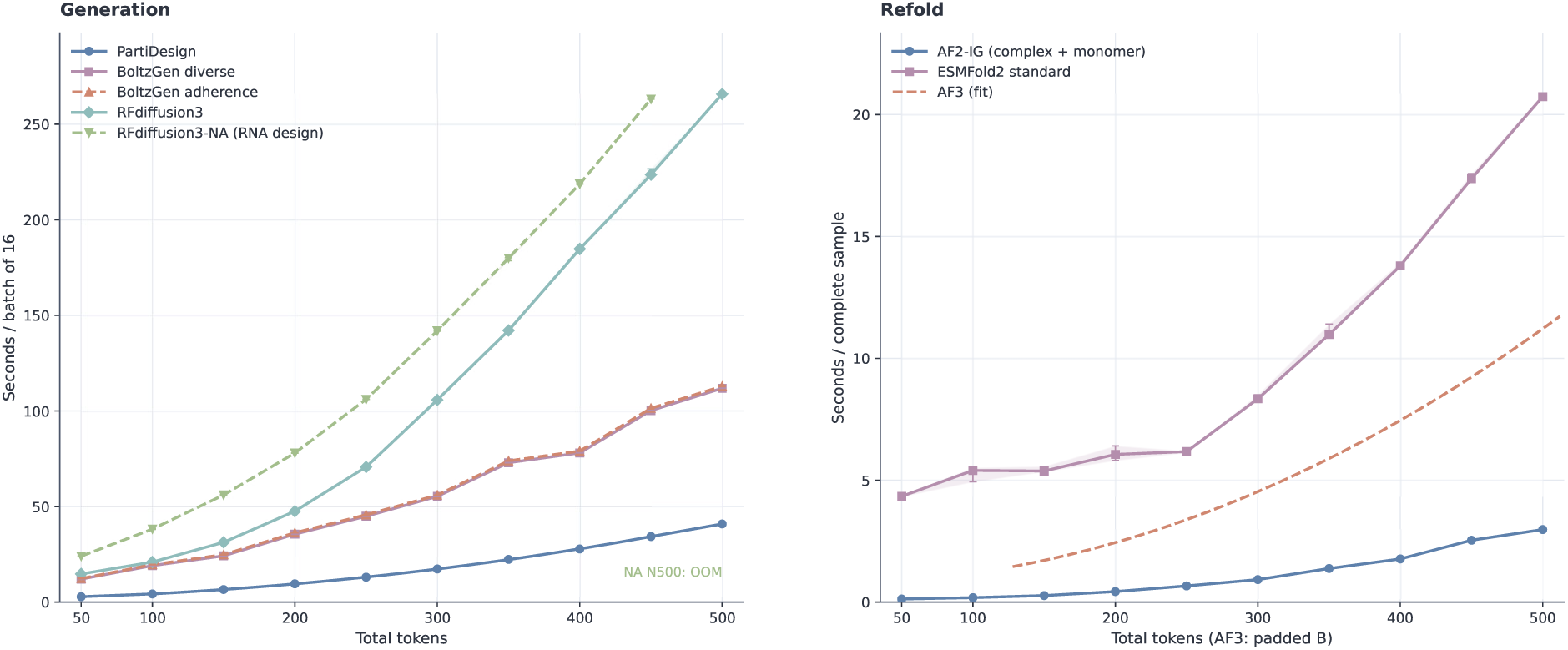
Steady-state compute cost of generation and structural validation. Generation timings use batches of 16, while each validation timing covers one complete sample, including both complex and monomer predictions for AF2-IG. Bands span the 10th–90th percentiles of repeated timings. Runtime is plotted against actual total token count for all methods except AF3, whose curve is a quadratic fit over padded bucket sizes 128–512. RFD3NA runs out of GPU memory at 500 tokens. Section F provides the raw AF3 measurements and the budget mapping.

Generation uses native batches of 16, so the per-candidate generation cost *g* is the batch latency divided by 16. The complete validation cost *v* for AF2-IG includes both complex and isolated designed-chain predictions; the redesign cost *m* measures one sequence designed for one backbone. We combine these costs with each case’s filter-pass rate to estimate successful outputs under a standardized allocation of one GPU for 24 hours <u>per case</u>.

For Self screening, the per-backbone cost is *C*_Self_ = *g* + *v*. Where a two-sequence protocol is available, its cost is

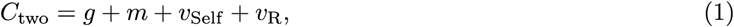

which becomes *g* + *m* + 2*v* when the two full refolds have the same cost. The redesign branch R is determined by the archived experiment: it is an ordinary redesign for most tasks, and Self-FI for nucleic-acid binding to ligands. If *p*_Self_ and *p*_R_ are the marginal passing probabilities and *p_∪_* is the probability that at least one of these two sequences passes, the expected successful backbone and sequence outputs are, respectively,

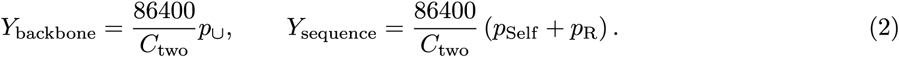

The union probability is estimated from paired outcomes on the same backbone, preserving their empirical dependence. A second sequence can increase backbone recovery while reducing daily output through its additional validation cost. ProtDBench uses separate single-sequence budgets for Self, Self-FI and ordinary MPNN@1; a redesigned sequence costs *g* + *m* + *v*.

One-time CPU setup and model initialization are amortized over the many candidates in a large benchmark and contribute little to its per-candidate cost. We compare synchronized, post-warmup model calls, excluding preprocessing, data transfer and scheduling. Non-AF3 costs use token-length curves; AF3 costs use measured bucket medians. Section F specifies the mappings and shared timing workloads. Per-candidate CPU chemical scoring and docking are reported separately for ligand design.

## 4 Protein design

### 4.1 Protein-binding Protein Binder Design

Protein binder design generates new sequences and structures that recognize specified protein surfaces, supporting diagnostic and therapeutic development. The ten-target ProtDBench panel supplies common target structures and binding-site hotspots and evaluates computational success, structural diversity, and throughput ^20^. Given the target structure and hotspots, PartiDesign generates a new protein chain with its sequence and atomic coordinates. Normal, *α*, and *β* conditions share the targets and sampling allocation; the latter two add coarse secondary-structure guidance.

Evaluation followed the published AF2-IG-Easy screen using AlphaFold2 initial-guess (AF2-IG) predictions of the complex and isolated binder ^20^. A sequence passed only when the same complex–monomer prediction pair met all four conditions: binder pLDDT *>* 0.8 on a 0–1 scale, interface pTM *>* 0.5, interface predicted aligned error (PAE) *<* 10.85 Å, and bound–unbound RMSD *<* 3.5 Å. pLDDT assesses local structural confidence, interface pTM and PAE assess confidence in the binder–target arrangement, and RMSD tests conformational consistency in isolation.

Under the ProtDBench compute budget, normal and *α* PartiDesign both yield more passing candidates from Self than from one ordinary MPNN redesign (Fig. 7). Their estimated ordinary-MPNN yields also exceed those of BoltzGen and RFD3. These estimates combine passing probabilities with measured generation, redesign, and complete AF2-IG costs (Section D.9).

**Figure 7.**
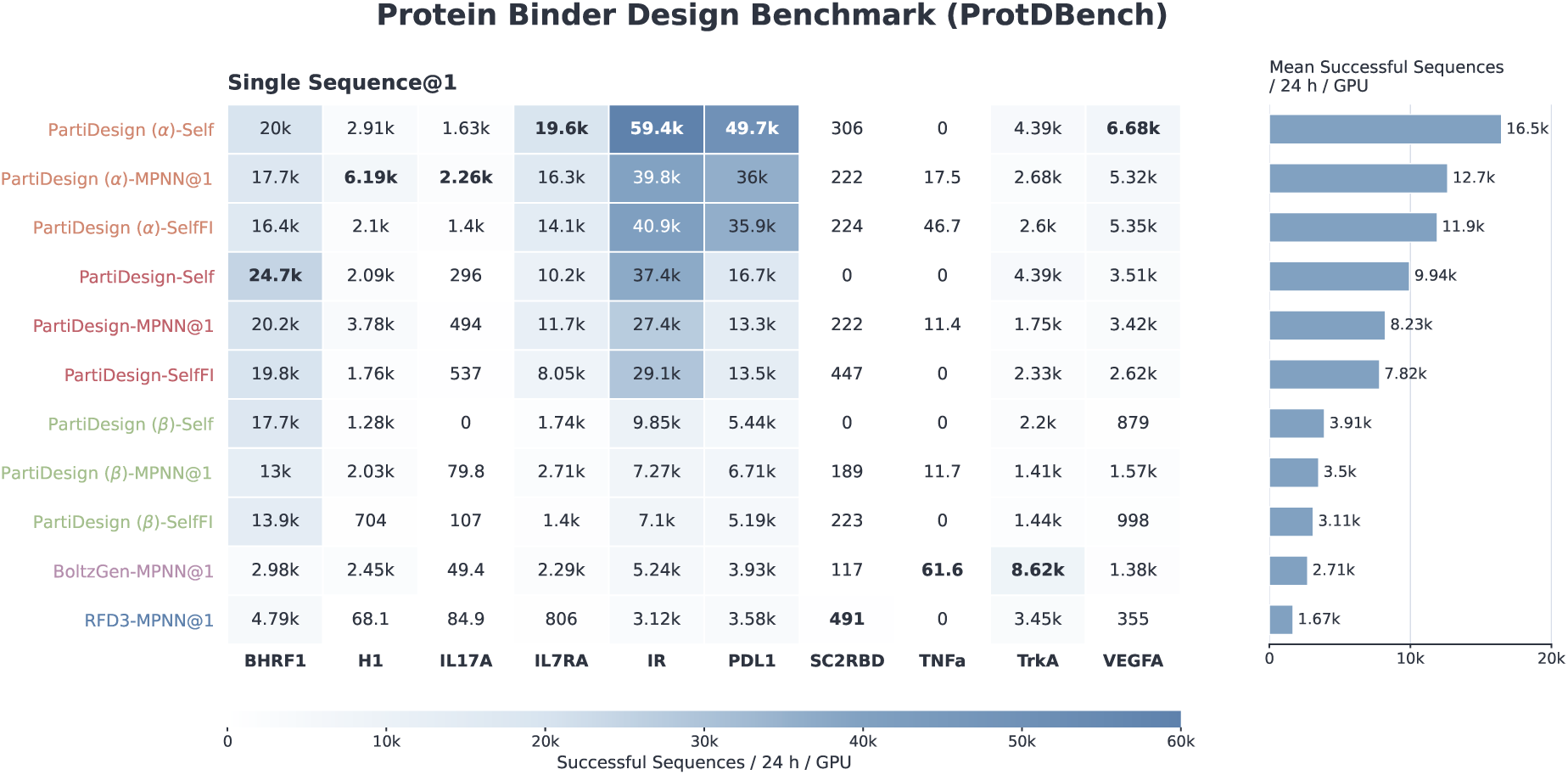
Estimated ProtDBench passing candidates per GPU-day under Single Sequence@1. Normal PartiDesign yields a macro-average 9,938.45 passing Self sequences or 8,225.70 passing ordinary MPNN sequences; *α* yields 16,459.36 and 12,650.12, respectively. The corresponding ordinary MPNN yields are 2,711.22 for BoltzGen and 1,673.84 for RFD3.

PartiDesign *α* has the highest ordinary MPNN@1 passing rate across the full comparison roster, ahead of PXDesign (Fig. 8). Normal PartiDesign also exceeds BoltzGen and RFD3, while the *β* condition falls below normal PartiDesign. For normal PartiDesign, ordinary redesign raises the raw passing rate over Self, but its added compute cost lowers passing-candidate yield.

**Figure 8.**
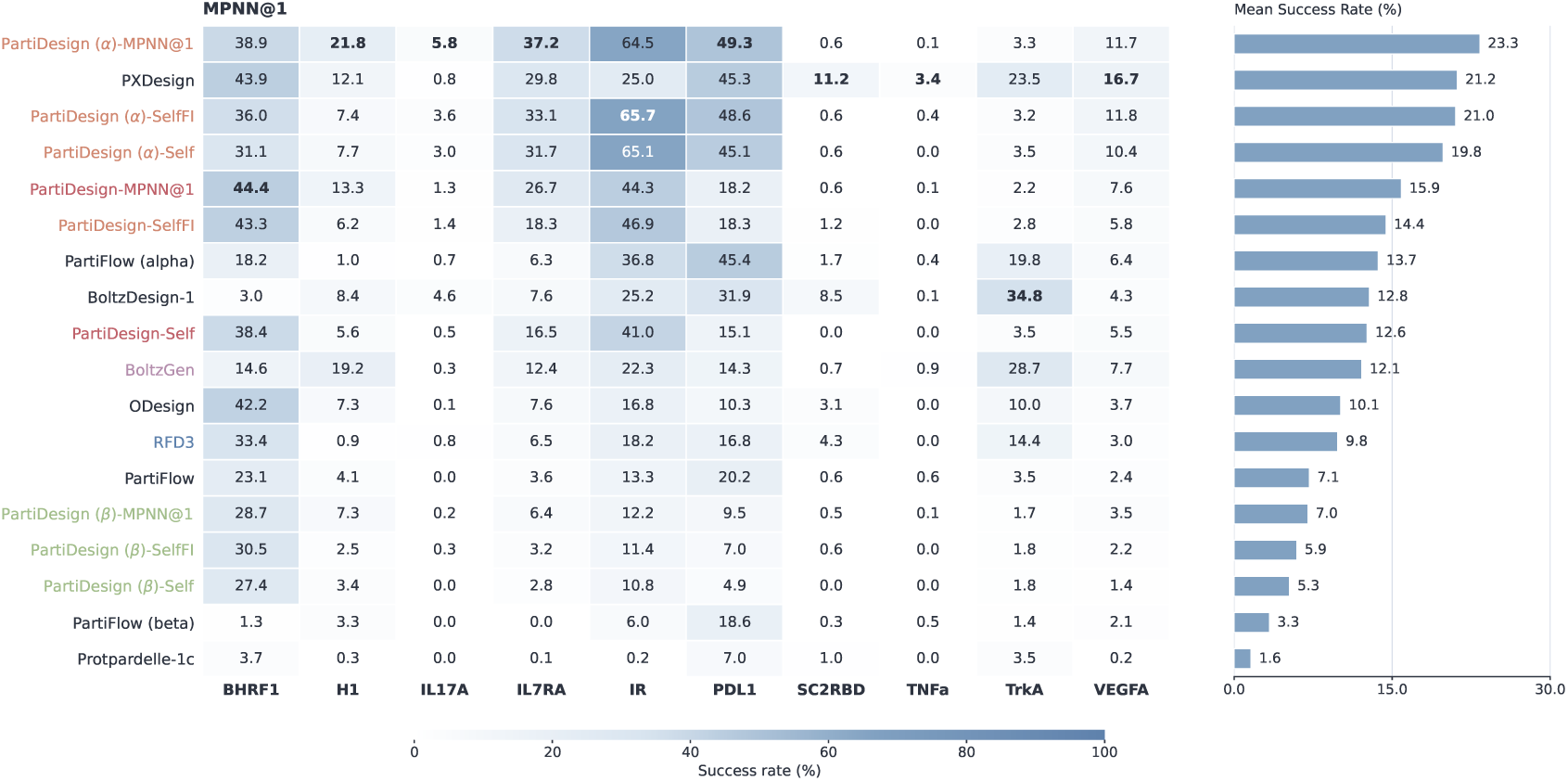
Protein-binder sequence passing on the ten ProtDBench targets under AF2-IG-Easy filtering. Ordinary MPNN@1 macro-average rates are 15.88% for normal PartiDesign, 23.33% for *α*, 7.00% for *β*, 12.10% for BoltzGen, 9.83% for RFD3, and 21.19% for PXDesign. Normal PartiDesign passes 3,416/22,720 ordinary sequences; its Self and Self-FI macro rates are 12.63% and 14.42%, respectively. The full frozen method roster and all three PartiDesign conditions are retained. Side bars give target-equally weighted means; target sample counts differ.

With eight ordinary draws, PXDesign has a higher successful-backbone rate than PartiDesign *α*, reversing their single-draw ordering (Fig. 9). Repeated sequence design therefore changes both the chance of finding a compatible sequence and which backbones count as successful.

**Figure 9.**
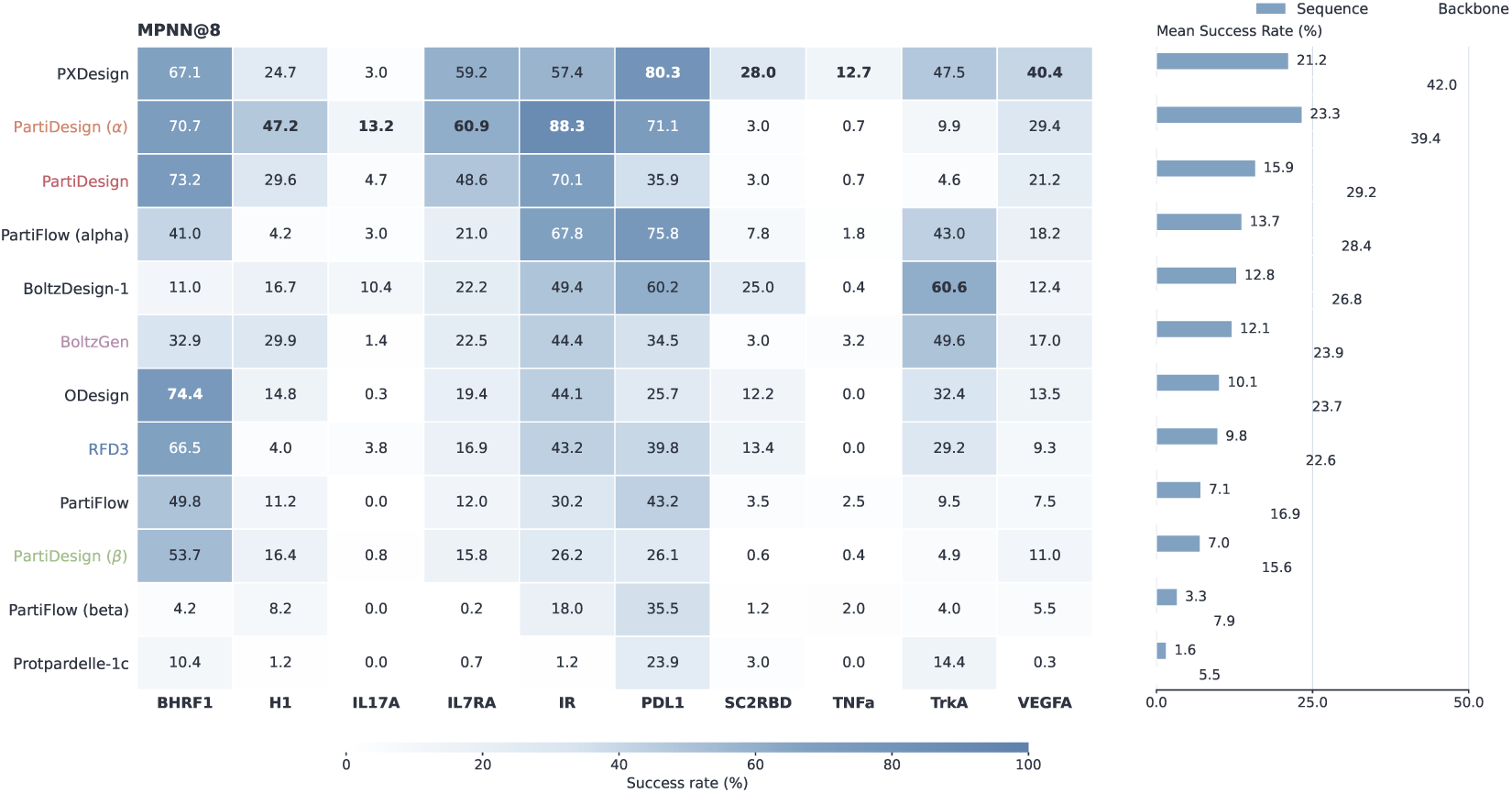
ProtDBench backbone passing with eight ordinary MPNN sequences per backbone. Macro-average successful-backbone rates are 42.03% for PXDesign, 39.45% for PartiDesign *α*, 29.15% for normal PartiDesign, 23.86% for BoltzGen, and 22.62% for RFD3. Self and Self-FI are excluded from this union. PartiFlow uses 400 backbones per target; the other displayed methods use the target-dependent ProtDBench allocation.

The effect of *α* conditioning varies by target. Its ordinary-MPNN successful-backbone count increases from 227 to 286 out of 324 for IR and from 102 to 202 out of 284 for PDL1, but decreases from 120 to 116 out of 164 for BHRF1. Both normal and *α* retain only 5/164 successful backbones for SC2RBD and 2/284 for TNFa. Secondary-structure guidance thus changes the design distribution without improving every recognition problem. Successful-backbone counts also need not track broad structural separation: the ProtDBench-only clustering analysis in Section D.3 reports absolute cluster counts within the ordinary MPNN@8 cohort. PartiDesign structures are clustered with US-align; the published counts were obtained with TM-align and extracted from ProtDBench figures (Figs. S1 and S2).

### 4.2 Protein-binding Peptide Binder Design

Peptide binders are compact molecular recognition scaffolds for diagnostic and therapeutic applications, while covalent cyclization constrains their conformational flexibility ^21^. PepDBench assesses target-conditioned peptide sequence and structure generation across linear, head-to-tail cyclic, and disulfide-linked topologies. Testing all three against the same six protein targets enables comparison under different covalent constraints. Each target and topology has 400 backbones across lengths 8–17 residues.

AF2-IG predictions of peptide–target complexes and isolated peptides were evaluated with an adapted BindCraft screen ^22^, with pLDDT normalized to 0–1. Success required complex peptide pLDDT ≥ 0.8, complex pTM ≥ 0.55, interface pTM ≥ 0.4, interface PAE ≤ 9.3 Å, monomer pLDDT ≥ 0.8, and bound– unbound RMSD ≤ 2.5 Å in the same complex–monomer prediction pair. These criteria assess confidence in the peptide fold and interface arrangement, together with conformational consistency in isolation.

The fixed-compute comparison changes the ordering seen in the six-sequence ALL sequence set, where BoltzGen has higher backbone passing rates for all three topologies (Section D.4). With Self plus one ordinary MPNN sequence, normal PartiDesign has higher estimated successful-backbone yield for linear and cyclic peptides, while BoltzGen retains the higher yield for disulfide peptides (Fig. 10). The budget uses the two-sequence protocol’s own probabilities and costs; the larger ALL union remains a separate fixed-sample comparison.

**Figure 10.**
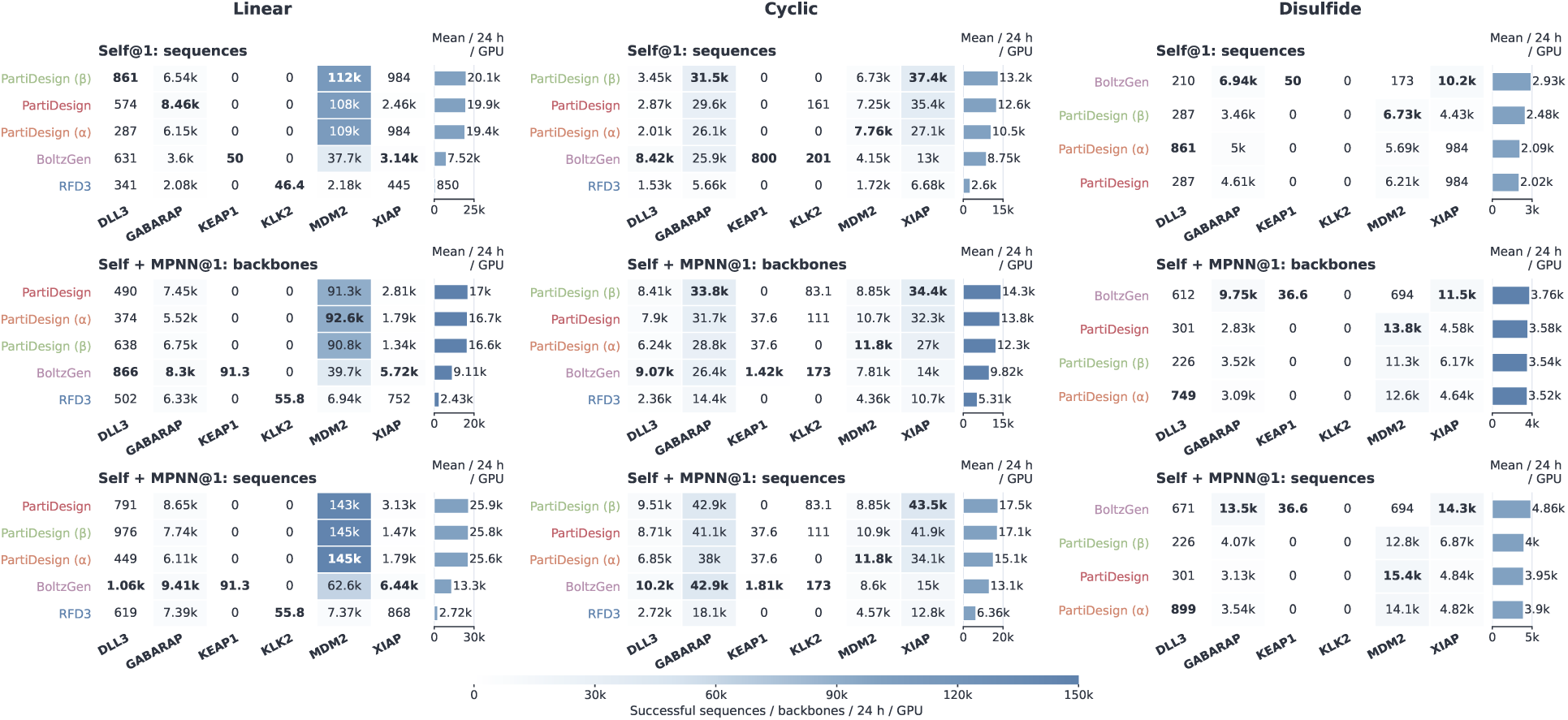
Estimated peptide-binder yield per GPU-day under Self and Self plus one ordinary MPNN sequence. With the latter protocol, normal PartiDesign yields an estimated 17,014.42 linear, 13,785.47 cyclic, and 3,583.01 disulfide successful backbones, compared with 9,110.63, 9,817.18, and 3,761.03 for BoltzGen. RFD3 disulfide remains unavailable; ALL six-sequence passing is not used as its probability.

### 4.3 Scaffolding proteins around functional motifs

Functional motif scaffolding embeds a specified residue geometry in a new protein, supporting the construction of catalytic and molecular recognition sites ^23^. The 30 official MotifBench v1.0 problems constrain motif geometry, residue identities, scaffold length, and motif placement. Their single-segment and multisegment motifs test increasingly complex geometric constraints within a standardized task collection.

Evaluation followed the MotifBench protocol: 100 backbones per problem, eight ProteinMPNN sequences per backbone, and ESMFold refolding ^23,24^. A backbone passed if at least one sequence simultaneously achieved reference-motif RMSD *<* 1 Å and scaffold C*α* self-consistency RMSD *<* 2 Å. These thresholds assess preservation of the functional site and structural consistency of the scaffold. The official MotifBench score accounts for both successful scaffolding and structural diversity ^25^.

PartiDesign’s MotifBench score of 41.86 lies below RFdiffusion3 on the published leaderboard, while successful scaffolds cover nearly all benchmark problems (Fig. 11). The score applies the official 50-of-100 subsampling and saturation rule and differs from the raw passing fraction. The PartiDesign result is a single archived run, whereas RFdiffusion3 reports three runs; these results do not establish a statistically significant difference. The two zero-success problems, 03_2CGA and 30_7UWL, remain in the denominator and score. No measured compute budget is available for this task.

**Figure 11.**
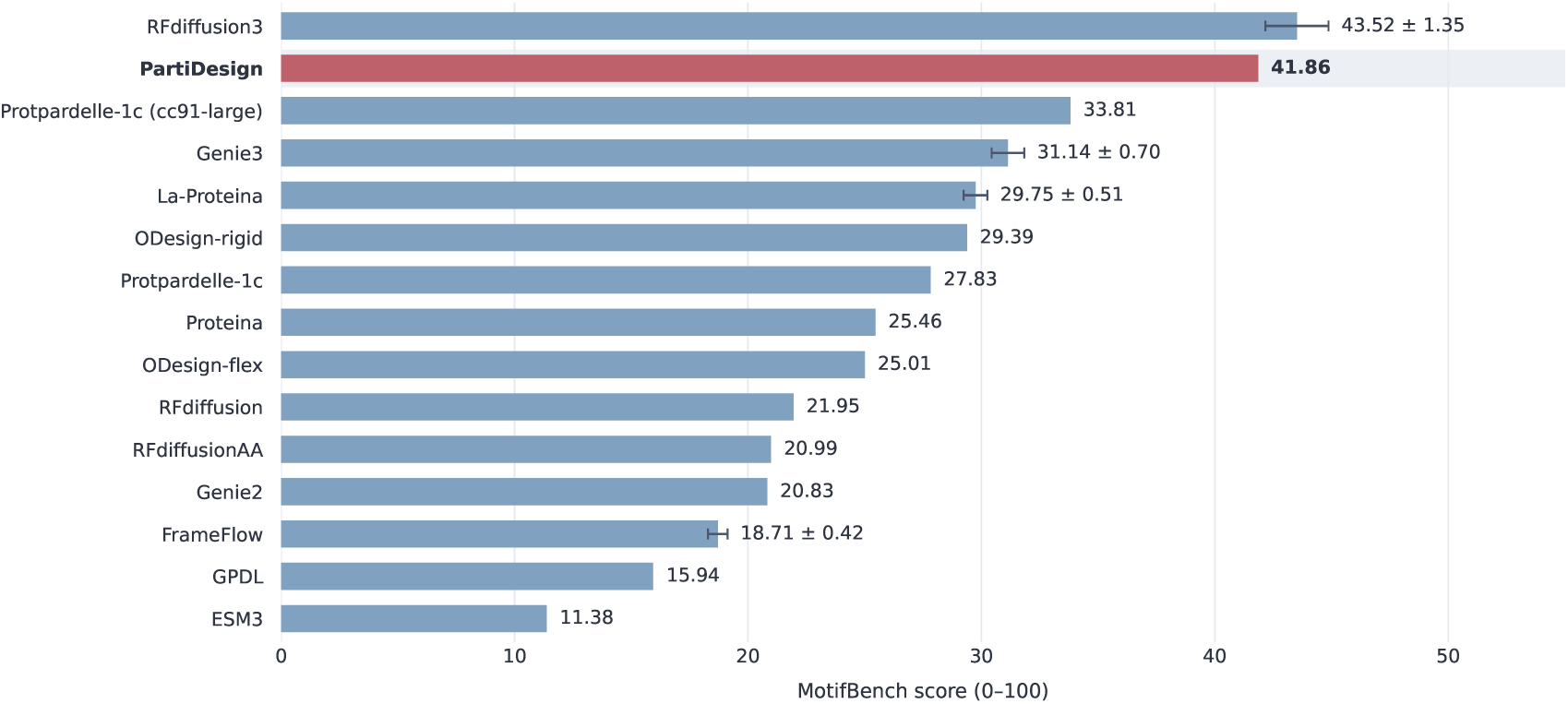
MotifBench comparison using the fixed official leaderboard and the PartiDesign run. PartiDesign obtains 1,078/3,000 passing backbones (35.93%) and solves 28/30 problems. Its score is 41.86, compared with 43.52*±*1.35 for RFdiffusion3. All 30 problems and all 14 public leaderboard entries are included. Published error annotations are preserved as reported and are not relabelled as confidence intervals.

### 4.4 Atomic Motif Enzyme Design Benchmark

De novo enzyme design seeks proteins that support a prescribed arrangement of catalytic functional groups around a substrate or reaction intermediate. Atomic motif scaffolding specifies the geometry of active-site atoms and associated ligands while leaving the surrounding protein scaffold to be designed. The 41-task Atomic Motif Enzyme (AME) benchmark, introduced with RFdiffusion2, supplies motifs comprising one to seven discontinuous segments of catalytic residues, termed residue islands ^26^. This range tests scaffolding under different levels of motif complexity.

We assigned catalytic residues explicit sequence positions before generation. This indexed formulation retains the supplied atomic geometry while fixing where the motif occurs along the chain. Unindexed conditioning allows the model to choose motif placements, but the AlphaProtein Novo authors report slightly better results with indexed conditioning in their experiments ^27^, motivating our indexed adaptation of AME. Normal and *α* PartiDesign used identical motif and ligand inputs to generate 180-residue proteins, differing only in the additional chain-level *α*-helical guide. RFD3 used its default generation settings. Each method or condition generated 100 backbones for each of the 41 cases, with all cases retained.

We applied the active-site preservation criteria used in Complexa with ESMFold2-standard refolding ^14^. Each backbone was evaluated with Self, one LigandMPNN redesign fixing residues at the 6 Å interface, and eight ordinary LigandMPNN redesigns; both redesign branches used temperature 0.1. A sequence passed when one refolded prediction satisfied all four criteria: N/C*α*/C backbone self-consistency RMSD ≤ 2 Å, catalytic motif RMSD ≤ 1.5 Å relative to the supplied reference, recovery of the specified catalytic residue identities, and no protein–ligand atom pair at a distance ≤ 1.5 Å. A backbone passed if any of its ten sequences met the complete criterion.

Both PartiDesign conditions have higher Self and ordinary MPNN@1 passing rates than RFD3 under this indexed ESMFold2 evaluation (Section D.6). Adding one ordinary sequence raises estimated passing-sequence output but reduces successful-backbone yield for both conditions because the extra refolding cost outweighs the increase in backbone passing probability (Fig. 12).

**Figure 12.**
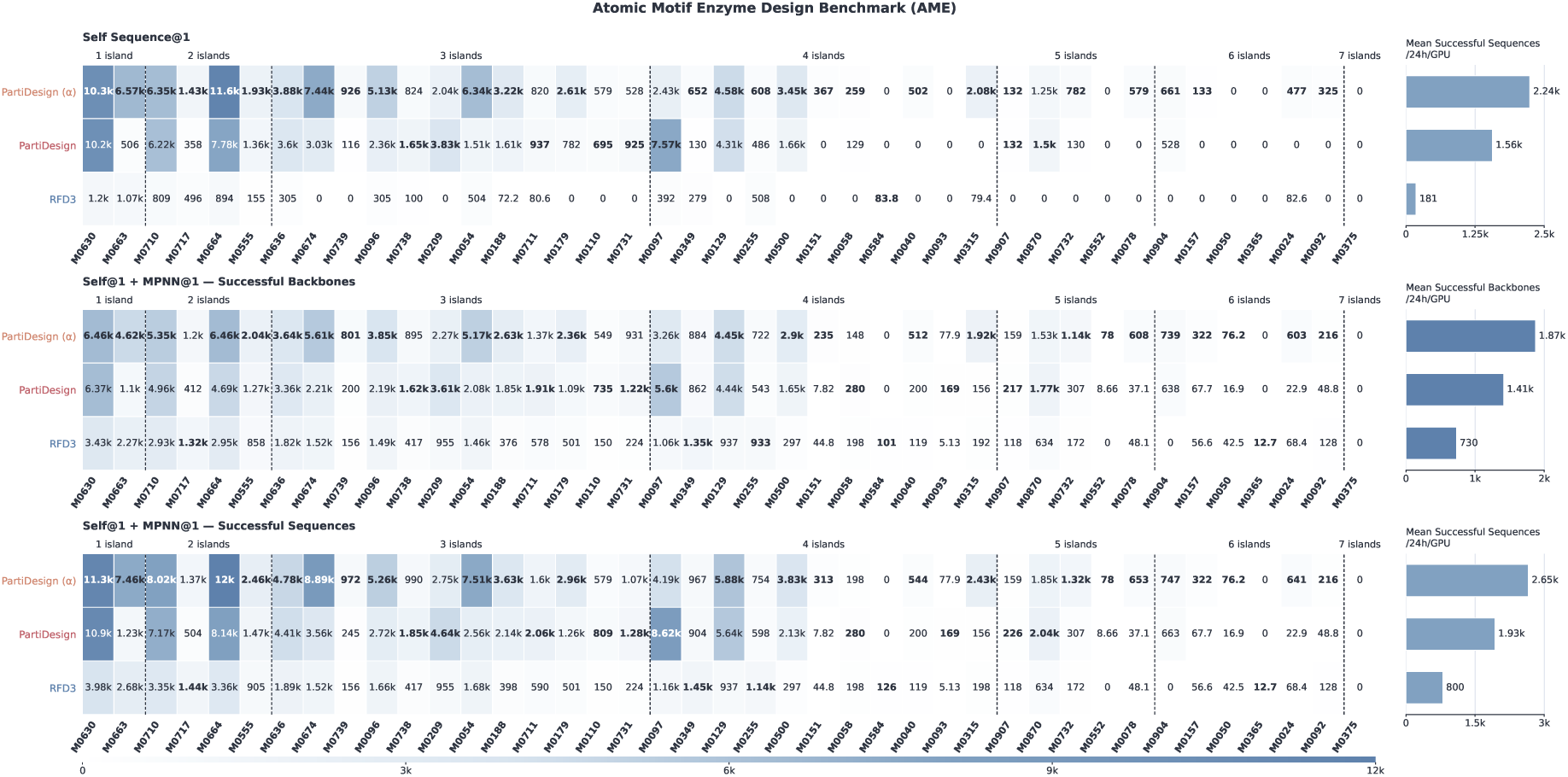
Estimated AME yield per GPU-day under ESMFold2-standard compute costs and the indexed motif protocol. Self yields 1,561.09 and 2,238.41 passing candidates for normal and *α* PartiDesign, respectively. Adding one ordinary MPNN sequence gives successful-backbone yields of 1,412.69 and 1,872.97. All 41 cases are retained, and the ten-sequence ALL union is excluded from this budget.

### 4.5 Nucleic Acid-binding Protein Binder Design

Protein binder design generates a new protein sequence, structure, and placement around a specified DNA or RNA target. Designed DNA-binding proteins support synthetic gene regulation ^28^. Five DISCO-derived targets, comprising three DNA and two RNA complexes, span both nucleic acid chemistries ^29^. The adapted inputs retain complete selected nucleic acid chains and define local interaction regions. Each case has 124 protein backbones across lengths 50–80 residues, with flexible context geometry during generation.

ESMFold2-standard evaluates geometric self-consistency between the predicted and generated complexes. Success requires joint complex RMSD *<* 5 Å after a single rigid-body alignment of protein C*α* and observed nucleic acid phosphate atoms, weighting each atom equally. This project-defined criterion tests whether the protein fold and its placement relative to the nucleic acid reproduce the design.

With Self plus one ordinary LigandMPNN sequence, BoltzGen has higher estimated successful-backbone yield than either PartiDesign condition, while both PartiDesign conditions exceed RFD3 (Fig. 13). These estimates use only the primary ESMFold2-standard records. The target-level results remain heterogeneous: the six-sequence sample set favours normal PartiDesign over BoltzGen on 7EDS and 8BWT, while BoltzGen leads on 6YMC (Section D.7).

**Figure 13.**
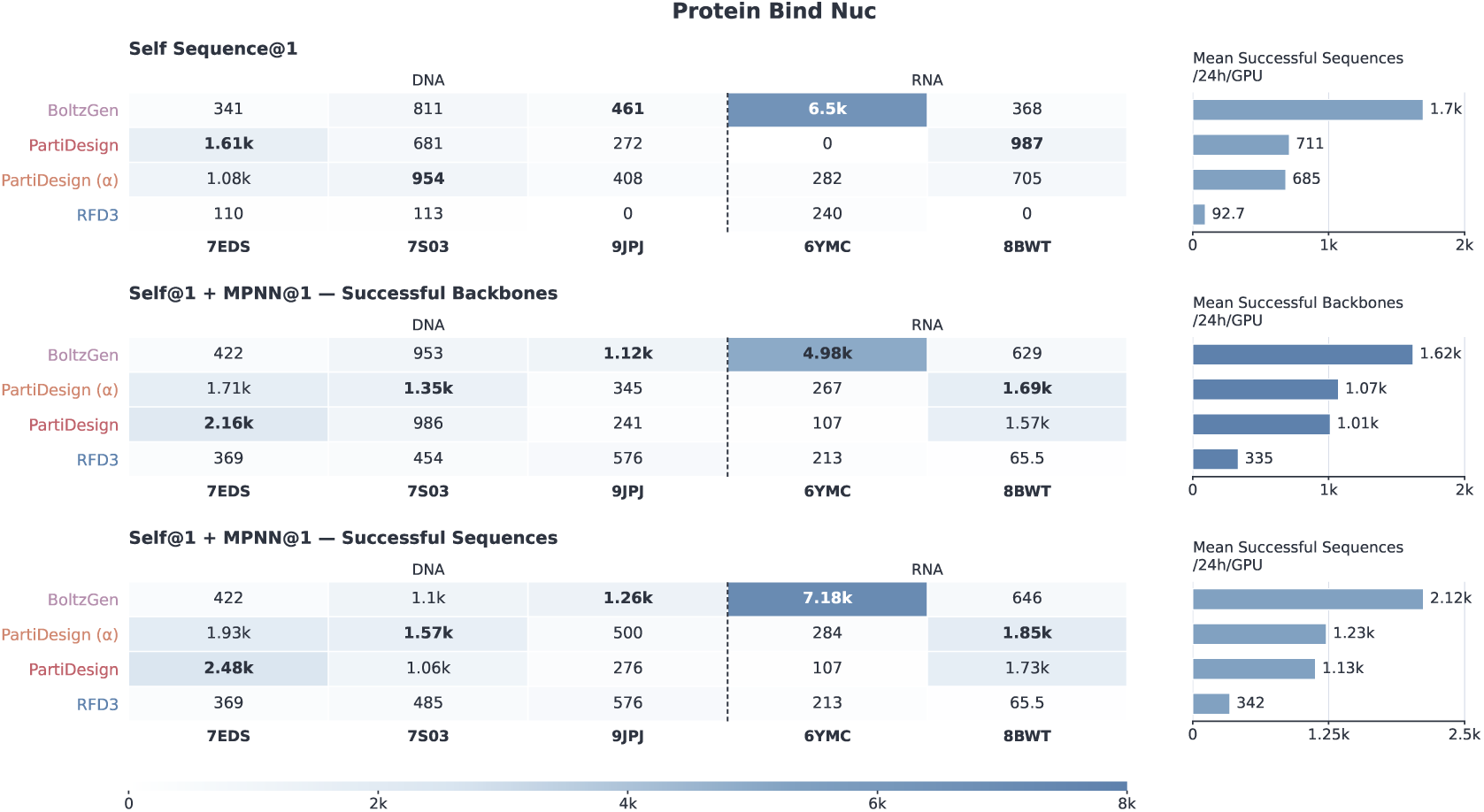
Estimated yield per GPU-day for protein design around DNA and RNA using the primary ESMFold2-standard records. With Self plus one ordinary LigandMPNN sequence, successful-backbone means are 1,011.42 for normal PartiDesign, 1,072.64 for *α*, 1,621.52 for BoltzGen, and 335.45 for RFD3. Secondary AF3 records are excluded.

### 4.6 Ligand-binding Protein Binder Design

Ligand-binding proteins provide recognition elements for molecular sensing and enzyme engineering ^7^. Fixed-ligand design generates a new protein sequence and scaffold around a known small molecule. The four Complexa-derived inputs—IAI, FAD, SAM, and OQO—fix ligand identity and conformation while testing pocket formation across distinct shapes and functional groups ^14^. We removed the original crystallographic protein and generated 200 new 100-residue scaffolds for each ligand.

Evaluation used ESMFold2-standard with thresholds adapted from Complexa’s RF3 ligand-binder screen ^14^. Success required minimum protein–ligand PAE *<* 2 Å, protein C*α* self-consistency RMSD *<* 2 Å, and protein-aligned ligand heavy-atom RMSD *<* 5 Å in one prediction. The PAE statistic is a single minimum over both protein-to-ligand and ligand-to-protein token pairs and identifies a confidently predicted cross-chain relationship. The RMSDs assess preservation of the protein fold and ligand pose between generated and predicted complexes.

Normal PartiDesign’s Self branch passes more sequences than its ordinary LigandMPNN branch, whereas BoltzGen and RFD3 show the opposite ordering (Section D.8). Normal PartiDesign also has higher estimated Self yield than either baseline, and adding one ordinary sequence reduces its successful-backbone yield (Fig. 14). Directly generated identities thus contribute substantially to this panel, with their value varying by ligand and redesign protocol.

**Figure 14.**
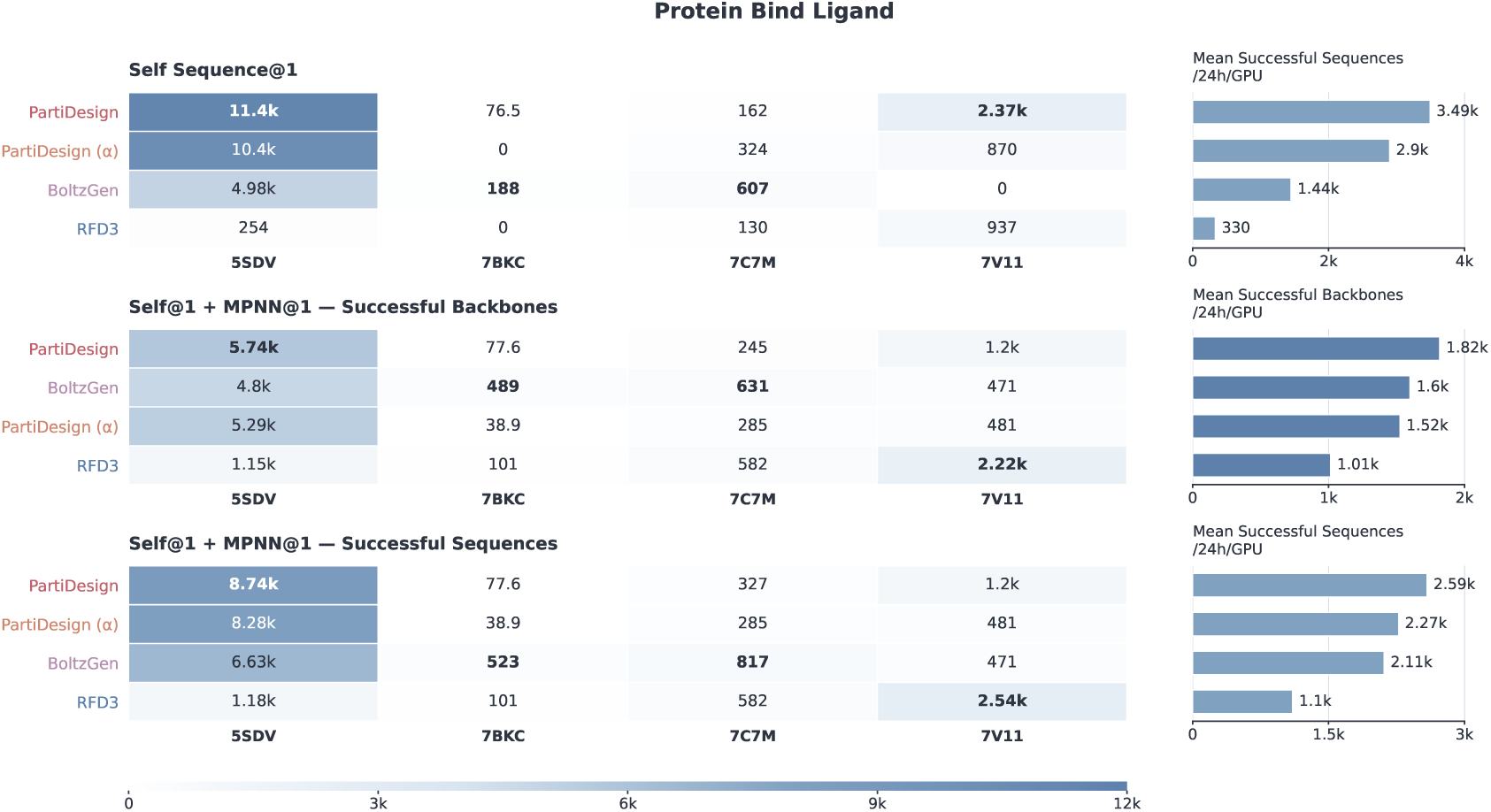
Estimated yield of protein candidates per GPU-day around the four specified ligands, using ESMFold2-standard validation costs. Normal PartiDesign’s Self yield is 3,491.13 passing candidates, compared with 1,443.46 for BoltzGen and 330.28 for RFD3. Adding one ordinary LigandMPNN sequence gives normal PartiDesign a successful-backbone yield of 1,817.04. Self-FI is not added to either protocol.

## 5 Nucleic acid design

PartiDesign generates DNA or RNA identity and structure in the same representation used for protein design. We evaluated two target classes: proteins, which provide extended and chemically heterogeneous interfaces, and small molecules, around which the nucleic acid must organize a recognition environment.

### 5.1 Protein-binding Nucleic Acid Aptamer Design

Protein-binding DNA and RNA aptamers serve as molecular recognition elements for target detection ^7^. Designing them requires coordination of nucleic acid sequence, fold, and placement on a protein surface. We evaluated PartiDesign using NACraft’s NA-12 benchmark inputs ^30,31^. Its six protein–DNA and six protein–RNA targets provide balanced coverage of both nucleic acid chemistries. For each target, the observed protein structure and four interface hotspots condition generation of a new nucleic acid chain at the source binder length; the source nucleic acid sequence and coordinates are removed. DNA lengths span 11–35 nucleotides and RNA lengths 12–71 nucleotides. Some supplied proteins are observed fragments rather than full-length chains. Each method and PartiDesign variant contributes 100 generated backbones per target. The paired variant adds a chain-level preference for paired nucleotides without specifying individual pairing partners.

We evaluated the generated sequence (Self), one fixed-interface NA-MPNN sequence (Self-FI), and four ordinary NA-MPNN sequences per backbone ^32^. AlphaFold 3 (AF3) predicts one complex for each sequence. Our project-defined whole-complex screen requires RMSD *<* 5 Å after a single, equally weighted rigid alignment over the combined protein C*α* and nucleic acid phosphate points of the generated and predicted complexes. The joint alignment assesses aptamer fold consistency and placement relative to the protein in one coordinate frame. Longer proteins contribute more points, and the criterion differs from the directional fits and confidence filters used in the source aptamer studies.

Adding one uniformly selected ordinary NA-MPNN sequence improves fixed-sample coverage and estimated passing-sequence yield for normal PartiDesign, but lowers successful-backbone yield under the compute budget (Fig. 15). The additional sequence incurs a redesign and a second complete AF3 call. These estimates use a cross-route generation proxy and measured fine-64 AF3 padded-shape timings; they exclude CPU feature construction and cache/search work. Normal PartiDesign has a higher raw Self passing rate than RFD3NA, with marked target-specific reversals (Supplementary Fig. S7). Full six-sequence results and protocol details appear in Supplementary Section E.2.

**Figure 15.**
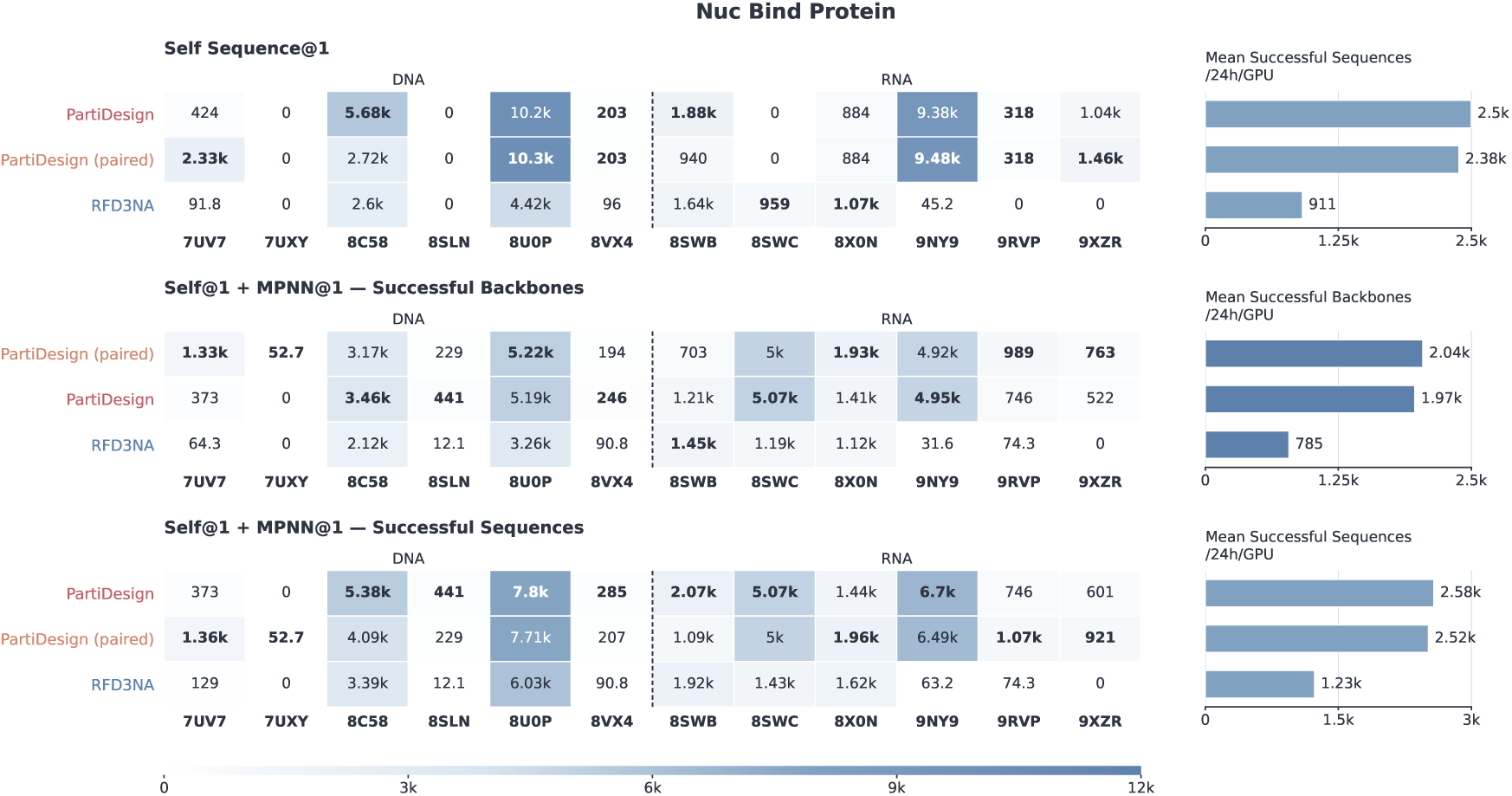
Standardized NA–protein design budgets. Normal PartiDesign Self@1 yields a case-macro estimate of 2,498 passing designs per GPU-day. Adding one ordinary NA-MPNN sequence gives 1,968 passing backbones and 2,576 passing sequences. Raw Self passing rates are 23.67% for normal PartiDesign and 15.75% for RFD3NA. Right-hand summaries are case-macro means. Generation is a cross-route proxy and AF3 cost uses directly measured fine-64 padded-shape medians.

### 5.2 Ligand-binding Nucleic Acid Aptamer Design

Small-molecule-binding aptamers provide recognition elements for biosensing, including electrochemical detection of neonicotinoid insecticides in food samples^33^. Designing such aptamers requires a nucleic acid sequence and fold that accommodate a specified ligand. Our six-case panel uses ethyl maltol, imidacloprid nitroimine, and vanillin as common targets for DNA and RNA design, each with a 20-nucleotide DNA or 30-nucleotide RNA partner. Generation is conditioned on the ligand’s chemical identity and supplied input conformation. These locally selected target–length combinations probe recognition around compact chemical contexts; their lengths were chosen using earlier computational results. Each case contains 100 backbones and two sequence branches, Self and Self-FI. NA-MPNN conditions sequence design on protein and nucleic acid backbone features but does not directly encode ligand atoms ^32^. We therefore omitted ordinary NA-MPNN redesign, which would resample ligand-contacting nucleotide identities without explicit ligand conditioning. Self-FI preserves generated nucleotide identities within 6 Å of the ligand and redesigns the remaining positions.

We refolded the generated complexes with AF3 and applied a project-defined dual-RMSD screen. Passing requires NA-P RMSD *<* 5 Å after fitting the nucleic acid to its generated backbone, and ligand-heavy-atom RMSD *<* 5 Å under that same transform, both in the same prediction. The first condition assesses preservation of the aptamer fold; the second retains errors in ligand placement relative to the aptamer. Ligand RMSD requires chemical identity and complete atom correspondence; unavailable or partially mapped ligand RMSD remains unavailable.

Adding Self-FI raises fixed-sample coverage for normal PartiDesign, yet its redesign and second AF3 call reduce both successful-backbone and passing-sequence yields under the compute budget (Fig. 16). Normal PartiDesign has higher Self and two-branch fixed-sample passing rates than RFD3NA (Supplementary Fig. S8).

**Figure 16.**
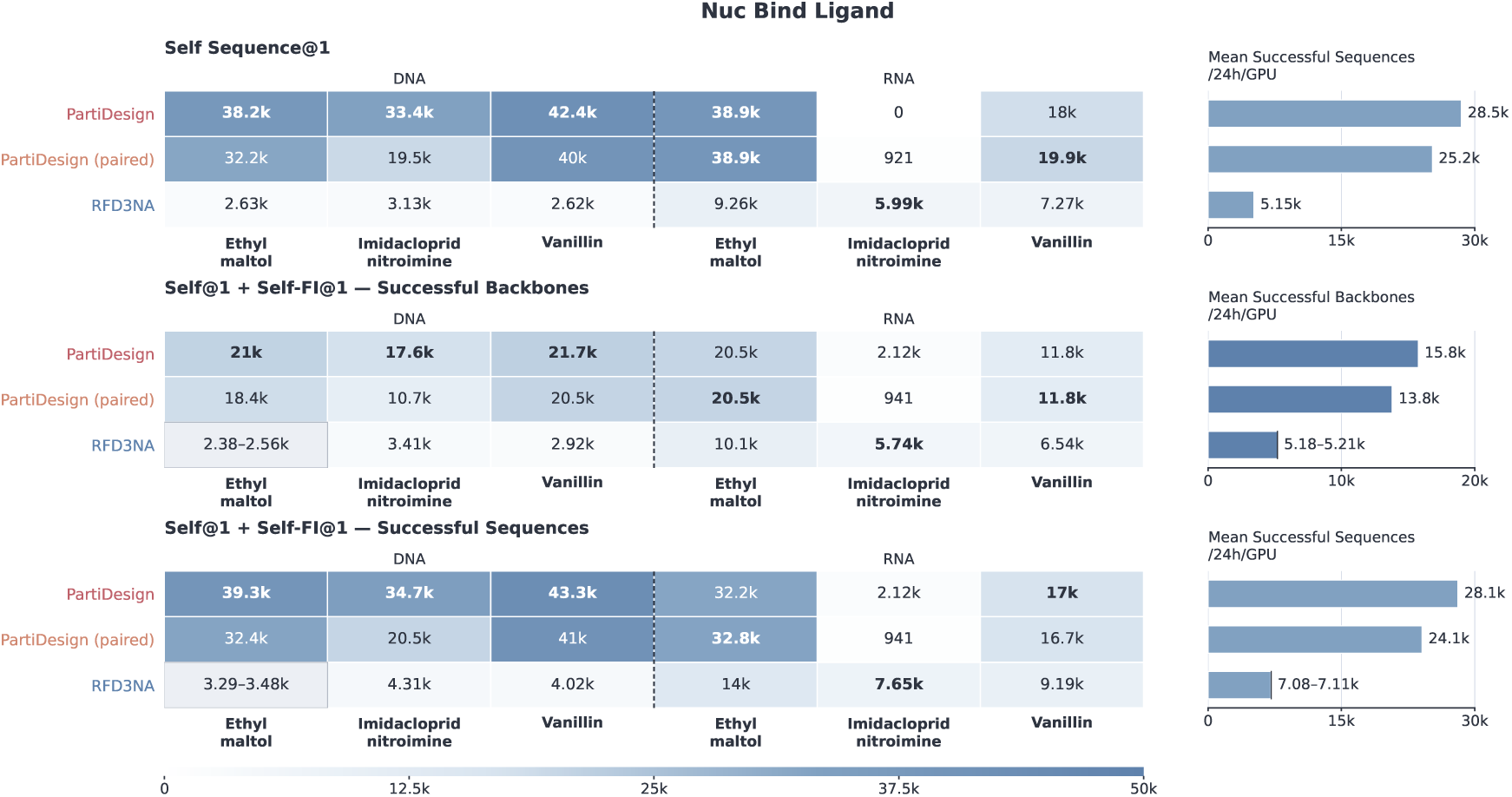
Standardized NA–ligand design budgets. Normal PartiDesign Self@1 yields an estimated case-macro 28,485 passing designs per GPU-day. Self plus Self-FI gives 15,761 passing backbones and 28,104 passing sequences. Passing on the fixed sample set rises from 61.33% for Self to 66.50% for the two-branch union; the corresponding RFD3NA rates are 18.33% and 29.19%. The protocol contains no ordinary NA-MPNN sequence. The RFD3NA DNA–ethyl-maltol missing redesign yields classification bounds, not a failure or a midpoint. The shared generation timing proxy and empty-MSA fine-64 AF3 timing do not measure the evaluation pipeline’s end-to-end throughput.

## 6 Ligand design

Pocket-conditioned molecular generation proposes a ligand’s chemical identity, graph, and three-dimensional conformation within a specified protein binding site for structure-based hit discovery ^7^. This extends conditional generation beyond a polymer alphabet to variable-size chemical graphs. We evaluated five PDB-derived pockets shared with ODesign’s evaluation: 2VT4, 5SDV, 6CM4, 7C7M, and 7V11, with eight explicit hotspot residues and 200 planned attempts per pocket. Heavy-atom count was sampled from the configured size distribution. The protein supplied the structural context; the generated molecule was neither constrained to the native ligand identity nor selected by native-pose RMSD.

Evaluation followed ODesign’s nominal chemical-processing and docking screen ^7^. Validity required RDKit recognition and successful AutoDock Vina docking. Passing additionally required QED *>* 0.25 and a best successful Vina docking score *<* −8.18 kcal/mol for the same postprocessed molecule. QED screens physicochemical drug-likeness ^34^, while Vina provides a model-based estimate of ligand–pocket energetic complementarity ^35^. Success rates use all planned generation attempts as the denominator. More than half of the planned attempts passed the complete screen; the raw pocket-level results appear in Supplementary Fig. S9 and Supplementary Section E.5.

The generation-only budget estimates vary strongly across the five pockets, reflecting pocket dependence in the chemical screen (Fig. 17).

**Figure 17.**
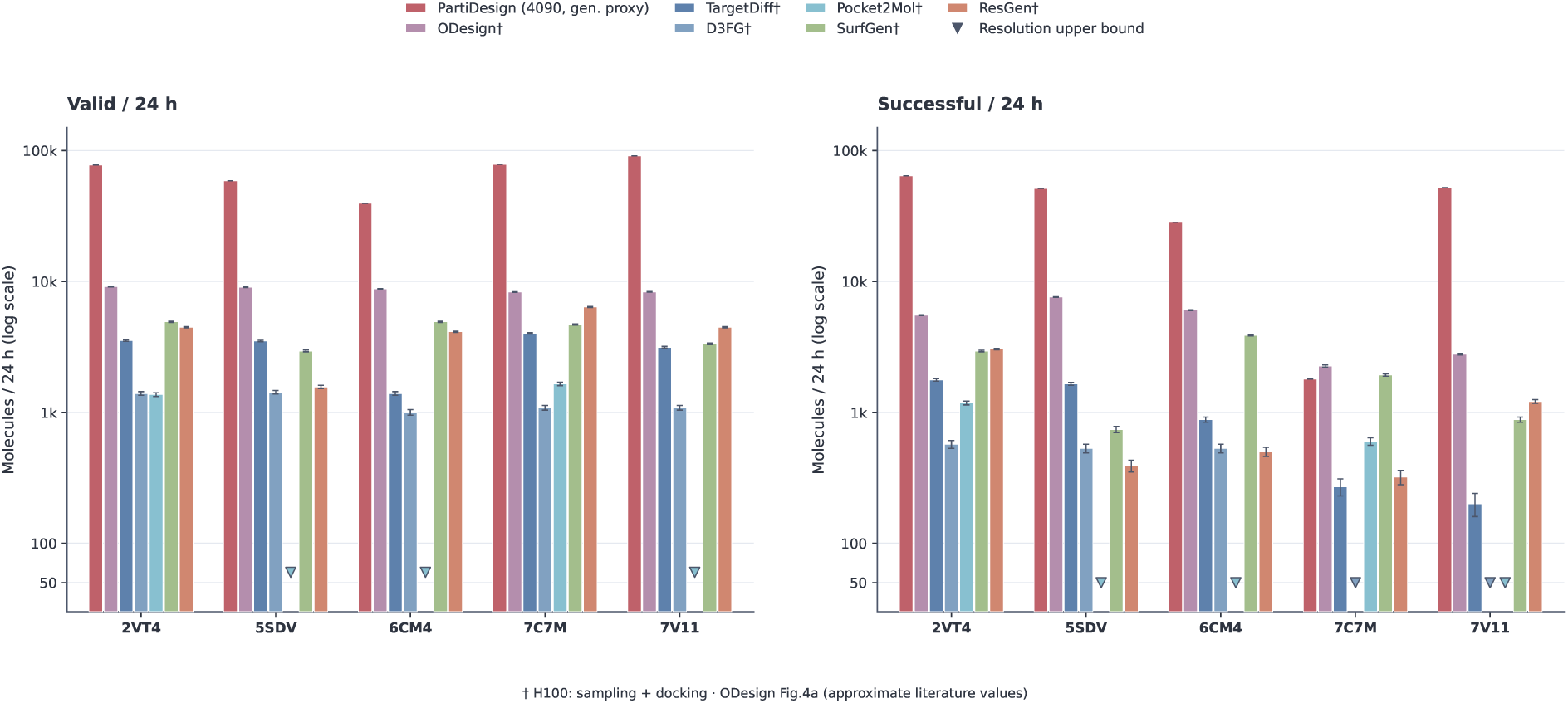
Ligand budget estimates over the five shared pockets. Across 1,000 planned attempts, 945 candidates were valid and 578 passed the complete chemical screen. PartiDesign’s generation-only estimates range from 1,793 to 64,126 passing candidates per GPU-day, using 4090-48G cross-route proxy costs and the complete 200-attempt denominator. Separate CPU chemical scoring averages 101–124 s per planned attempt across pockets and lies outside the GPU budget. Literature series, marked *†*, are raster-digitized aggregates from ODesign Figure 4a for ODesign, TargetDiff, D3FG, Pocket2Mol, SurfGen, and ResGen ^7^; their scope is H100 sampling plus docking. Downward triangles denote positive resolution upper bounds for unresolved small bars. Differing hardware and timing boundaries preclude a normalized speed comparison.

The literature comparison uses the results reported for ODesign and five other methods in ODesign Fig. 4a, which cover sampling and docking on H100 GPUs ^7^. Differences in hardware and timing boundaries, together with incompletely matched pocket preparation, limit this comparison to qualitative trends and preclude a normalized speedup estimate.

## Supplementary Information

### A Training data, representation and optimization

#### A.1 Data sources and sampling

PartiDesign is trained on molecular bioassemblies represented by atom and token arrays. Each training example begins with either a chain or a pair of interacting molecular endpoints. The sampler selects a data source, an eligible chain or interface, and a primary design endpoint through hierarchical sampling with replacement. A single PDB entry can contribute several assemblies and multiple chains or interfaces, so input structures, indexed examples and sampled training examples have different counts. Table S1 lists the source probabilities for the universal VAE and FM mixture.

**Table S1.** Data sources and sampling probabilities for universal VAE/FM training. The PDB subsets share an assembly library. Probabilities describe initial sampling requests; replacement after unsuccessful preparation can alter the distribution of accepted examples.

| Source | Structural content | Probability |
|---|---|---|
| AFDB-v4 | Predicted protein monomers selected using the Genie 2 representative set | 10% |
| Teddymer | Synthetic protein interfaces derived from predicted domain–domain interactions | 20% |
| PDB protein–protein subset | High-confidence protein–protein complexes selected following Complexa | 6% |
| CPSea | Cyclic-peptide complexes with explicit cyclization connections | 4% |
| NuFold-DB | 2026-01-13 rank-1 predicted RNA structure release | 5% |
| PDB nucleic monomer | Nucleic-acid chains from experimental assemblies | 5% |
| PDB protein–nucleic | Protein–nucleic-acid interfaces | 15% |
| PDB nucleic–nucleic | Nucleic-acid–nucleic-acid interfaces | 4% |
| PDB nucleic–ligand | Nucleic-acid–ligand interfaces | 4% |
| PLINDER | Protein–ligand systems from release 2024-04/v1, selected following Complexa | 15% |
| CrossDocked2020 | Positive cross-docked poses from release v1.3 | 12% |

The AFDB component is defined by 588,571 representative identifiers selected by Genie 2, with a maximum sequence length of 256 and a minimum pLDDT of 80 ^36^. The corresponding structures were extracted from the AFDB-v4 representative database using FoldComp ^37^. The count refers to the selected input set before training-time filtering.

Teddymer supplies synthetic binder–target arrangements derived from domain–domain interactions in predicted protein monomers, following the Proteina-Complexa construction ^14^. The input collection contains 519,558 structures and 1,558,096 chain or interface index rows, before training-time filtering.

The experimental assembly library derives from the Protenix-v2 PDB index dated 2026-01-07 ^38^. Entries require a PDB release date before 2025-07-01, a resolution of at most 4.5 Å and at most 20 chains per assembly. The PDB protein–protein subset comprises high-confidence complexes selected following Complexa ^14^. Nucleic-acid chains and the protein–nucleic, nucleic–nucleic and nucleic–ligand interfaces are drawn from the same assembly library.

CPSea contributes cyclic-peptide interfaces whose explicit cyclization connections are retained during structural preprocessing ^39^. Within this source, the universal sampler selects isopeptide, head-to-tail and disulfide categories with probabilities 0.50, 0.25 and 0.25, respectively, and assigns the peptide as the primary design endpoint.

NuFold-DB contributes predicted RNA from its 2026-01-13 rank-1 release, converted into the same assembly representation. RNA structures are generated by the NuFold prediction method ^40^.

PLINDER structures come from release 2024-04/v1 ^41^, using the system selection distributed with Complexa ^14^. Preprocessing uses structural references matched to this release and checks molecular graphs before accepting a converted system.

CrossDocked2020 uses the v1.3 release ^42^. Selection starts from positive poses in the training partition, excludes validation PDB entries and chooses poses deterministically before geometry checks and structural conversion. CrossDocked2020 and the PDB subsets can contain related PDB structures.

RecentPDB and PoseBusters provide VAE/FM validation examples. Training excludes PDB entries present in these validation sets. This exclusion acts at the PDB-entry level and does not implement homology- or ligand-scaffold-based partitioning.

The universal weights and endpoint rules yield primary design-modality probabilities of 55% protein, 23.5% nucleic acid and 21.5% ligand. RNA and DNA share a nucleic sampling class while retaining separate model modalities. Dedicated presets restrict the primary design modality. The protein preset assigns 15, 25, 10, 10, 10 and 30% to AFDB, Teddymer, the PDB protein–protein subset, CPSea, protein–nucleic interfaces and PLINDER, respectively. The nucleic preset assigns 10, 20, 40, 10 and 20% to NuFold, PDB nucleic monomers, protein–nucleic, nucleic–nucleic and nucleic–ligand sources. The ligand preset assigns 70, 10 and 20% to PLINDER, nucleic–ligand and CrossDocked2020 sources. As with the universal mixture, these probabilities apply before replacement of unsuccessfully prepared examples.

### A.2 Cropping and conditional information

The sampling controls below specify hierarchical draws with replacement. The universal mixture allows any primary design modality; the dedicated presets set design_modality to prot, nuc or ligand. Each epoch schedules 10,000 optimizer steps for VAE/FM sampling.

name: universal

strategy: hierarchical_with_replacement

design_modality: null # draw any eligible modality optimizer_steps_per_epoch: 10000

replacement: true # sample with replacement

All three training stages use at most 512 tokens per example. The following excerpt defines the FM crop. Chain examples use contiguous segments; interfaces mix spatial crops with spatial-interface-contiguous crops, which retain contiguous design segments around an interface. If the latter route fails, a spatial crop is attempted for the same source interface and design endpoint.

crop:

crop_size: 512 # maximum tokens per example

route_by_index_type:

chain: contiguous

interface:

spatial_interface: 0.30

spatial_interface_contiguous: 0.70

fallback_by_crop_method:

spatial_interface_contiguous: spatial_interface

preserve_complete_ligand: true # retain complete molecules

max_attempts_per_row: 4

interface_postcheck:

require_both_source_endpoints: true

resolved_heavy_atom_contact_cutoff_nm: 0.5

An accepted interface crop retains both selected endpoints and at least one resolved-heavy-atom contact within 0.5 nm. Interface design segments span 50–256 tokens with 10-token anchor padding. Ligand components remain intact. VAE training uses contiguous chain crops and spatial-interface-contiguous interface crops, without FM design conditions.

After cropping, the primary endpoint identifies the initial design unit. The universal FM recipe permits joint spatial design with probability 0.50 on eligible interfaces; otherwise, other design units contribute full context. The recipe excludes ion endpoints and ligand–ligand interface classes. Conditions are drawn from eligible visible information only after roles have been resolved.

Supported modified polymers are mapped to canonical model slots while retaining their chemical annotations and connectivity. Chain breaks are annotated independently of residue numbering. Coordinates are converted from Å to nm, centered using the available condition geometry, perturbed by a normally distributed translation with standard deviation 0.1 nm and globally rotated. Separate masks identify valid tokens, resolved atoms, design roles, denoising targets and visible conditions. Training does not use MSA or template features.

Motif conditioning preserves selected structural information during generation. The universal recipe draws full-token and selected-atom motifs independently, with probabilities 0.25 and 0.15 for proteins and 0.15 and 0.10 for RNA/DNA. Polymer motifs cover a sampled fraction of 0.05–0.30 in one to four segments. Full-token motifs reveal all resolved atoms in the modality’s canonical slots; selected-atom motifs use a tip/random mixture for proteins and resolved sugar or base atoms for nucleic acids. Ligand motifs are drawn with probability 0.15 and select half of the ligand tokens without replacement.

Additional conditions expose token centers, molecular identities or secondary structure according to the following configuration. Partial identity conditioning reveals 20% of eligible identities. These draws may overlap with motif conditions and therefore do not define mutually exclusive task fractions.

condition:

optional:

token_center_probability: 0.10

token_type_mode_weights: # visible molecular identity

none: 0.70

full: 0.15

partial: 0.15

partial_token_type_fraction: 0.20

polymer_ss:

mode_weights: # fine secondary structure

none: 0.84

token_only: 0.08

token_and_pair: 0.08

polymer_coarse_ss:

probability: 0.08 # whole-design class

Fine secondary-structure conditions label individual tokens and, when selected, their pairwise relationships. Coarse conditions describe the design as a whole. For proteins, the *α* class requires at least 40% *α* and at most 10% *β*, whereas the *β* class requires at least 30% *β* and at most 20% *α*. These assignments require at least ten annotatable residues and 80% annotation coverage. Nucleic-acid labels use reliable base geometry and detected base pairs: the paired class requires at least 30% paired tokens, and the unpaired class at most 1%, with at least four reliable tokens and 80% coverage. Intermediate or insufficiently annotated structures receive the null coarse label.

Hotspot conditioning uses representative-token contacts within 0.8 nm, a sampled retained fraction between zero and 0.2, and dropout probability 0.2. Topology conditioning samples nonstandard edges specified by explicit atom endpoints, excluding normal polymer template bonds. Edges with both endpoints conditioned are exposed. With one endpoint conditioned, visible polymer–ligand edges can be exposed but ligand–ligand edges cannot. With neither endpoint conditioned, polymer–ligand exposure has probability 0.05 and ligand–ligand exposure is zero. The universal recipe does not reveal concrete bond order.

The ligand-specific recipe additionally mixes generation and fixed-graph docking at 0.70/0.30. Its docking branch conditions full ligand identity and graph, including bond order. These training conditions differ from the broader explicit topology constraints accepted at inference.

### A.3 Local variational representation

The local representation pairs each token center with a latent *z_i_* ∈ R^8^. Polymer features encode identity, canonical atomic coordinates, relative/local geometry and torsions; ligand features encode element, coordinate, charge and degree. Pair features describe sequence separation, chain and molecular relationships, center distances, local-frame orientation and bonds. The VAE encoder and decoder each contain 12 Transformer layers with token width 768, 12 attention heads and pair width 256. The encoder outputs a diagonal Gaussian *q_ϕ_*(*z_i_* | *A*) = N (*µ_i_,* diag(exp(2*ℓ_i_*))). Reparameterization samples *z_i_* = *µ_i_* + exp(*ℓ_i_*) ⊙ *ɛ_i_*, with *ɛ_i_* ∼ N (0*, I*)^43^. The training configuration disables latent normalization.

The decoder predicts polymer coordinates and identities from centers, local latents and molecular/chain features. Polymer atom arrays use 37 protein, 27 RNA and 27 DNA slots within a shared 91-slot representation, with masks identifying valid observed atoms. The ligand graph decoder has three message-passing layers, node width 256 and edge width 128. It predicts bond categories, formal-charge classes and element identities. Ligand atoms use their token centers as positions and therefore require no second per-residue coordinate reconstruction.

The optimized objective is

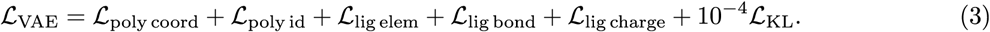

The coordinate term sums squared errors over observed polymer atoms and Cartesian dimensions in nm, then normalizes by the valid polymer-token count per sample. Polymer identity and ligand element terms use masked cross entropy. Ligand bond loss uses unique unordered intra-molecule pairs; bond and charge losses are normalized within each ligand molecule and averaged across molecules. Their configured weights are all one. The KL term sums latent dimensions and averages over valid tokens and samples,

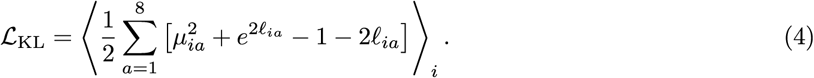

The KL coefficient is constant; annealing is disabled. Reconstruction diagnostics and motif-specific logged losses are not additional optimized terms.

### A.4 Flow matching on centers and local latents

Flow matching learns the joint distribution of token centers and local latents while keeping the trained VAE frozen. The clean latent endpoint is a posterior sample from that encoder. Write *x^c^*_1_ ∈ ℝ^*N* × 3^ for clean centers and *x^z^*_1_ ∈ ℝ^*N* × 8^ for clean latents. For each component *r* ∈ {*c, z*}, an independent standard-normal endpoint *x^r^*_0_ and time *t_r_* define a linear conditional path ^44^:

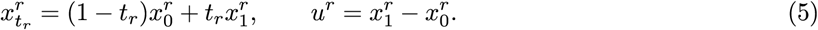

Noise is masked on valid tokens without enforcing zero center of mass. Time is sampled independently per sample and component: *t_c_* ∼ 0.02 *U* (0, 1) + 0.98 Beta(1.9, 1.0) and *t_z_* ∼ 0.20 *U* (0, 1) + 0.80 Beta(1.0, 1.5). Each expression denotes a mixture draw, rather than an arithmetic average of two times.

The 14-layer denoiser has token width 768, 12 attention heads, pair width 256 and condition width 256. Its token inputs contain corrupted centers/latents, self-conditioned clean predictions, visible identities, optional centers, motifs, hotspots and local SS. Independent 256-dimensional time embeddings condition both token and pair processing; coarse SS contributes a 64-dimensional embedding. Pair inputs include relative chain/sequence features, modality and molecular relations, corrupted/self-conditioned/optional center distances, hotspot pairs, visible topology and SS-pair features. Pair representations provide attention biases; iterative pair updates and triangular multiplication are disabled in this configuration.

For the predicted velocity *v^r^*_θ_, the implied clean endpoint is *x̂^r^*_1_ = *x^r^_tr_* + (1 − *t_r_*)*v^r^*_θ_. The implemented loss is

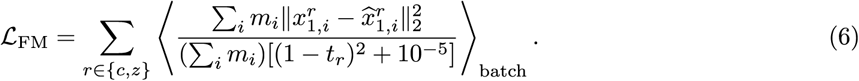

The objective weights clean-endpoint error by time to implement velocity matching with the stated stabilizer. Both product-component weights and both polymer/ligand token weights are one, and the sum over coordinate or latent dimensions is retained. The universal training configuration uses flexible context, so *m_i_* covers all valid tokens. The same implementation also supports rigid context: only denoised tokens contribute, context centers remain fixed and context latents are encoded from context alone using the posterior mean. Rigid clamping and flexible reconstruction thus define different supervision boundaries.

Self-conditioning is selected with probability 0.50 for a training forward. A preliminary pass predicts from the same corrupted state without gradients; its clean prediction is detached and reused in the final forward. Without that pass, the self-condition is zero in flexible mode and the fixed context in rigid mode. Neither pass optimizes the frozen VAE. Motif conditions are indexed to the ordinary token sequence.

### A.5 Optimization and training settings

Table S2 summarizes the optimization settings. Main FM training used 128 Ascend 910B devices with 64 GB each, a batch of eight per device and no gradient accumulation, giving a global batch of 1,024. Training ran for approximately one month. VAE and PartiScore each used 32 Ascend devices, with their respective batch sizes and objectives.

**Table S2.** Training settings. Scheduled bounds specify the maximum training schedule and do not denote completed optimizer updates. VAE training uses the FP32 default of the Ascend training configuration.

| Setting | VAE | Main FM | PartiScore |
|---|---|---|---|
| Devices | 32 Ascend | 128 Ascend 910B, 64 GB | 32 Ascend |
| Batch/device | 20 | 8 | 8 |
| Gradient accumulation | 1 | 1 | 1 |
| Global batch | 640 | 1,024 | 256 |
| Optimizer | AdamW, AMSGrad | Adam | Adam |
| Learning rate | 10<sup>−4</sup> | 10<sup>−4</sup> | 10<sup>−3</sup> |
| Weight decay | 10<sup>−2</sup> | 0 | 10<sup>−8</sup> |
| LR schedule | Constant | Constant | Constant |
| Gradient norm clip | 1 | 1 | 1 |
| EMA decay | 0.999 | 0.999 | 0.999 |
| Token cap | 512 | 512 | 512 |
| Numerics | FP32 | BF16 mixed, eager | FP32 |
| Scheduled bounds | 20 epochs | 100 epochs<br>10,000 steps/epoch | 100 epochs<br>1,000 steps/epoch |
| Reported duration | — | Approximately one month | — |

All stages use fixed learning rates, and the FM optimizer updates only the denoiser parameters. Ascend FM training runs eagerly with HCCL distributed data parallelism. FM and PartiScore use four data-loader workers per device. FM validation and checkpointing occur at epoch boundaries, whereas VAE validation occurs every 5,000 steps. Exponential moving averages are updated after each optimizer step with decay 0.999. A complete-state resume restores the optimizer and training state; initializing from denoiser weights alone begins a new optimizer history. Section B describes the PartiScore training objective and checkpoint metadata.

## B PartiScore mixture density network

### B.1 Structure inputs and network

PartiScore is a mixture density network (MDN) for scoring interfaces across the protein, RNA, DNA and small-molecule modalities supported by PartiDesign. It adapts the distance-likelihood approach of DeepDock and RTMScore, which use MDNs to learn intermolecular distance distributions for protein–ligand pose assessment^45,46^. PartiScore uses the same residue/ligand-heavy-atom tokenization as PartiDesign. Its structure encoder embeds atomic identity, modality, canonical atom slot, element and token-internal radial geometry, pools atom features within each token and projects them to a 256-dimensional node representation. The 91 polymer atom slots are augmented by one slot for a ligand token’s heavy atom. Six node/pair update blocks use eight-head pair-biased attention, residual feed-forward updates and dropout 0.1; pair representations have width 128.

The structure encoder uses 64-dimensional atom embeddings and 16 radial basis channels over intra-token distances. Pair inputs concatenate source-molecule equality, chain equality, ordered modality pair, signed sequence separation, bond category and binned token-center distance. Signed separation is clipped to [−63, 63], with separate categories for different chains and ligand-containing pairs. Center distance uses 30 bins over the configured 0.1–3.0 nm interval and is exposed only within a source molecule. Cross-source center-distance features are zeroed, so the distance between scored partners is evaluated against the predicted density without being supplied directly as that pair’s distance feature. Identities, intramolecular geometry and topology remain available for inferring compatible interactions.

A symmetric readout combines *h_i_* + *h_j_*, |*h_i_* − *h_j_*| and the averaged directed pair representation to predict ten mixture logits, ten means and ten scales. Mixture weights use softmax, means use softplus and scales use softplus plus 0.1 Å. PartiScore takes structures directly and requires no VAE encoding.

### B.2 Training distribution and distance likelihood

PartiScore samples five interface classes: PLINDER protein–ligand (40%), PDB protein–protein (25%), protein–nucleic acid (20%), nucleic acid–nucleic acid (7.5%) and nucleic acid–ligand (7.5%). The PDB interfaces use the assembly library described in Section A. These probabilities apply to initial sampling requests; unsuccessful preparation and replacement can change the distribution of accepted examples. Training uses at most 512 tokens, with spatial-interface and spatial-interface-contiguous crops selected at 70% and 30%, respectively. FM role and condition sampling are disabled, while covalent and source-molecule annotations are retained to define geometric supervision and scoring scope.

For each unique unordered pair of valid tokens belonging to different source molecules, define the target as the minimum distance between observed heavy atoms,

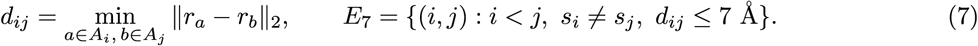

Here *A_i_* contains the token’s valid heavy atoms and *s_i_* denotes source-molecule ownership. Tokens without valid atoms and padded positions are excluded, and distances are converted from canonical nm coordinates to Å. The model represents the conditional distance density as

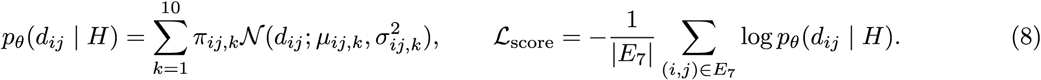

*H* comprises the structure-derived node and pair features. The likelihood is evaluated in the log domain using log-softmax mixture weights and log-sum-exp aggregation. In distributed training, the sum and edge count span the global batch across ranks, so every supervised contact has equal weight. A globally empty edge set skips the update. Training uses distance likelihood without binding-affinity labels, decoy classification labels or a contrastive negative-pose objective. Docking decoys are used only for monitoring.

Optimization uses Adam with learning rate 10*^−^*^3^, betas (0.9, 0.999), weight decay 10*^−^*^8^, gradient-norm clipping at one and EMA decay 0.999. Training used 32 Ascend devices, batch eight per rank, accumulation one and FP32, giving global batch 256. The schedule allowed up to 100 epochs of 1,000 steps, with four data-loader workers per rank. Data preparation excludes validation and benchmark PDB IDs from training sources. This entry-level exclusion does not enforce sequence-homology or chemical-cluster separation.

### B.3 Inference score and molecular scope

For inference, we adapt BioScore’s combination of a contact-normalized distance likelihood and a logarithmic contact-count term to PartiDesign token pairs ^47^. Let *E*_5_ contain the eligible unordered token pairs with *d_ij_* ≤ 5 Å, under the molecular scope defined below. The ranking score is

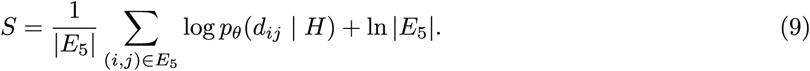

Higher scores rank more favorably: the first term measures the mean log likelihood of the observed contact distances, and the second rewards contact support. The score is not calibrated to binding free energy or experimental success probability, and its dependence on contact count limits comparisons across unrelated tasks.

All-interface scoring evaluates eligible pairs from different source molecules. For design search, design units connected by covalent bonds within the designed region form one scoring group; connections through context do not merge separate designs. The contact set includes pairs between distinct design groups and between design and context. It excludes context–context contacts, contacts within a design covalent group and directly covalently bonded token pairs. Covalent bonds remain network inputs even when their token pairs are excluded from score aggregation. One score is computed over the union of eligible contacts.

Scoring returns no value when no eligible contact or interface is present. During design search, a candidate with no contacts is assigned −∞ for ranking, whereas an inapplicable interface is treated as a preparation failure. Search rewards, final scoring and optional relaxation are enabled independently. When final scoring follows relaxation, the score is recomputed from the relaxed coordinates.

### B.4 Model weights and validation

The released structure checkpoint was saved at epoch 18 and global step 9,500. The export does not specify whether raw or EMA weights were selected.

Training monitoring includes MDN NLL on held-out structures, the 285-complex CASF-2016 docking suite ^48^ and the 79-target BioScore protein–protein docking suite ^47^. CASF monitoring measures Top-1/2/3 recovery of poses with RMSD ≤ 2 Å, both including and excluding the crystal pose. BioScore monitoring uses the supplied DockQ labels and the authors’ success and hit definitions at ranks 1, 5 and 10, together with Spearman and Kendall correlations over all poses and the twenty poses with highest DockQ. CrossDocked monitoring is disabled. The enabled suites are used repeatedly during training and are therefore monitoring sets. The docking analyses in Section G report pose recovery under their respective evaluation protocols.

## C Shared evaluation and statistical definitions

### C.1 Primary structural predictors and sequence branches

Each task uses the predictor in Table S3 to assess its designated design region or interface, with task-specific contexts, fixed motifs and chemical identities. Sections D and E specify thresholds, alignment atoms and sequence counts.

**Table S3.** Primary evaluators and the structural or chemical properties assessed in each task.

| Design task | Primary evaluator | Main evaluation target |
|---|---|---|
| ProtDBench | AF2-IG | Interface confidence and bound/unbound structural agreement |
| PepDBench | AF2-IG peptide | Complex confidence and peptide conformation |
| AME | ESMFold2-standard | Scaffold consistency, atomic motif and ligand accommodation |
| Protein binding ligand or nucleic acid | ESMFold2-standard | Designed protein, interface and relative geometry |
| Nucleic acid binding protein or ligand | AF3 | Nucleic-acid structure and complex arrangement |
| MotifBench | Official ESMFold protocol | Motif scaffolding and official benchmark score |
| Ligand design | RDKit/QED/Vina | Molecule validity, drug-likeness and docking score |

Self evaluates the generated sequence; Self-FI redesigns eligible positions while fixing interface identities. Ordinary MPNN preserves task constraints without fixing additional interface identities. All branches share their originating backbone, so ten sequences can represent one structural sample. RFD3 cyclic peptides retain cyclic topology and use CyclicMPNN.

ESMFold2-standard uses ESMC-6B representations with query-only inputs, although its architecture supports MSA conditioning. AF3 reuses matching context MSA caches and keeps designed chains query-only. These inputs differ from the empty-MSA AF3 timing fixture, which excludes retrieval and preprocessing.

### C.2 Threshold application and missing outcomes

Criteria are applied to unrounded values in their declared units, retaining each strict or inclusive boundary. Confidence scales and PAE units are matched to the task-specific thresholds. For the ESMFold2 protein–ligand filter, the bidirectional minimum is

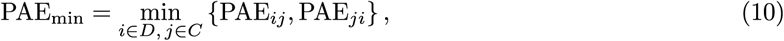

where *D* and *C* are the valid design and target token sets. This statistic selects the lowest individual token-pair error across the two directional interface blocks. All required criteria are evaluated together within one complete prediction. When several predictions are available, at least one complete prediction must satisfy the full set.

Unavailable metrics, incomplete refolds and unresolved identity mappings are distinct from evaluated failures. Planned-attempt denominators can include generation failures with unavailable metrics, as in ligand design. Untested methods and cases remain missing; classification completeness determines point estimates or bounds. Sequences are not deduplicated or filtered for cysteine content.

### C.3 Sequence, backbone and case aggregation

For case *c*, let *I_ik_* denote passage of redesign *k* on backbone *i*, among *n_c_* backbones and *K_c_* available ordinary redesigns. Then

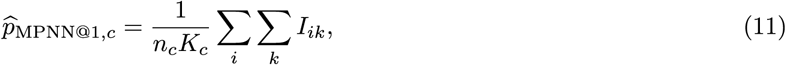

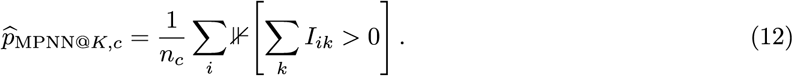

The first quantity estimates single-draw sequence passing; the second measures any-of-*K* backbone passing. Self and Self-FI each use one sequence. ALL joins the task-specific branches, reporting sequence and backbone rates separately. Variants, evaluators and topologies remain separate unless pooling is specified.

The case mean, *C^−1^* Σ_*c*_*p̂_c_*, weights cases equally and can differ from pooled ratios when sample counts vary. Daily yields likewise allocate 24 hours per case before averaging. Published aggregates keep their original granularity; figure-digitized results are identified separately from directly calculated results. Heatmaps summarize benchmark draws, while between-run uncertainty and significance tests require independent replicates.

## D Protein benchmark details

### D.1 Structural predictors and sequence branches

ProtDBench and PepDBench use AF2-IG complex and design-chain monomer predictions; MotifBench uses its official ESMFold workflow. AME and protein design around nucleic acids or small molecules use ESMFold2-standard. Additional AF3 predictions are evaluated separately.

Self is the generated sequence. Self-FI redesigns one sequence while preserving identities at design positions within 6 Å of the context. Ordinary MPNN redesign leaves these interface identities free while protecting required motif identities and task-specific fixed positions. ProteinMPNN handles ordinary protein and linear/disulfide-peptide redesign ^49^; CyclicMPNN handles head-to-tail cyclic peptides. AME and the two nonprotein-context tasks use LigandMPNN ^50^. Temperatures are 0.0001 for ProtDBench and PepDBench and 0.1 for AME and the nucleic-acid and small-molecule tasks. The number of ordinary draws is eight for ProtDBench and AME, four for PepDBench and nucleic-acid contexts, and one for small-molecule contexts.

MPNN@1 estimates single-draw passing probability from the fraction of passing ordinary sequences. ALL (Union in the nucleic-acid panel) joins Self, Self-FI, and ordinary draws: ten sequences in AME, six in PepDBench and nucleic-acid contexts, and three in small-molecule contexts. A backbone passes if one complete prediction satisfies the entire criterion. Missing or nonfinite required values and incomplete predictions are unavailable; without an observed pass, an unresolved draw leaves backbone status unresolved. All primary protein results are classifiable. Sequences are not deduplicated or filtered for cysteine content.

Figure side bars weight cases equally, so macro-averages can differ from pooled fractions, as in ProtDBench. Applying the criteria reproduces all 719,400 sequence classifications across the five non-MotifBench tasks.

### D.2 ProtDBench allocation, filtering, and comparisons

The ten targets are BHRF1, H1, IL17A, IL7RA, IR, PDL1, SC2RBD, TNFa, TrkA, and VEGFA. Four backbones at each allowed integer length give 2,840 PartiDesign backbones per condition (Table S4). Each of the six public ordinary-MPNN sets also contains 2,840 backbones; PartiFlow uses 400 per target and 4,000 per condition. Comparisons include PXDesign, BoltzDesign-1, ODesign, Protpardelle-1c, all three PartiFlow conditions, PartiDesign, BoltzGen, and RFD3 (Table S5).

AF2-IG uses model 1, one sample, and three recycles for separate complex and monomer predictions. All four AF2-IG-Easy inequalities must hold for one sequence: binder pLDDT *>* 0.8, interface pTM *>* 0.5, normalized interface PAE *<* 0.35, and bound–unbound RMSD *<* 3.5 Å. PAE normalization divides by 31, making its threshold equivalent to 10.85 Å. The ProtDBench criterion adds no pTM or energetic filter ^20^. Bound–unbound RMSD values are retained as evaluated, but their atom selection is incompletely specified: the evaluation implementation specifies paired C*α* atoms, whereas the budget protocol specifies backbone triplets.

**Table S4.** ProtDBench allocation and ordinary MPNN@8 successful backbones for PartiDesign. Each entry counts physical backbones, excluding Self and Self-FI.

| Target | Backbones | Normal | *α* | *β* |
|---|---|---|---|---|
| BHRF1 | 164 | 120 | 116 | 88 |
| H1 | 324 | 96 | 153 | 53 |
| IL17A | 364 | 17 | 48 | 3 |
| IL7RA | 284 | 138 | 173 | 45 |
| IR | 324 | 227 | 286 | 85 |
| PDL1 | 284 | 102 | 202 | 74 |
| SC2RBD | 164 | 5 | 5 | 1 |
| TNFa | 284 | 2 | 2 | 1 |
| TrkA | 284 | 13 | 28 | 14 |
| VEGFA | 364 | 77 | 107 | 40 |

**Table S5.** Ten-target macro-average ordinary MPNN passing rates in ProtDBench. MPNN@1 is a sequence rate and MPNN@8 a backbone rate; pooled counts are not substituted for these means.

| Method or condition | MPNN@1 (%) | MPNN@8 (%) |
|---|---|---|
| PartiDesign α | 23.33 | 39.45 |
| PXDesign | 21.19 | 42.03 |
| PartiDesign normal | 15.88 | 29.15 |
| PartiFlow α | 13.68 | 28.35 |
| BoltzDesign-1 | 12.84 | 26.81 |
| BoltzGen | 12.10 | 23.86 |
| ODesign | 10.11 | 23.67 |
| RFD3 | 9.83 | 22.62 |
| PartiFlow normal | 7.14 | 16.95 |
| PartiDesign β | 7.00 | 15.59 |
| PartiFlow β | 3.35 | 7.90 |
| Protpardelle-1c | 1.60 | 5.52 |

### D.3 ProtDBench successful-structure clustering

Clustering includes backbones with at least one passing ordinary MPNN sequence among eight draws, excluding Self and Self-FI. PartiDesign uses design-chain C*α* coordinates and all pairwise US-align alignments^51^, averaging each pair’s two length-normalized TM-scores. Five-decimal scores are compared inclusively at TM ≥ 0.4, 0.6, 0.8, 1.0, without added tolerance. Greedy clustering selects the structure with the most unassigned neighbours, breaks ties by stable sample identifiers, and removes the centre and its neighbours. Each cluster contributes once to the absolute case-level count *K*; neither *K/N* nor *K/S* is reported.

Successful-backbone counts and structural separation can change differently (Fig. S1). The 120 successful normal-PartiDesign BHRF1 backbones form *K* = 8, 63, 109, 120 clusters at TM thresholds 0.4, 0.6, 0.8, and 1.0. On H1, *α* increases successful backbones from 96 to 153 while the TM0.4 cluster count falls from seven to six. More passing backbones therefore need not represent more broadly separated structures.

The published BoltzGen and RFD3 counts in Fig. S1 retain the original TM-align/greedy protocol. Daggered values are digitized reconstructions; public TM0.4 values were unreported and remain unavailable. Different alignment engines and the absence of compute normalization limit comparisons of small cross-method differences in absolute *K*.

**Figure S1.**
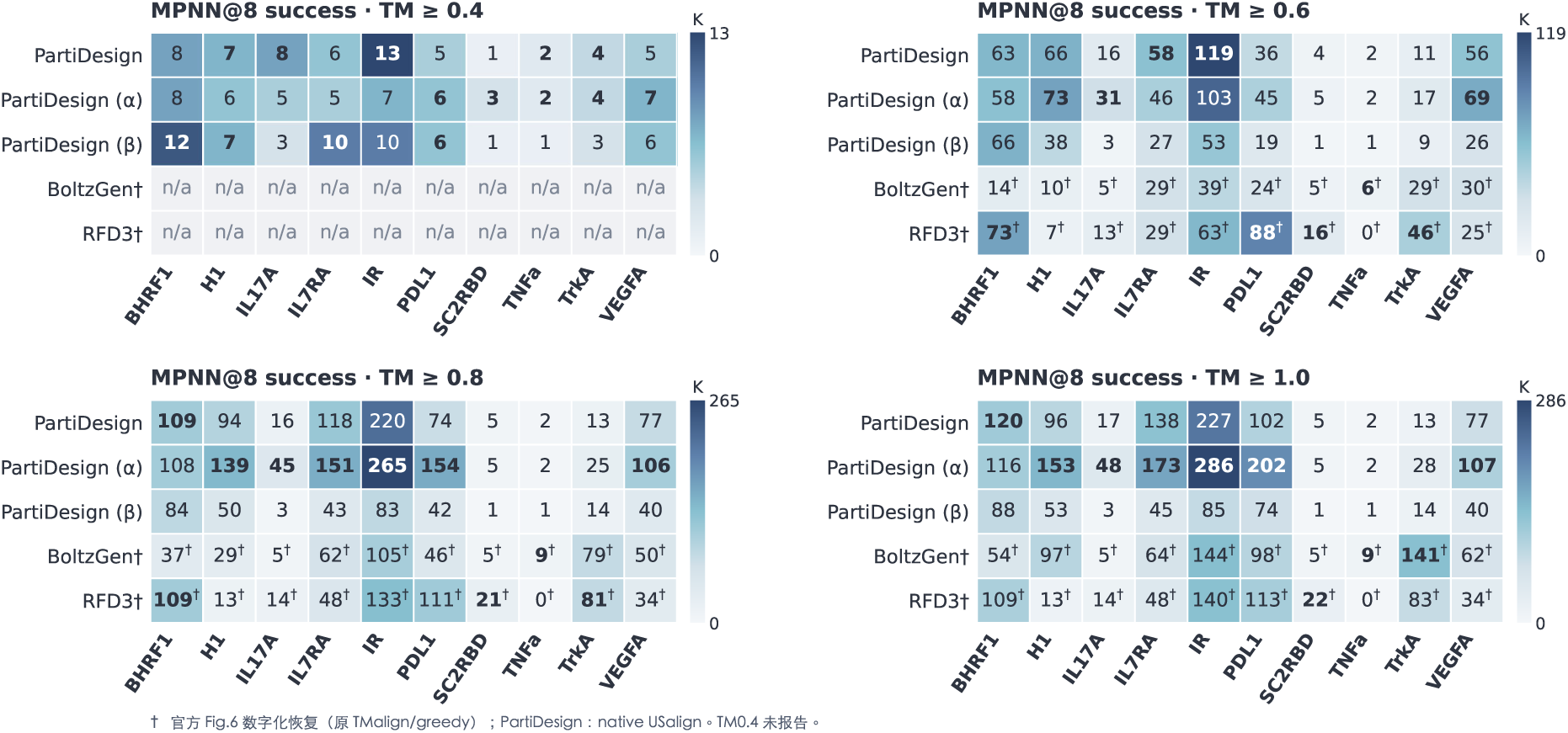
Absolute cluster counts for successful ordinary MPNN@8 backbones per ProtDBench target. Four TM thresholds are shown for PartiDesign US-align clusters. Daggered BoltzGen and RFD3 counts are digitized from the published TM-align figure; unreported TM0.4 entries remain unavailable. Self and Self-FI are excluded.

The expanded comparison includes PXDesign, BoltzDesign-1, ODesign, and Protpardelle-1c (Fig. S2). All 180 public counts were digitized from ProtDBench Figure 6/A.5 using bar-boundary intervals and official sample allocations. Each interval admits one integer count; these are figure-derived values rather than an author-released exact count table.

**Figure S2.**
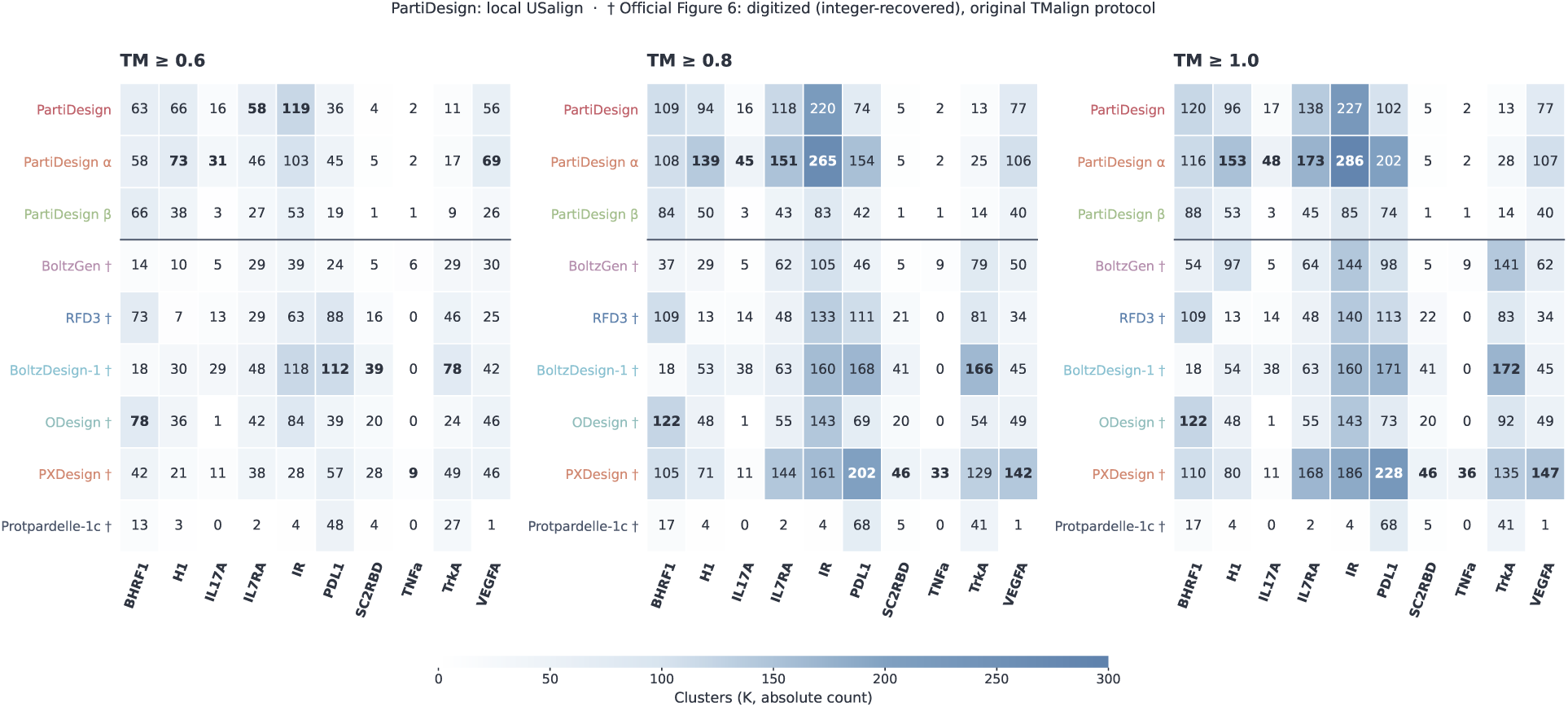
Expanded ProtDBench cluster comparison at TM0.6, TM0.8, and TM1.0. PartiDesign uses US-align; daggered public-method counts are digitized from published TM-align results. Values are absolute case-level counts of ordinary MPNN@8 successful backbones.

### D.4 Peptide topology and confidence filters

PepDBench contains DLL3, GABARAP, KEAP1, KLK2, MDM2, and XIAP in each of three topologies. Forty backbones at each integer length from 8 to 17 give 400 per case and 7,200 per PartiDesign condition. Linear peptides have open termini; cyclic peptides have head-to-tail connectivity; disulfide tasks retain the specified cysteine identities and bond. Normal, *α*, and *β* conditions share the target, length, and topology allocation. The RFD3 cyclic panel uses only CyclicMPNN, and no RFD3 disulfide evaluation is available.

AF2-IG uses model 1, one sample, and three recycles for both complex and monomer. Six conditions must hold simultaneously: complex binder-mean pLDDT ≥ 0.8, complex pTM ≥ 0.55, interface pTM ≥ 0.4, normalized interface PAE ≤ 0.3 (9.3 Å), monomer pLDDT ≥ 0.8, and bound–unbound RMSD ≤ 2.5 Å. The last metric aligns monomer and complex peptide CA coordinates and then measures all-heavy-atom RMSD. The confidence subset follows the BindCraft peptide-filter configuration, without its full energetic and surface-filtering workflow ^17^. Each configured prediction and its scope mean must pass; metrics from different predictions cannot form a passing sequence.

Self and Self-FI each have 400 sequence trials per case, ordinary MPNN@1 has 1,600, and ALL has 2,400 trials across 400 physical backbones. Across six targets, normal PartiDesign’s ALL passing-sequence counts are 1,926/14,400 linear, 1,032/14,400 cyclic, and 315/14,400 disulfide. Ordinary MPNN@1 passing for linear peptides is 1,425/9,600 (14.84%) for normal PartiDesign, 1,458/9,600 (15.19%) for BoltzGen, and 483/9,600 (5.03%) for RFD3.

Under the six-sequence ALL union, normal PartiDesign passes 449/2,400 linear, 389/2,400 cyclic, and 146/2,400 disulfide backbones (18.71%, 16.21%, and 6.08%; Fig. S3). BoltzGen passes 562/2,400, 772/2,400, and 262/2,400, respectively. RFD3 passes 239/2,400 linear and 402/2,400 cyclic backbones. The *α* counts are 435, 366, and 137 successful backbones; *β* gives 436, 410, and 139. No PartiDesign secondary-structure condition leads across all three topologies. The available Protpardelle-1c linear and RFdiffusion cyclic comparisons contribute 112/2,400 and 123/2,400 successful backbones, with all six targets retained.

**Figure S3.**
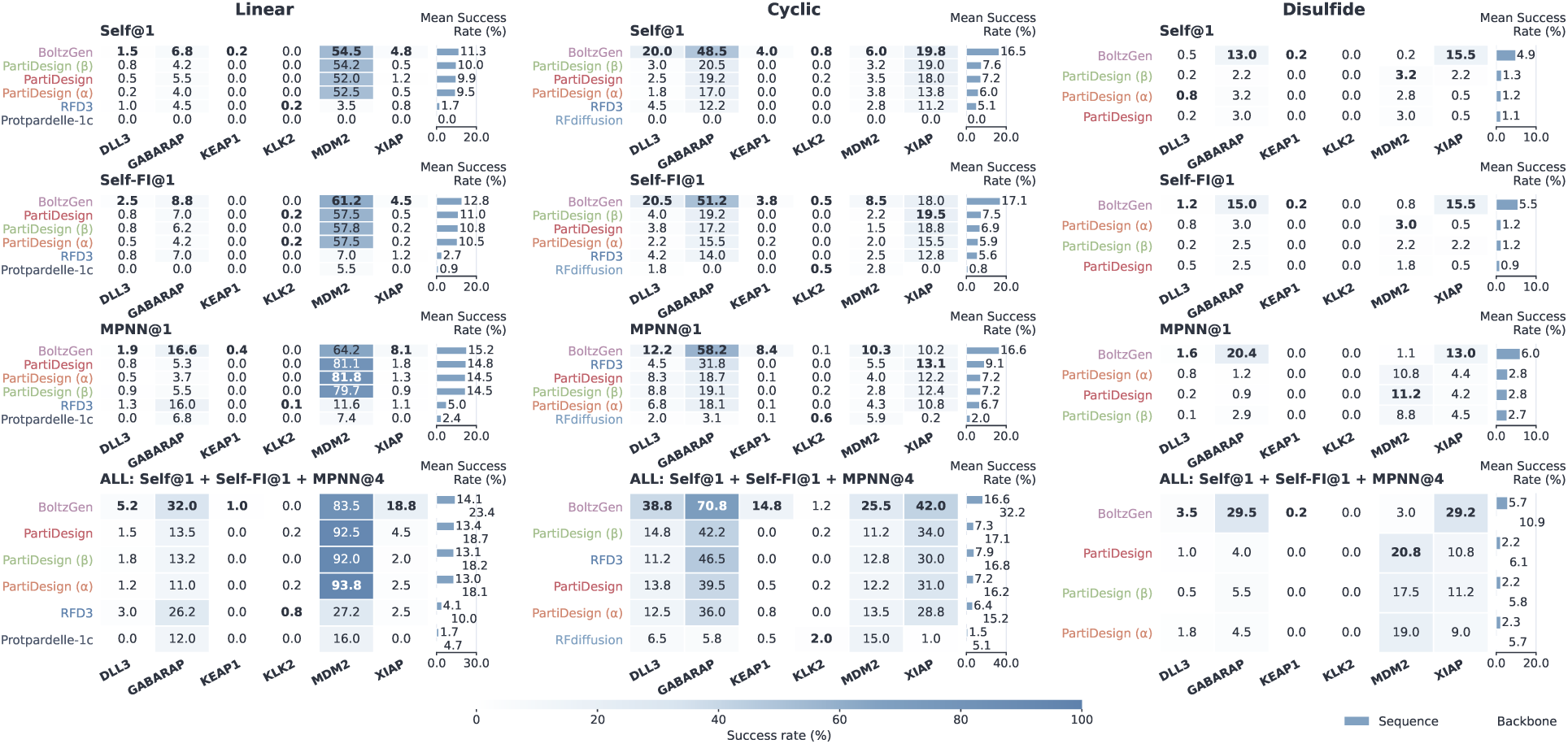
Peptide-binder passing across six targets and three topologies. Self and Self-FI each use one sequence; MPNN@1 averages four ordinary draws, and ALL is their six-sequence backbone union. Each case has 400 backbones. Linear and disulfide redesign use ProteinMPNN; cyclic redesign uses CyclicMPNN. RFD3 disulfide results are unavailable.

### D.5 Native MotifBench evaluation and score

All 30 MotifBench v1.0 problems in the three ten-problem groups are used ^23^. Each has 100 backbones and eight whole-backbone ProteinMPNN sequences per backbone at temperature 0.1 with fixed motif identities, giving 24,000 ESMFold evaluations. A sequence must have motif RMSD *<* 1 Å and scaffold C*α* self-consistency RMSD *<* 2 Å; a backbone passes if any of its eight sequences passes. Motif RMSD uses N/CA/C/O atoms, following the official implementation rather than the N/CA/C selection described in the benchmark paper.

The official scorer clusters successful backbones with Foldseek at TM threshold 0.6, alignment type 1, and alignment mode 2. It samples 50 of all 100 backbones without replacement 1,000 times and scores each subset as 110*K/*(*K* + 5), where *K* counts successful full-set clusters represented in the subset. Failed backbones remain in the sampling population. Scores are averaged across subsets and all 30 problems, without a novelty term. This clustering is used only for the official score.

PartiDesign’s displayed score is 41.86, with 1,078 passing backbones and no successes on 03_2CGA or 30_7UWL. RFdiffusion3 reports 43.52±1.35 across three runs; the leaderboard does not define the ± statistic, and PartiDesign’s single run supplies no between-run error estimate. Software and checkpoint versions are not fully specified. No compute-budget estimate is reported for this task.

### D.6 Indexed atomic-motif enzyme design

All 41 RFdiffusion2 AME cases are used ^26^. Assigning motif residue indices and contigs produces 180-residue designs with fixed motif positions, restricting placement freedom relative to the original unindexed benchmark. The case counts for one through seven discontinuous residue islands are 2, 4, 12, 11, 5, 6, and 1; each island is a residue segment, not an individual catalytic atom.

Each of the three method/condition rows has 4,100 backbones and 41,000 sequence evaluations with ESMFold2-standard; six additional normal-PartiDesign AF3 cases are excluded. The transferred Complexa criterion ^14^ requires design-chain N/CA/C self-consistency RMSD ≤ 2 Å, supplied-reference motif-atom RMSD ≤ 1.5 Å, correct motif residue identities, and no protein–ligand atom distance ≤ 1.5 Å. The collision test includes covalent or bonded neighbours and its boundary. All criteria must hold in one complete prediction, without an additional ligand-pose RMSD, pLDDT, interface pTM, or PAE gate.

Self passing is 484/4,100 (11.80%) for normal PartiDesign and 702/4,100 (17.12%) for *α*, compared with 79/4,100 (1.93%) for RFD3 (Fig. S4). Ordinary MPNN@1 passing is 5,605/32,800 (17.09%), 7,491/32,800 (22.84%), and 4,103/32,800 (12.51%), respectively. The sequence-level advantage persists after redesign with indexed inputs and ESMFold2 evaluation.

Across all ten sequences, normal PartiDesign, *α*, and RFD3 yield 6,736, 9,064, and 4,579 passing sequences out of 41,000, respectively. Their ALL successful-backbone counts are 1,563/4,100, 1,893/4,100, and 1,310/4,100. These rates use the transferred predictor and indexed inputs, which differ from the original RFdiffusion2 and Complexa evaluation protocols.

**Figure S4.**
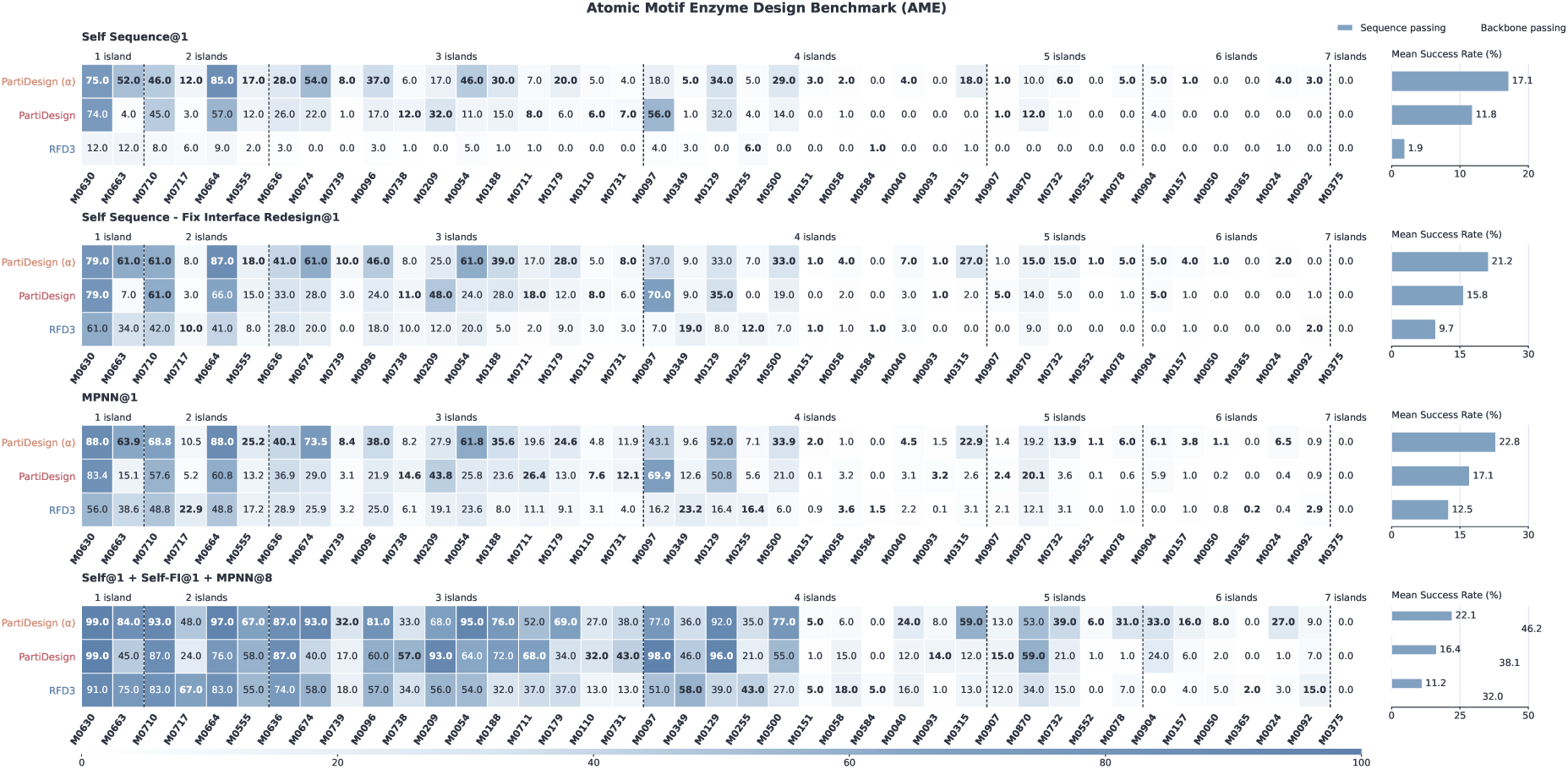
AME passing across all 41 atomic-motif problems, grouped by discontinuous residue-island count. Each case has 100 backbones. MPNN@1 averages eight ordinary sequences; ALL joins Self, Self-FI, and eight ordinary sequences. Results use ESMFold2-standard and indexed AME inputs.

### D.7 Protein design with nucleic-acid context

The five DISCO-derived inputs are DNA 7EDS (F/R, 30+30 nucleotides), 7S03 (B/C, 26+26), and 9JPJ (A/B, 27+27), and RNA 6YMC and 8BWT (A, 26 each) ^29^. Native proteins and other components are removed. Model 6 of the 8BWT NMR ensemble is selected as the nucleic-acid P-RMSD medoid, independently of design success. We select four-nucleotide hotspot patches F8–9/R22–23, B17–18/C8–9, A14–15/B14–15, and A12/A14–16 for either RNA; these are not published DISCO annotations. Full-chain contexts and 124 requests per case differ from DISCO’s cropped contexts and 100-request protocol.

PartiDesign uses seed 2026, batch size 16, 400 sampling steps, and flexible context constraints. Four requests at each integer protein length 50–80 give 124 backbones per case. ESMFold2-standard uses standard mode, one sample, 100 steps, 20 loops, language-model dropout 0.3, and seed 101. A single rigid least-squares alignment fits all corresponding protein C*α* and valid nucleic-acid P points with equal weights, requiring total RMSD *<* 5 Å. All polymer chains must map correctly. Genuinely absent reference 5*^′^* phosphates are omitted; missing expected atoms or incomplete correspondence make the metric unavailable. No separate chain fits or extra confidence or collision thresholds are used.

The joint fit differs from published nucleic-acid-anchored protein RMSD and protein-anchored nucleic-acid RMSD screens. Longer contexts contribute more fitted points, so low joint RMSD can coexist with interface errors. Primary passing-sequence totals are 277/3,720 for normal PartiDesign, 314/3,720 for *α*, 523/3,720 for BoltzGen, and 113/3,720 for RFD3. Ordinary MPNN@1 passing is 221/2,480 (8.91%), 253/2,480 (10.20%), 369/2,480 (14.88%), and 93/2,480 (3.75%), respectively.

The six-sequence union passes 140/620 (22.58%) normal and 149/620 (24.03%) *α* PartiDesign backbones, compared with 205/620 (33.06%) for BoltzGen and 75/620 (12.10%) for RFD3 (Fig. S5). Target dependence is pronounced: normal PartiDesign passes 50/124 union backbones on DNA 7EDS and 47/124 on RNA 8BWT, whereas BoltzGen passes 18/124 and 19/124; on RNA 6YMC, BoltzGen passes 101/124 and normal PartiDesign 6/124.

**Figure S5.**
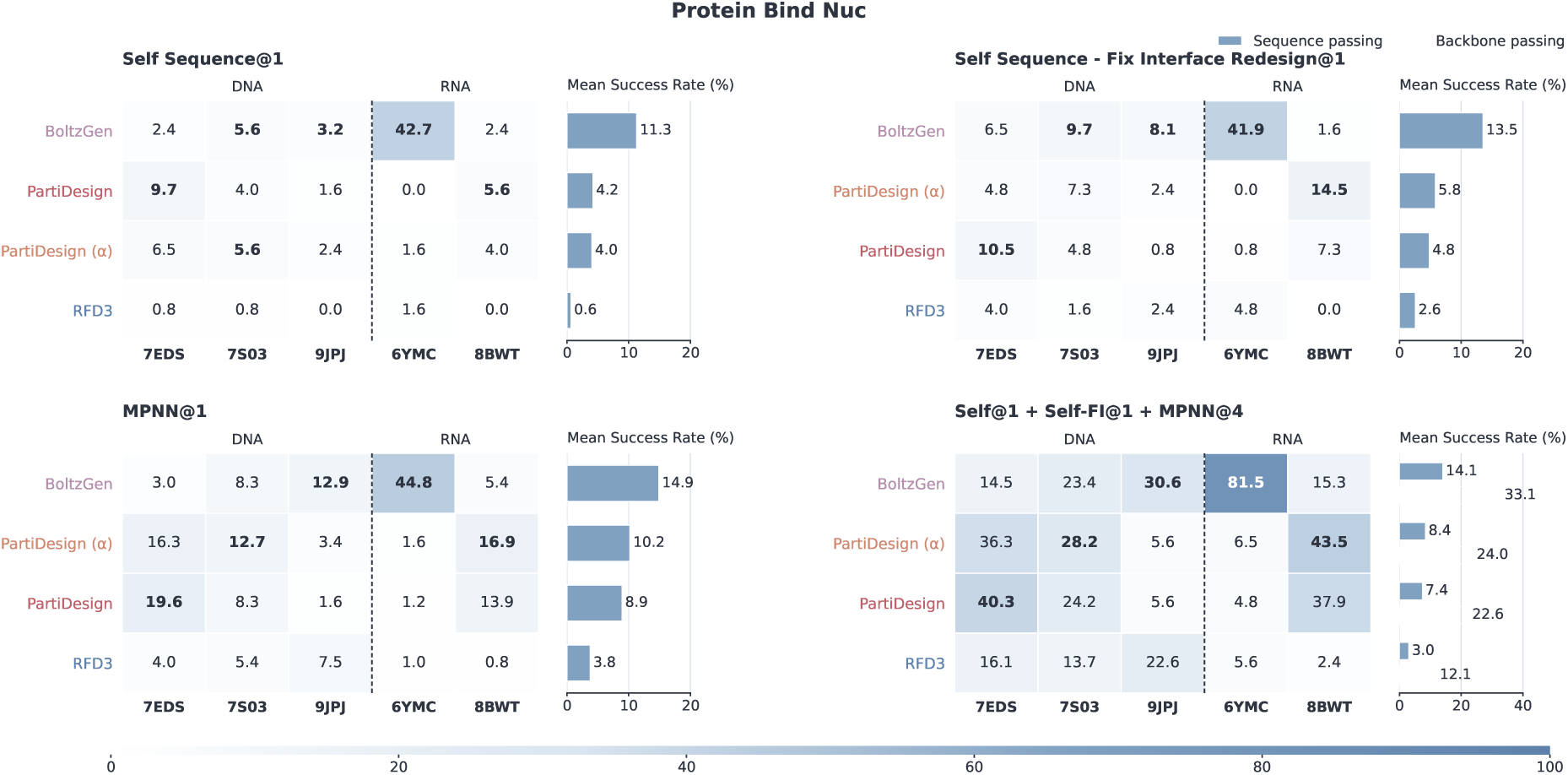
Protein binders for three DNA and two RNA targets under the project-specific joint CA/P RMSD filter and ESMFold2-standard evaluation. Each case has 124 backbones. Self and Self-FI each use one sequence, ordinary MPNN@1 averages four draws, and Union reports their six-sequence backbone union.

### D.8 Protein design with a specified ligand

The four Complexa/NVIDIA inputs are 5SDV–IAI, 7BKC–FAD, 7C7M–SAM, and 7V11–OQO ^14^. Only ligand identity and conformation are supplied; the original protein is removed. Each method or condition generates 200 new 100-residue backbones for each of four ligands, excluding joint-SMILES designs and additional example ligands. Normal and *α* PartiDesign, BoltzGen, and RFD3 each evaluate Self, Self-FI, and one ordinary LigandMPNN sequence per backbone.

Three strict conditions must hold in one ESMFold2-standard prediction: minimum directional protein–ligand PAE *<* 2 Å, protein C*α* self-consistency RMSD *<* 2 Å, and protein-aligned ligand heavy-atom RMSD *<* 5 Å. PAE is the minimum over protein-to-ligand and ligand-to-protein token blocks, excluding context–context entries, rather than a directional average or all-contact constraint. Ligand RMSD fits identity-matched protein heavy atoms excluding OXT, with a C*α* fallback when needed, then measures matched ligand heavy atoms. The separately calculated C*α*-only ligand RMSD is not used. Both RMSDs compare generated and refolded complexes. Incomplete chemical or atom mapping makes the metric unavailable. No extra pLDDT, interface pTM, mean PAE, or collision gate is applied. These Complexa-inspired thresholds are transferred to ESMFold2-standard, whose confidence calibration can differ.

Normal PartiDesign Self passes 175/800 sequences (21.88%), compared with 87/800 (10.88%) for BoltzGen and 21/800 (2.63%) for RFD3 (Fig. S6). Ordinary LigandMPNN@1 reverses this ordering: 81/800 (10.13%) for normal PartiDesign, 144/800 (18.00%) for BoltzGen, and 103/800 (12.88%) for RFD3. Their three-sequence ALL unions pass 186/800 (23.25%), 201/800 (25.12%), and 136/800 (17.00%) backbones. The *α* condition reaches 153/800 (19.12%), below normal PartiDesign. Normal Self success is concentrated in 5SDV (142/200), with 1/200 for 7BKC, 2/200 for 7C7M, and 30/200 for 7V11.

**Figure S6.**
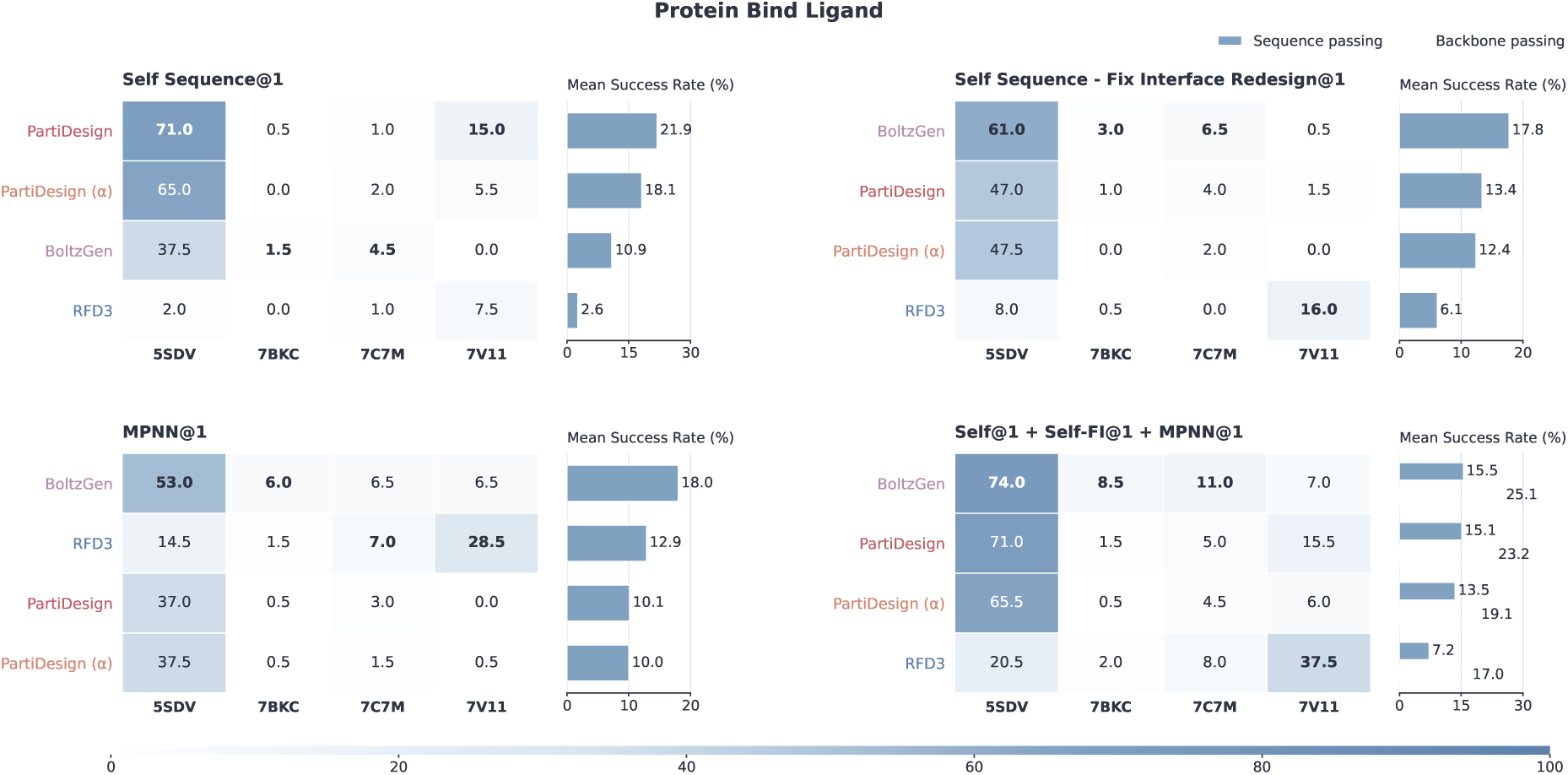
Protein design around four specified small molecules. Each ligand has 200 new protein backbones per method or condition. Self, Self-FI, and ordinary LigandMPNN@1 each use one sequence; ALL is their three-sequence backbone union. All displayed results use ESMFold2-standard and the transferred Complexa ligand-binder criteria.

### D.9 Task-specific standardized compute budgets

Each case independently receives one 4090-48G GPU for 24 hours. Costs cover steady-state model computation, excluding loading, compilation, warmup, CPU features, I/O, transfers, and scheduling. Generation cost is batch-size-16 latency divided by 16; redesign costs one sequence per backbone. AF2-IG validation includes complex and monomer predictions; ESMFold2-standard tasks use their complete validation cost. Timings use a shared 50:50 context/design fixture rather than case-specific compositions, with case lengths mapped to the timing curves.

ProtDBench’s Single Sequence@1 protocol costs *g* + *v* for Self and *g* + *m* + *v* for either redesign branch. Sequence and backbone yields coincide. Ordinary MPNN probability is estimated from eight draws, not their MPNN@8 union. Other protein budgets compare Self with Self plus one ordinary sequence at costs *g* + *v* and *g* + *m* + 2*v*. Backbone yield uses their paired OR probability; sequence yield uses the sum of marginal probabilities. Self-FI and larger ALL unions are excluded. Only methods with timing measurements receive estimates.

Task budgets appear in Figs. 10, 12, 13, and 14. Protein–nucleic-acid estimates use ESMFold2-standard, excluding secondary AF3 evaluations. Added validation can raise passing-sequence output while lowering successful-backbone output when its cost outweighs the gain in OR probability. These are model-compute estimates, not end-to-end screening times.

## E Cross-modal design protocols and extended results

### E.1 Shared nucleic acid evaluation

AF3 uses one complex sample, ten recycles, seed 101, and Triton attention. NA–protein evaluation uses cache-only MSA handling with templates enabled. NA–ligand evaluation enables templates and uses automatic/cache-reuse mode for normal PartiDesign, and cache-only mode for paired PartiDesign and RFD3NA. Protein/RNA contexts may use matching cached features; designed chains are query-only, and DNA and ligand chains use no native MSA search. Automatic mode can reuse caches without a fresh search per candidate. Additional ESMFold2 evaluations do not enter these AF3 summaries.

Sequence and union-backbone rates use classified sequences and determinable backbones, respectively. A passing branch establishes backbone success; otherwise a missing branch leaves it unknown. Case-macro means weight cases equally. Both gates compare generated complexes with sequence-specific AF3 predictions, without extra ipTM, pLDDT, PAE, or clash thresholds. Genuinely absent reference 5*^′^* phosphates are omitted; missing expected points, incomplete chemical mapping, or nonfinite coordinates make a metric unavailable.

### E.2 Protein-conditioned DNA and RNA

The twelve NACraft NA-12 targets ^30^ derive from post-13-January-2023 PDB releases, excluding ambiguous sequences, excessive protein length or chain count, and unavailable coordinates. Table S6 lists design lengths and observed protein inputs. Preparation removes native nucleic acid, water, ions, and additives without reconstructing unresolved protein residues. Ten inputs with ambiguous SEQRES/coordinate alignment omit SEQRES while preserving observed coordinates, author numbering, insertion codes, alternate locations, chain breaks, and connectivity. Observed protein inputs can therefore be shorter than the reported full sequences.

Each method/variant has 1,200 backbones and 7,200 AF3-evaluated sequences: Self, Self-FI, and four ordinary NA-MPNN draws per backbone. Self-FI fixes generated design identities at the 6.0 Å context interface and redesigns the remaining positions at temperature 0.1. Ordinary redesign also uses NA-MPNN ^32^. The complete six-sequence union is distinct from an ordinary MPNN@1 sequence rate. PartiDesign normal and paired generation use seed 2026, native batch size 16, 400 sampling steps, and flexible context constraints. The paired condition is coarse and does not encode a specified base-pair map.

The NA–protein gate fits all corresponding protein C*α* and valid observed NA-P points in one rigid transformation and requires combined RMSD *<* 5.0 Å. Equal point weights give longer proteins more influence. This whole-complex consistency screen does not measure native-binder recovery and can miss interface errors.

PartiDesign achieved a case-macro Self passing rate of 23.67%, compared with 15.75% for RFD3NA (Fig. S7). The corresponding ordinary NA-MPNN sequence rates were 17.67% and 15.17%. Allowing any of all six evaluated sequences to pass increased backbone coverage to 37.25% for PartiDesign and 21.42% for RFD3NA. The paired variant reached 37.08%, providing no aggregate improvement over normal generation. Results varied substantially by target: normal PartiDesign passed at least one sequence for 100/100 backbones on DNA target 8U0P and 97/100 on RNA target 9NY9, but 0/100 on DNA target 7UXY. RFD3NA exceeded normal PartiDesign on RNA targets 8SWB and 8X0N. The average advantage thus coexists with target-specific reversals and difficult targets for both methods.

**Table S6.** NA–protein input scope. Protein lengths refer to observed input residues and the NA-12 reported protein sequence, respectively. The source nucleic acid sets design length but is removed before generation.

| PDB | Design | Length (nt) | Protein length (aa) |  |
|---|---|---|---|---|
|  |  |  | Observed | Reported |
| 7UV7 | DNA | 21 | 212 | 263 |
| 7UXY | DNA | 32 | 213 | 263 |
| 8C58 | DNA | 14 | 376 | 395 |
| 8SLN | DNA | 29 | 240 | 291 |
| 8U0P | DNA | 11 | 322 | 340 |
| 8VX4 | DNA | 35 | 311 | 387 |
| 8SWB | RNA | 20 | 149 | 149 |
| 8SWC | RNA | 20 | 149 | 294 |
| 8X0N | RNA | 12 | 181 | 202 |
| 9NY9 | RNA | 19 | 349 | 357 |
| 9RVP | RNA | 34 | 123 | 137 |
| 9XZR | RNA | 71 | 176 | 293 |

**Figure S7.**
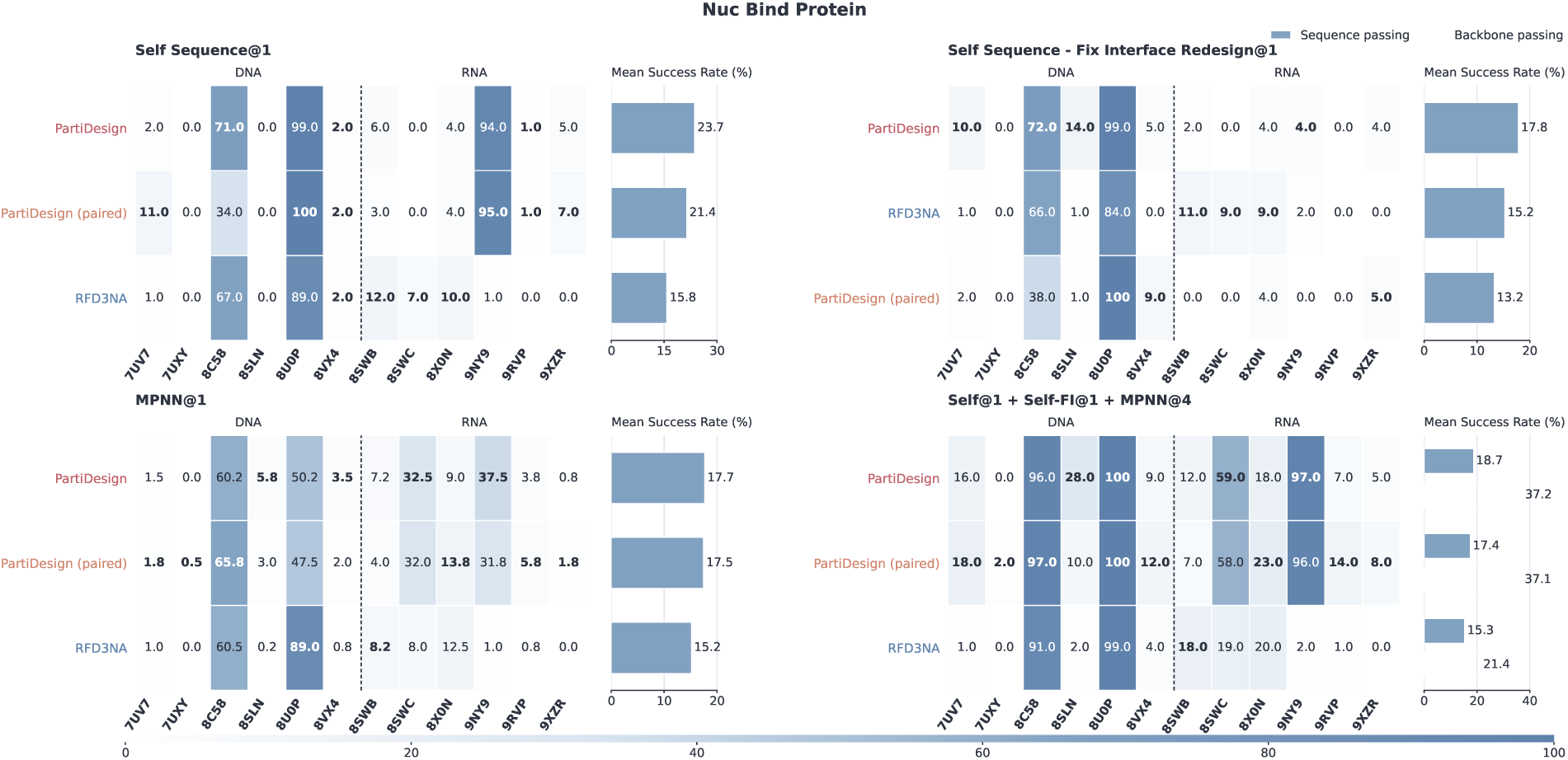
Protein-conditioned DNA and RNA design. Each of twelve NA-12 targets has 100 backbones per method/variant, evaluated with AF3 for Self, Self-FI, and four ordinary NA-MPNN sequences. MPNN@1 is the ordinary sequence rate; Union reports six-sequence backbone coverage and sequence passing. Summaries weight cases equally. Combined protein-C*α*/NA-P RMSD must be *<* 5 Å, without extra confidence or clash filters.

### E.3 Small-molecule-conditioned DNA and RNA aptamer design

Ethyl maltol, imidacloprid nitroimine, and vanillin each condition DNA20 and RNA30 designs. Each method/variant has 600 backbones and 1,200 planned sequences: Self and Self-FI, without ordinary NA-MPNN. The supplied SMILES defines ligand chemistry, including the specific nitroimine identity. These six settings use lengths selected from earlier ESMFold2 results and are not a prospectively blinded benchmark. PartiDesign uses seed 2026, batch size 16, 400 sampling steps, and flexible context constraints, with scoring and postprocessing disabled. Fixed-interface NA-MPNN preserves ligand-proximal nucleotide identities but does not consume ligand atom features.

The dual-RMSD gate first fits the designed nucleic acid’s valid P points and requires NA-P self RMSD *<* 5.0 Å. The identical transformation is applied to the ligand, whose complete heavy-atom RMSD must also be *<* 5.0 Å. Both conditions must hold in the same prediction; the ligand is never fitted independently. The two criteria jointly assess preservation of the aptamer fold and the designed ligand pose.

Normal PartiDesign reached 61.33% Self sequence passing and 66.50% any-of-two backbone coverage, compared with 18.33% and 29.19% for RFD3NA (Fig. S8). The paired variant reached 54.33% and 58.17%, respectively. Normal PartiDesign’s union coverage ranged from 91/100 for DNA–vanillin to 9/100 for RNA–imidacloprid nitroimine; RFD3NA reached 33/100 on the latter case. The difference between the DNA and RNA results shows that a result for one nucleic acid chemistry does not transfer uniformly to the other, even with the same ligand.

One RFD3NA DNA–ethyl-maltol Self-FI redesign failed, leaving the only unknown primary AF3 classification. This case has 100 Self and 99 Self-FI evaluations, 199 classified sequences, and 13 passing/99 determinable backbones. Its planned-100-backbone interval is 13–14%. Missing supplementary ESMFold2 results do not alter the AF3 denominator.

**Figure S8.**
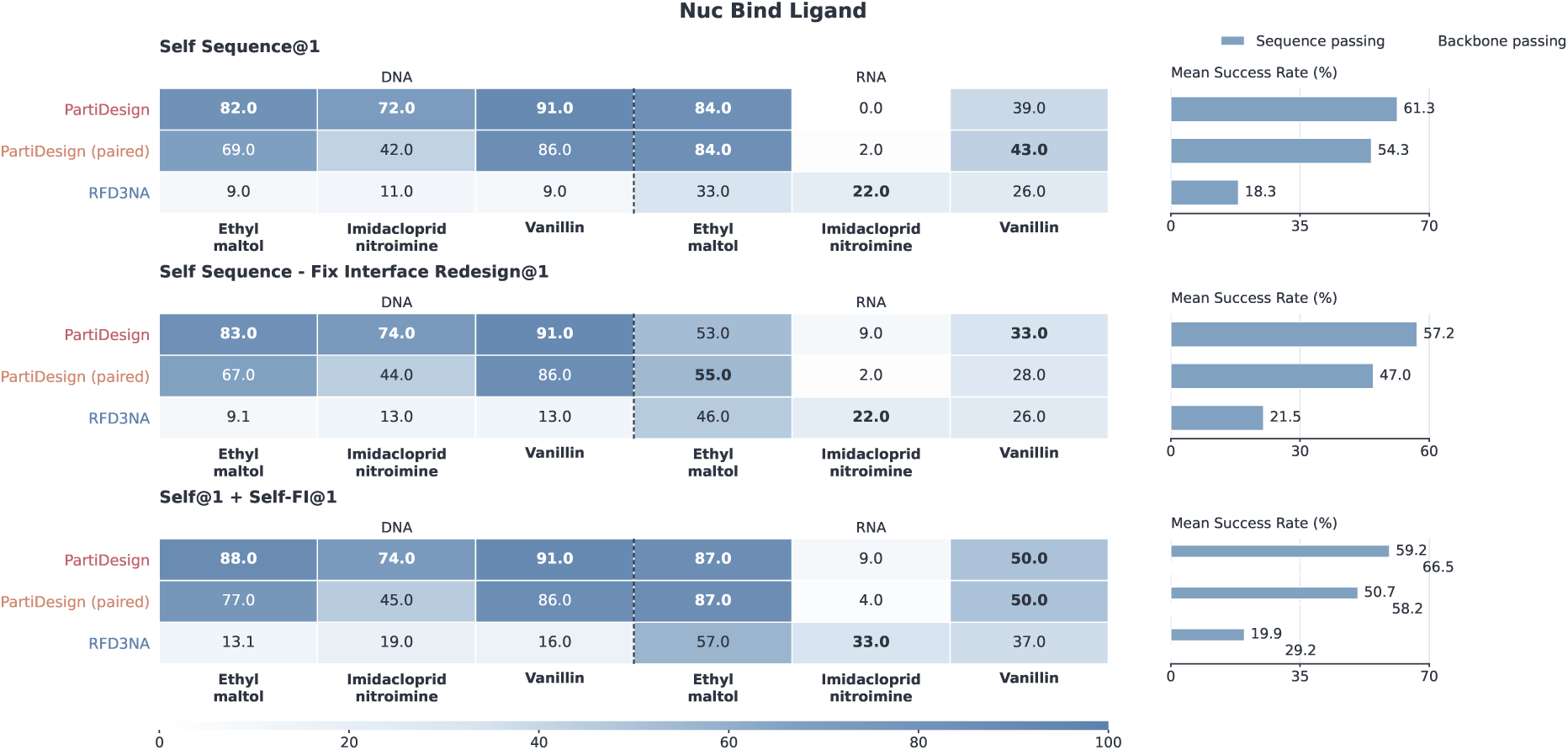
Small-molecule-conditioned DNA and RNA design. Three fixed ligands are paired with DNA20 and RNA30 designs, with 100 generated backbones per case and method/variant. Self and Self-FI are the only evaluated sequence branches. Both NA-P self RMSD and NA-aligned ligand-heavy-atom RMSD must be *<* 5 Å in the same AF3 prediction. Union is any-of-two backbone coverage. Case-macro summaries retain the RFD3NA DNA–ethyl-maltol redesign failure as unavailable, giving a 99-backbone determinable denominator for that union cell.

### E.4 Task-specific nucleic acid budget interpretation

Figures 15 and 16 allocate one 4090-48G GPU for 24 hours per case, using fine-64 AF3 timings. The NA–protein two-sequence protocol joins Self with one uniformly selected ordinary NA-MPNN draw, estimated from four draws paired with the same backbone’s Self result. It does not use the six-sequence union probability. The NA–ligand protocol instead joins Self and Self-FI, charging one redesign and two AF3 calls without ordinary MPNN.

AF3 cost maps the canonical-token proxy *N* to *B* = max(128, 64⌈*N/*64⌉) and uses the measured median for that padded shape: 128, 192, 256, 320, 384, 448, or 512. Structural evaluation used coarser padding than these fine-64 timing runs. Timings use empty MSA, one sample, ten recycles, and Triton attention, excluding CPU feature construction and cache/search work. The protein-route generation fixture approximates the nucleic-acid route despite different case compositions. Normal PartiDesign’s two-sequence case-macro estimates are 1,968 passing NA–protein backbones and 2,576 passing sequences per day, versus 15,761 backbones and 28,104 sequences for NA–ligand. The unknown RFD3NA redesign propagates to bounds, not a filled point estimate.

### E.5 Ligand generation, chemical screening, and literature budget scope

The five ligand-design inputs use pocket structures, eight explicit hotspots per case, and 200 requested attempts. Exact matching to the external crop/hotspot collection was not verified. ODesign’s sixth target, 7BKC, was not generated with PartiDesign. Generation uses seed 2026, 400 steps, flexible context coordinates, scoring and postprocessing enabled, and a sampled ligand heavy-atom count.

Graph reconstruction and postprocessing precede RDKit recognition, QED calculation, and CPU docking with Vina 1.2.7 and Meeko 0.8.0, using one CPU, seed 101, exhaustiveness 8, and 20 poses. Validity requires RDKit recognition and successful docking; passing additionally requires QED *>* 0.25 and best successful Vina affinity *<* −8.18 kcal/mol for the same candidate, with equality failing both strict thresholds. This screen follows ODesign ^7^, using QED and Vina as defined in their original studies ^34,35^. No sequence redesign or refolding is performed.

Across 1,000 planned attempts, 945 generated candidates were valid and 578 passed the complete screen (Fig. S9). Validity was 94.5% and planned-attempt passing was 57.8%. The denominator includes 55 generation failures, whose scientific metrics remain unavailable, and 367 valid candidates that failed the QED/Vina gate. Passing rates were 80.5% for both 2VT4 and 5SDV, 70.5% for 6CM4, 2.0% for 7C7M, and 55.5% for 7V11. On 7C7M, 175/200 attempts yielded valid dockable molecules, but only 4/200 passed the complete screen. Chemical validity and passing quality therefore differ sharply for this pocket.

**Figure S9.**
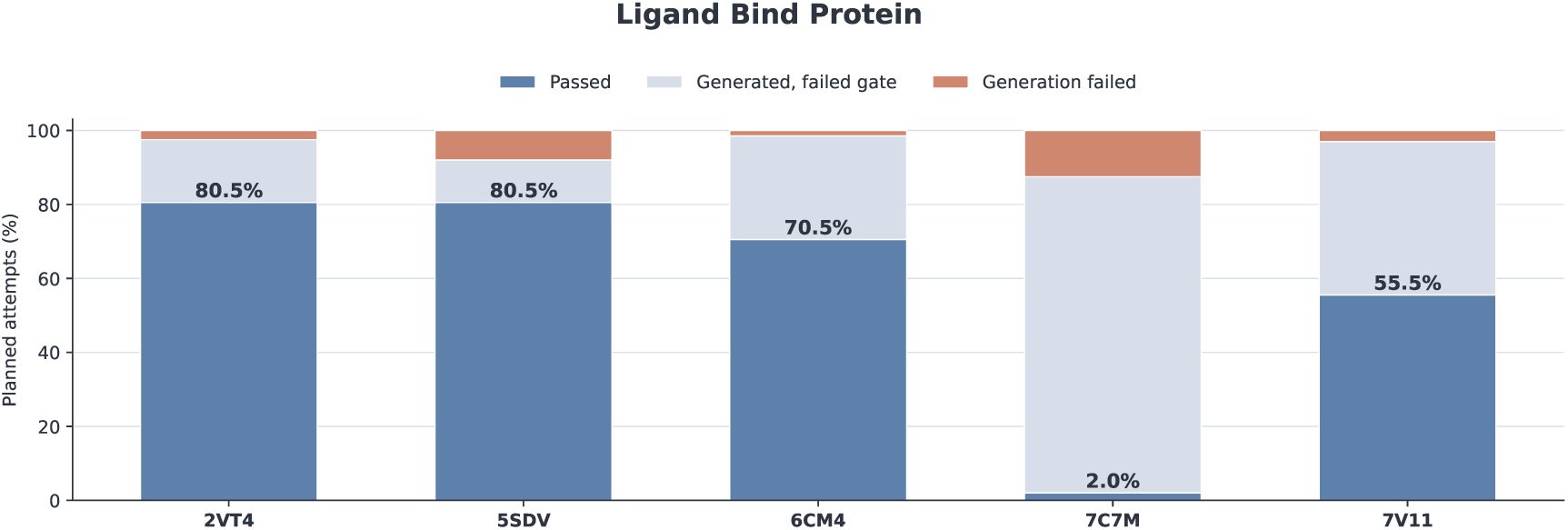
Protein-conditioned ligand generation across five pockets. Each stacked bar uses all 200 planned attempts. Validity requires both RDKit recognition and successful Vina docking; passing also requires QED *>* 0.25 and Vina Dock *< −*8.18 kcal/mol for the same postprocessed candidate. The three categories distinguish passing candidates, valid candidates failing the remaining gate, and generation failures. Overall, 945/1,000 attempts are valid and 578/1,000 pass. No refolding or native-ligand RMSD filter is used.

Table S7 records all planned-attempt outcomes.

The ligand budget averages batch-size-16-amortized generation cost across all 200 planned molecular sizes, including failures. Multiplying 86400*/g* by valid/200 or passed/200 gives valid or passing candidates per generation-only GPU-day. The measured protein-route fixture approximates the ligand route. CPU QED/Vina times include preparation, docking, and result serialization but exclude input loading. Mean costs per planned attempt are 104.450, 123.935, 100.570, 106.645, and 122.840 s in table order. Times have one-second resolution with unspecified CPU hardware and contention; a 9,891 s attempt contributes to the 2VT4 mean. GPU estimates exclude CPU cost and concurrency.

**Table S7.** Ligand generation and planned-attempt screening. Valid candidates all have successful docking results. Generation failures have unavailable metrics and contribute no passes among the 200 planned attempts.

| Pocket | Planned | Valid | Passed | Generation failures |
|---|---|---|---|---|
| 2VT4 | 200 | 195 | 161 | 5 |
| 5SDV | 200 | 184 | 161 | 16 |
| 6CM4 | 200 | 197 | 141 | 3 |
| 7C7M | 200 | 175 | 4 | 25 |
| 7V11 | 200 | 194 | 111 | 6 |
| Total | 1,000 | 945 | 578 | 55 |

Figure 17 includes 60 digitized ODesign Figure 4a aggregates for six methods across five shared pockets: H100 sampling-plus-docking estimates from 100 molecules per target, compared with PartiDesign’s 200 planned attempts and 4090 generation-only proxy cost. Eight bars below raster resolution have positive upper bounds marked by downward triangles. Digitization intervals describe image resolution, not sampling uncertainty; these aggregates provide neither sample-level outcomes nor a hardware-normalized speedup.

## F Steady-state timing and case-wise compute budgets

### F.1 Workloads and timing boundaries

Timing workloads vary the total token count *N* = *N*_context_ + *N*_design_ from 50 to 500 in increments of 50, with a 50:50 context/design split. PartiDesign, BoltzGen and RFD3 use protein-design fixtures; RFD3NA uses RNA design. Generation is measured at native batch size 16, refolding per complete validation sample, and MPNN per sequence designed for one backbone. Each profile therefore retains its own molecular workload and timing unit.

All profiles use one 4090-48G GPU; refold timing also assigns one CPU worker. Inputs and native configurations are fixed. Models are loaded, compiled where applicable and warmed up before formal measurements, with device synchronization around each timed invocation. The boundary excludes loading, compilation, warmup, CPU feature processing, host/device transfer, file I/O and scheduling. PartiDesign generation includes context VAE encoding, the solver and decoding. CPU proposal construction, molecular assembly, output writing, chemical reconstruction, relaxation and optional scoring lie outside this interval.

PartiDesign uses 400 flow steps, automatic checkpoint routing to the protein member for this fixture, flexible context and Inductor compilation. RFD3 and RFD3NA use 200 actual sampler steps at native precision. BoltzGen’s diverse and adherence weights are measured separately with 500 steps, three recycles and 16 diffusion samples. Latency ratios compare these native inference recipes, whose step counts and molecular workloads differ.

The timing table contains 49 generation points and 20 non-AF3 refold points, each with ten formal repeats. RFD3NA runs out of GPU memory at *N* = 500. One complete AF2-IG measurement includes both complex and isolated designed-chain predictions and is charged once per validation. ESMFold2-standard is measured at its complete-sample call boundary.

### F.2 AF3 timing measurements

AF3 timing uses empty MSA, one prediction sample, ten recycles, Triton attention and CPU1/GPU1. This compute fixture is distinct from the cached-context inputs used for scientific evaluation. We measured fifteen input lengths using 64-token bucket increments, with 20 repeats per length. Figure S10 shows the raw measurements against actual input length.

For the budget calculation, a case with *N* ≤ 512 uses bucket size *B* = max(128, 64⌈*N/*64⌉). Table S8 lists the measured bucket medians. Inputs assigned to the same bucket have the same estimated validation cost; short inputs use the smallest measured bucket, B128.

**Table S8.** Measured AF3 bucket costs used for budget calculations. The intervals are the 10th–90th percentiles of repeated timings.

| Bucket size *B* | Formal calls | Median (s) | p10–p90 (s) |
|---|---|---|---|
| 128 | 40 | 1.705526 | 1.703231–1.707275 |
| 192 | 40 | 2.282435 | 2.276628–2.296040 |
| 256 | 60 | 3.344011 | 3.335972–3.348155 |
| 320 | 40 | 4.778231 | 4.773346–4.783769 |
| 384 | 40 | 7.141255 | 7.135227–7.144864 |
| 448 | 40 | 9.124284 | 9.116376–9.131211 |
| 512 | 40 | 11.765804 | 11.760666–11.774885 |

We compared nonnegative-coefficient polynomials of degree one to three using leave-one-bucket-out error. The selected descriptive quadratic is

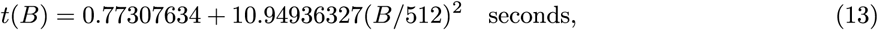

over 128 ≤ *B* ≤ 512, with leave-one-bucket-out RMSE 0.2303 seconds and MAPE 6.04%. The main-text smooth AF3 curve uses this fit. Budget calculations use the measured bucket medians.

**Figure S10.**
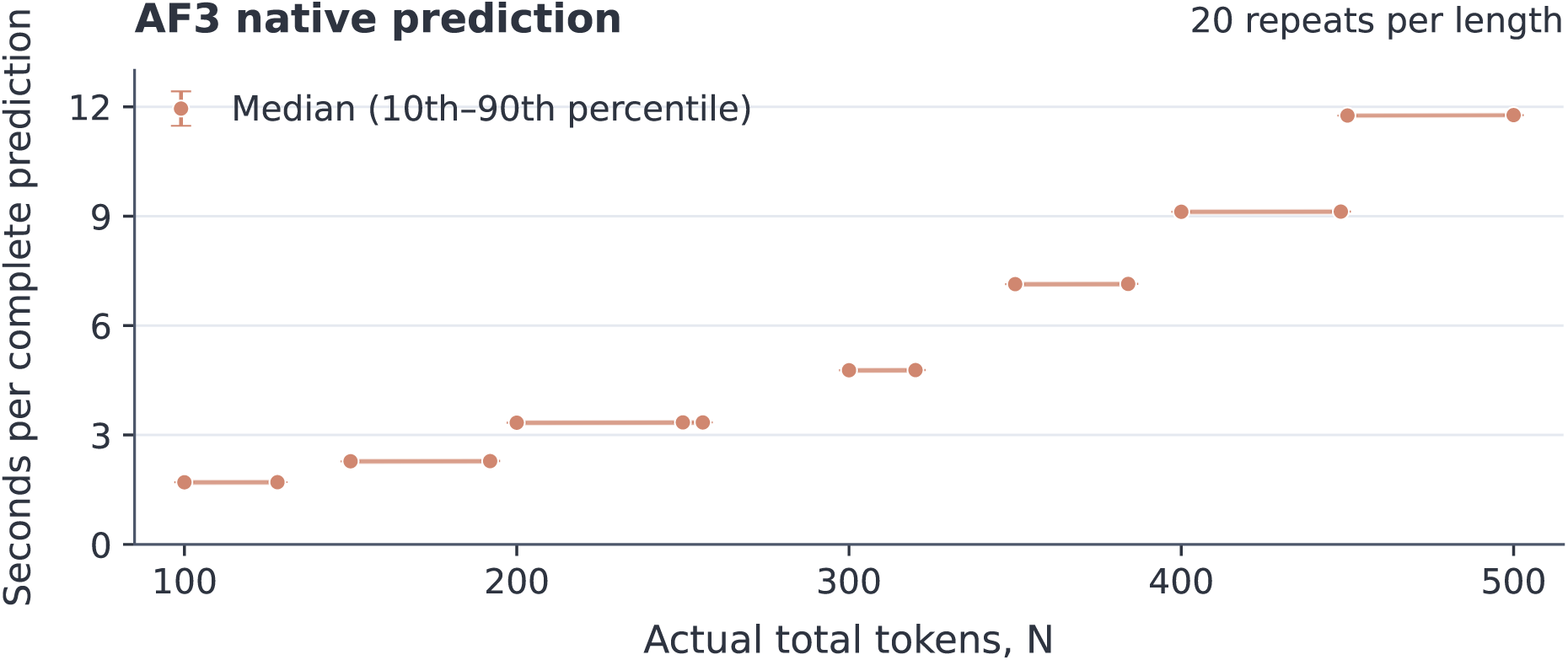
Raw AF3 timing measurements. Points show the median of 20 complete predictions at each actual input length, with 10th–90th percentile error bars. Short segments join input lengths assigned to the same bucket. The budget uses the bucket medians in Table S8.

### F.3 Single-sequence redesign costs

The MPNN timing table contains 595 fixed conditions: 318 LigandMPNN, 113 ProteinMPNN, 95 NA-MPNN, 39 CyclicMPNN and 30 motif ProteinMPNN. Each condition measures one sequence designed for one backbone and can share a target with other conditions. Each condition uses five warmup calls and 20 measured calls, totaling 2,975 warmup and 11,900 measured calls. Protocols without sequence redesign omit the MPNN call.

Per-condition medians and the recorded task–backend association define *m_c_*. The plotted intervals summarize repeated compute calls; MPNN uses empirical percentiles and AF3 uses linear interpolation.

### F.4 Cost and passing probability for each protocol

For a case, average the generation and validation cost over its declared length distribution and native profile. The generation contribution is a batch-of-16 latency divided by 16. Self costs *g* + *v*. A single redesigned sequence alone costs *g* + *m* + *v*. A two-sequence screening protocol costs *g* + *m* + *v*_Self_ + *v*_R_, with the same-cost form *g* + *m* + 2*v* used when applicable. Self-FI also requires one MPNN call; its fixed-identity constraints change the sequence protocol, not the number of sequence-design calls.

Each budget retains the sequence branches evaluated for that task. Most two-sequence comparisons pair Self with one uniformly selected ordinary MPNN redesign; nucleic-acid binding to ligands pairs Self with its available Self-FI sequence. ProtDBench evaluates Self, Self-FI and ordinary MPNN@1 as separate single-sequence protocols.

To estimate the two-sequence success probability when *K* ordinary draws are available, define *s_i_* ∈ {0, 1} for the Self outcome and *q_i_* = *K*^−1^ ∑_*k*_ *I_ik_* for the conditional success of one uniformly selected redesign. Then

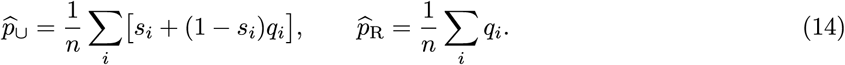

For a single Self-FI draw, *q_i_* is its binary outcome. The estimate preserves the empirical dependence between Self and redesign on each backbone. Successful backbones per GPU-day are 86400*p̂∪*/*C*; successful sequences per GPU-day are 86400(*p̂*Self + *p̂*R)/*C*. The two counts coincide for a single-sequence protocol. Unresolved outcomes retain their classification bounds.

Each case receives its own 24-hour allocation before averaging across cases; classification bounds are retained for incomplete outcomes. Costs are standardized estimates from shared fixtures. Non-AF3 costs use the recorded length-to-cost fits and recipe associations. Requests outside the measured 50–500-token grid retain their extrapolation flag. BoltzGen budgets for ProtDBench, PepDBench and protein binding to nucleic acids or ligands use an equal mixture of diverse/adherence latencies, including experiments that used the diverse member. RFD3 budgets use the current native timing curve as the standardized cost profile. These fixtures approximate case costs using a common native inference configuration.

PartiDesign generation uses the protein timing fixture as a cross-member proxy for the omni member in AME, the nucleic-acid and ligand members, and peptide experiments with unavailable historical member records. Cost can also vary with molecular composition and context/design ratio at a fixed token count. Ligand budgets charge GPU generation only and report QED/Vina CPU costs separately. The ODesign literature estimates instead use H100 sampling plus docking and a different attempt budget, so they provide a qualitative comparison across timing scopes and hardware. All daily outputs retain the stated compute boundary, excluding loading, compilation, optional relaxation, candidate scoring and multiprocess scheduling.

## G Extended docking capabilities

Fixed-identity docking tests recovery of a specified molecule’s bound pose. We examine ssRNA on protein receptors, small molecules on nucleic acid receptors, protein–ligand docking, and four cyclic-peptide examples. Molecular identities are fixed; native design coordinates are used only for evaluation. PartiScore ranks the saved refined PartiDesign candidates, while receptor-aligned pose RMSD measures agreement with the experimental reference.

### G.1 ssRNA and nucleic-acid-receptor docking

Protein–ssRNA docking predicts the bound placement and conformation of a specified RNA sequence. The nine examples comprise resolved, unpaired fragments from structures released between June 2022 and January 2023: 7F3I, 7F3J, 7SOS, 7SOV, 7TUV, 7UU4, 7V2Z, 7XS4, and 8EDJ. Each fragment contains four to six contiguous observed nucleotides, has no detected base pairs, and places every nucleotide within 5 Å of protein. Modified nucleotides and source cofactors are retained. The related 7SOS/7SOV structures are distinct inputs from the same biological system, and the nine examples do not constitute a nonredundant survey of RNA recognition. LARP1 recognition of RNA 3*^′^* ends illustrates the biological setting ^52^.

Small-molecule docking to structured nucleic acids supports binding-mode analysis for RNA-targeted compounds ^53^. The evaluation set contains 124 RNA/DNA–ligand complexes with fixed ligand identities. Preparation requires a nucleic acid receptor and one connected, chemically valid ligand. One of the 125 initial inputs, 1Y27, is excluded because its nominal GDP ligand is covalently continuous with RNA. The retained set is not claimed to reproduce a complete public benchmark.

Each case has forty saved candidates generated with 400 sampling steps, flexible context, scoring, and postprocessing, giving 360 ssRNA poses and 4,960 ligand poses. All have complete atom mappings and available RMSDs. We analyze these saved structures without additional sampling or refolding.

Pose evaluation fits the receptor alone and measures the generated molecule under the same rigid transformation; the generated molecule receives no independent superposition. For ssRNA, all corresponding protein C*α* atoms are anchors, and RMSD includes every native-observed RNA heavy atom. Residues are matched by fixed component sequence and order, then by atom name and element, including modified components. Unobserved terminal phosphates and extra output terminal atoms do not enlarge the reference atom set. For ligands on nucleic acids, the anchors are observed P atoms in RNA/DNA linking residues. Ligand-heavy-atom RMSD uses the minimum over chemical graph symmetries consistent with the fixed ligand identity.

Top1 is the RMSD of the highest PartiScore-ranked candidate. Top5 best is the lowest RMSD among the first five ranks; Oracle is the lowest among all forty. Both minima use the native reference retrospectively and therefore describe sampled pose availability rather than prospective ranking. No binary docking threshold is applied to these two sets.

Per-case ssRNA distributions distinguish pose coverage from candidate ranking (Fig. S11). In 7V2Z, RMSD falls from 13.90 Å for Top1 to 1.44 Å for Top5 best, indicating that a close pose was sampled but not ranked first. By contrast, the Oracle RMSDs for 7UU4 and 8EDJ remain above 17 Å, indicating limited pose recovery within their forty candidates. Rank-one examples 7TUV and 7SOS illustrate bound placements at 1.95 and 2.12 Å RNA-heavy-atom RMSD (Fig. S13).

Ligand docking to nucleic acid receptors has a Top1 median of 1.203 Å across all 124 cases (Fig. S12). The three rank-one examples in Fig. S14 show the receptor environment and the reference/prediction overlay for A2F, SAM, and FMN ligands.

**Figure S11.**
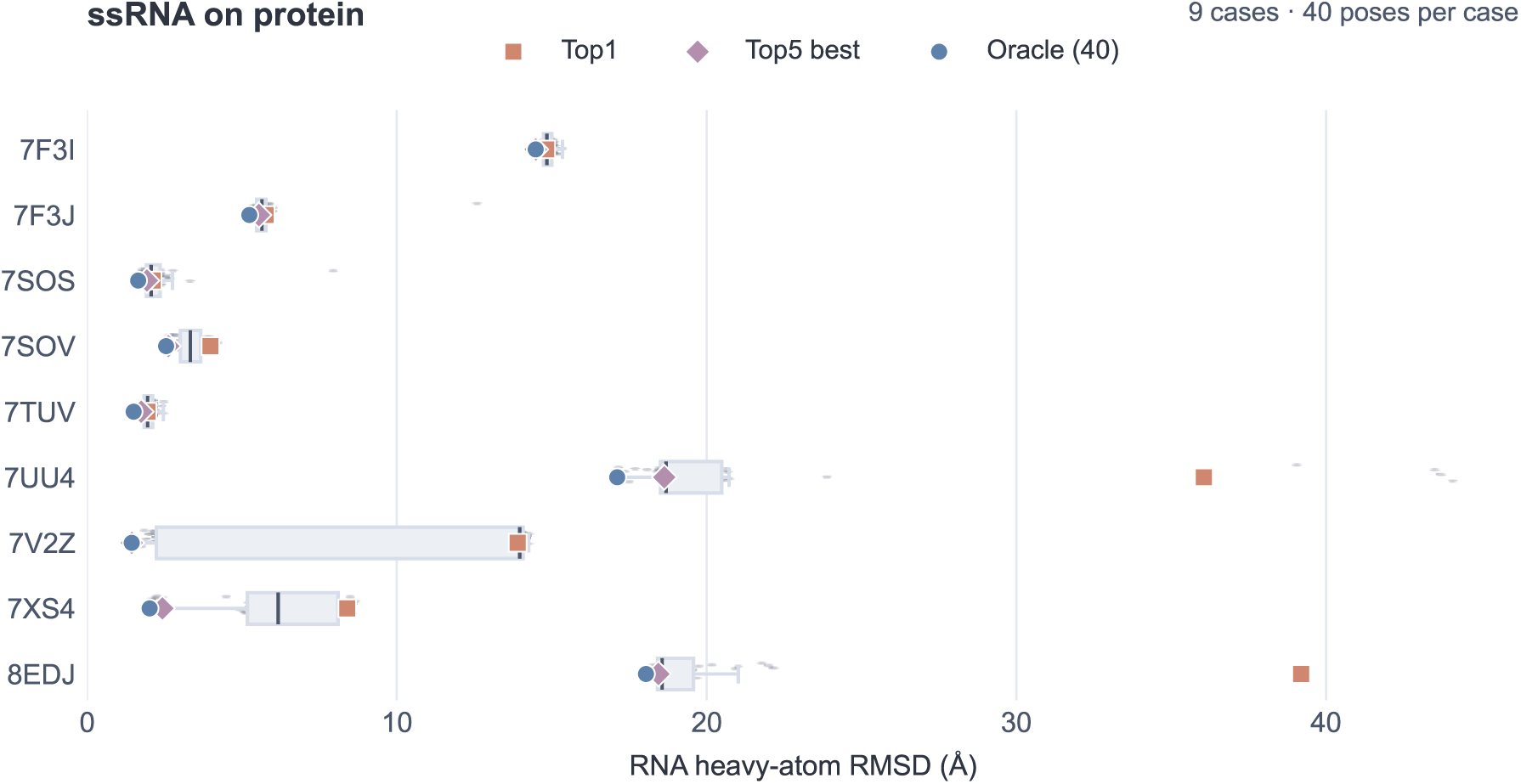
ssRNA docking to proteins. All nine cases have forty poses. Points show individual poses; boxes indicate the interquartile range, median and 1.5-IQR whiskers. Colored markers identify Top1, Top5 best and Oracle. RNA-heavy-atom RMSD is measured after fitting receptor C*α* atoms.

**Figure S12.**
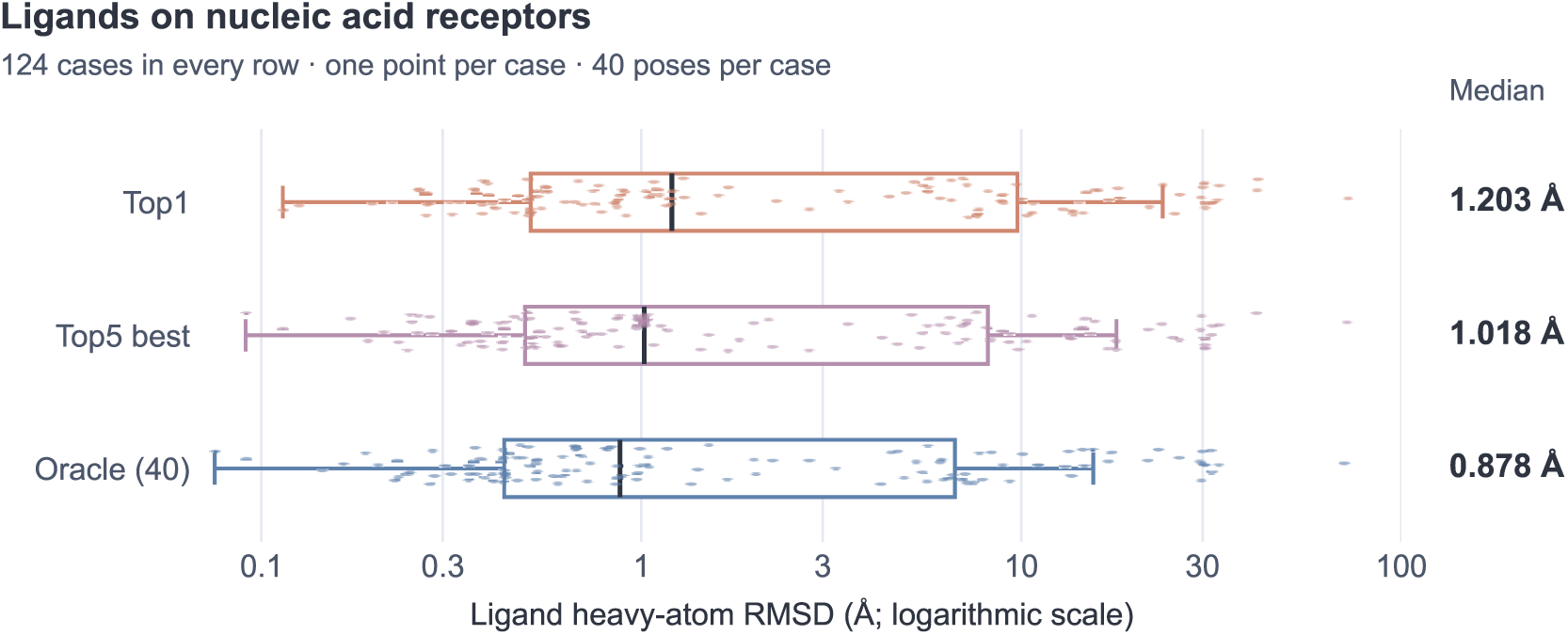
Ligand pose RMSD on nucleic acid receptors. Each row shows all 124 cases for Top1, Top5 best or Oracle among forty candidates. Boxes indicate the interquartile range, median and 1.5-IQR whiskers. Numbers at right are medians in Å. Ligand-heavy-atom RMSD uses symmetry-aware correspondence after receptor-P fitting.

**Figure S13.**
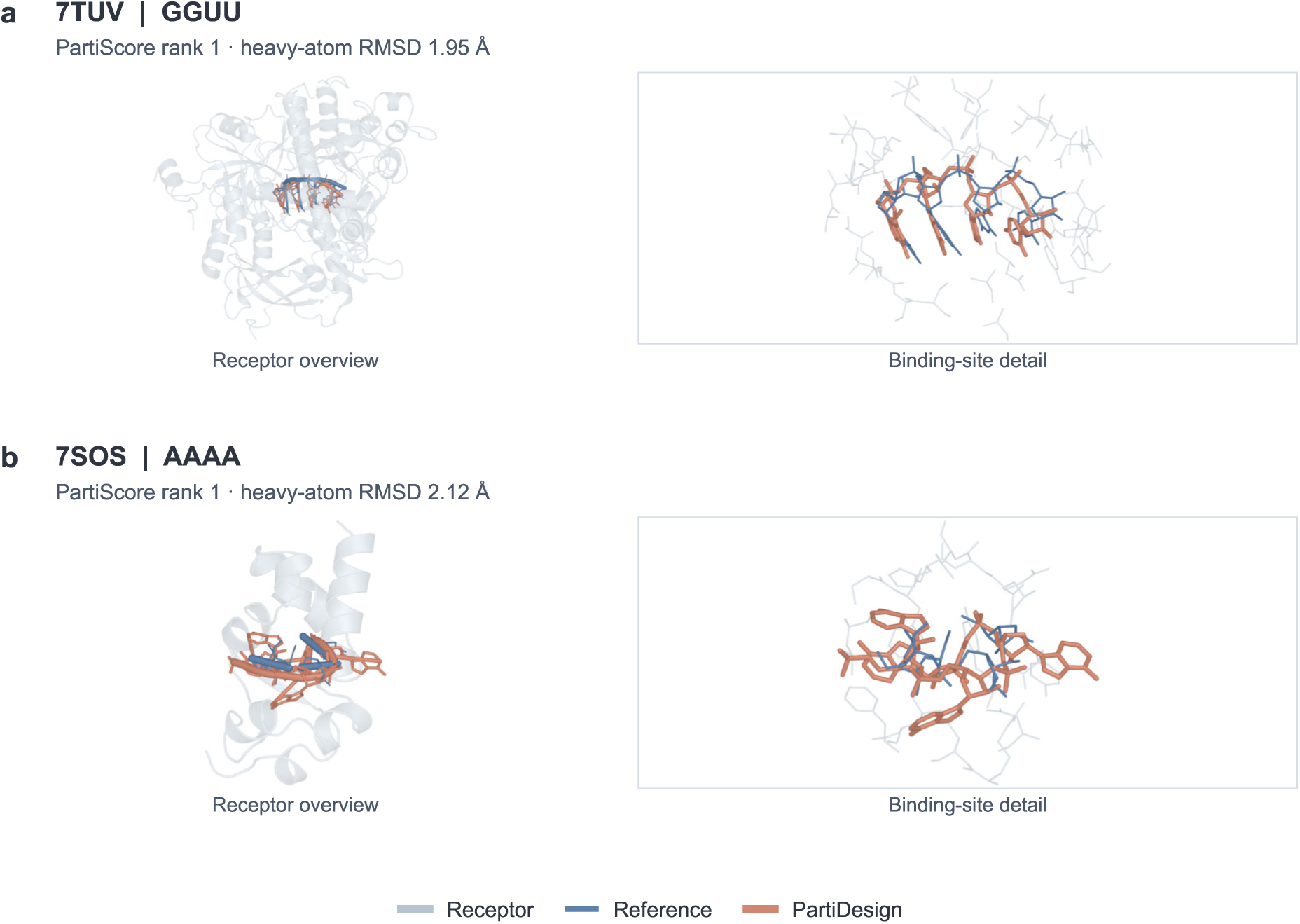
Selected rank-one ssRNA docking examples. Receptor overviews locate the RNA within the complex; enlarged views compare the experimental reference (blue) with the saved refined PartiDesign pose (orange). 7TUV and 7SOS contain the fixed GGUU and AAAA fragments, respectively. Labels report complete native-observed RNA-heavy-atom RMSD after receptor-C*α* fitting, with no RNA self-fit. These examples illustrate individual placements; Fig. S11 reports all nine cases.

**Figure S14.**
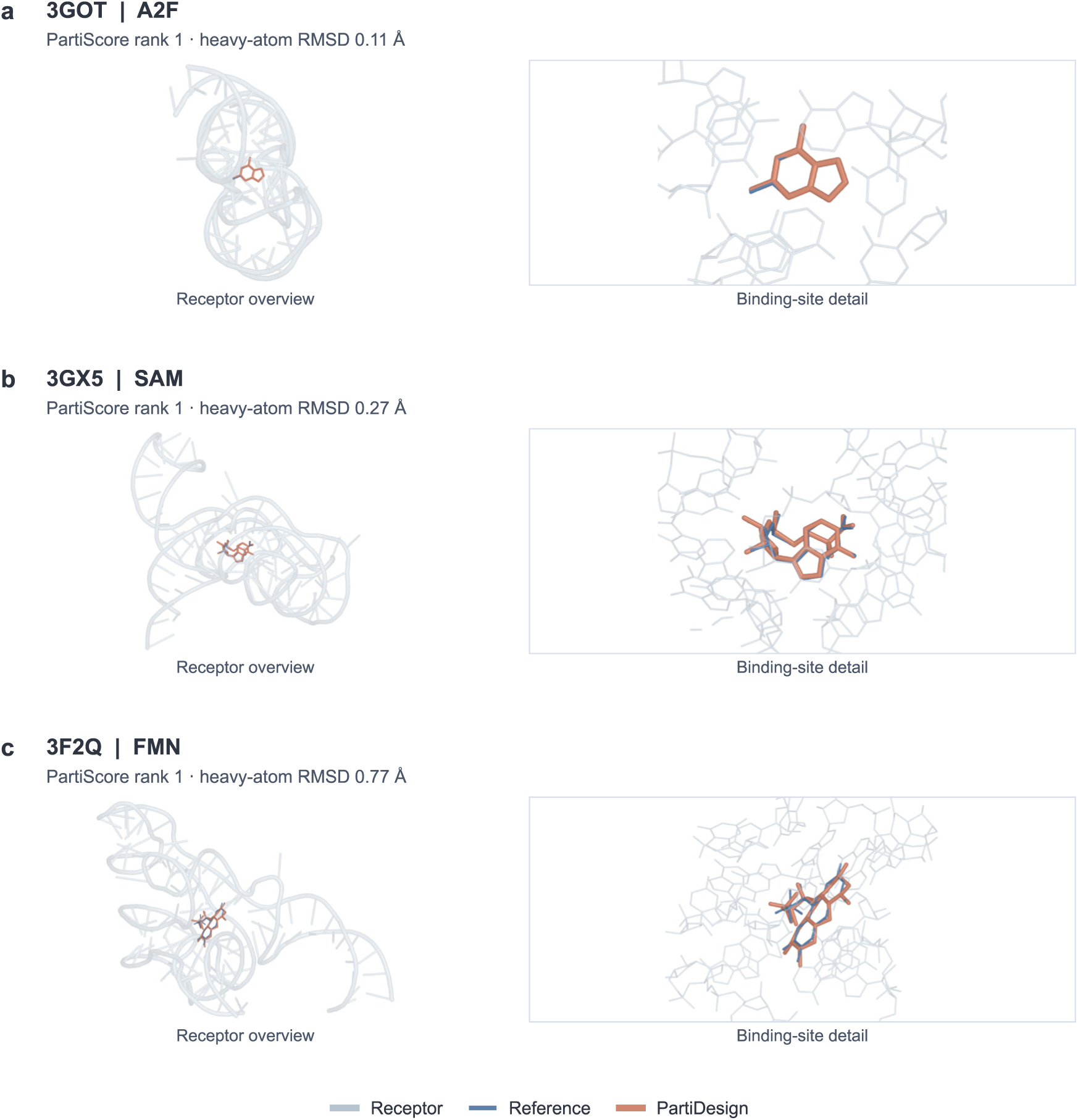
Selected rank-one ligand poses on RNA receptors. 3GOT (A2F), 3GX5 (SAM), and 3F2Q (FMN) compare saved refined PartiDesign candidates (orange) with the reference ligands (blue). The pale receptor overview establishes the binding site; the enlarged view shows the local RNA environment. Labels report symmetry-corrected ligand-heavy-atom RMSD after receptor-P fitting, with no ligand self-fit. These illustrative examples do not replace the complete 124-case distribution in Fig. S12.

### G.2 Protein–ligand docking on the PoseBusters-derived set

PoseBusters assesses physical validity alongside pose accuracy ^54^. The comparison contains 308 common protein–ligand cases, with forty poses per case from PartiDesign, RoseTTAFold3 (RF3), and DynamicBind ^55,56^. RF3 denotes the structure predictor and is distinct from RFD3. PartiDesign and RF3 receive native-ligand-centered 10 Å pockets, whereas DynamicBind receives full receptors; validity checks use full conditioning receptors. These different input scopes limit direct method attribution.

Symmetry-corrected ligand-heavy-atom RMSD is measured in the receptor frame without an independent ligand fit. RMSD-only passing requires RMSD ≤ 2 Å across all 308 cases. Joint passing requires the same candidate to pass this gate and all 27 saved PoseBusters checks, covering chemical consistency, intramolecular geometry and energy, proximity, clashes, and volume overlap. Joint evaluation uses the fixed intersection of 296 cases with complete flags for every candidate of every method. The twelve other cases remain in the RMSD-only analysis. Saved Boolean results support joint classification, but the records do not support a complete replay of every underlying continuous check.

PartiDesign uses refined-structure PartiScore ranks, RF3 its saved ranking score, and DynamicBind its output rank. RF3 score ties are resolved by source-sample order; ties occur at the Top1 cutoff in 205 cases and the Top5 cutoff in 140. Top1/Top5 ask whether any of the first one/five candidates passes; Oracle@40 asks whether any of forty passes. The case sets stay fixed as *k* changes.

**Figure S15.**
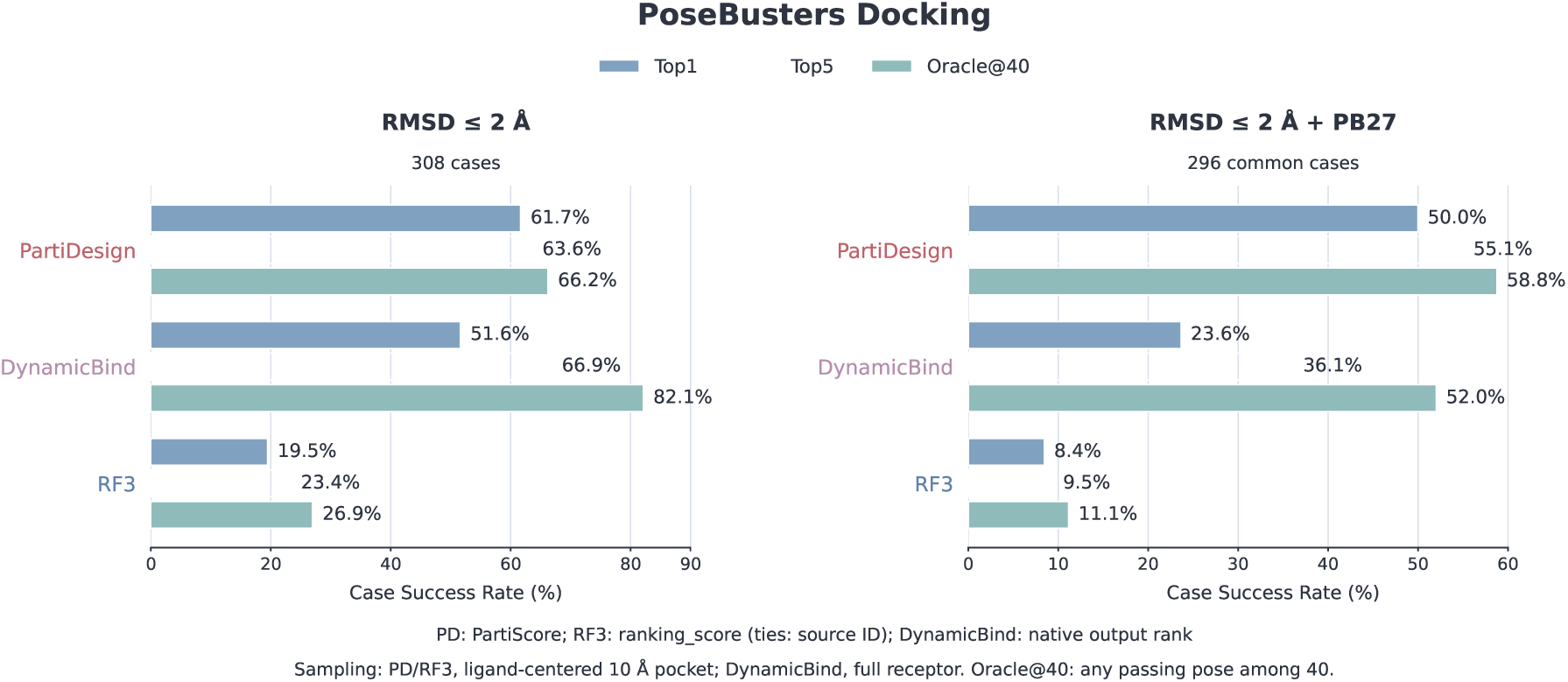
Ranked pose recovery on the PoseBusters-derived set. RMSD-only uses 308 cases and the inclusive *≤* 2 Å gate. Joint evaluation uses 296 common cases and additionally requires all saved PB27 checks for the same candidate. Each method supplies forty poses. Top1/Top5 follow each method’s ranking; Oracle@40 uses all forty. Pocket-conditioned PartiDesign/RF3 and full-receptor DynamicBind have different input scopes.

PartiDesign achieves RMSD-only Top1/Top5/Oracle counts of 190/196/204 out of 308; RF3 reaches 60/72/83 and DynamicBind 159/206/253. On the 296-case joint set, the corresponding counts are 148/163/174, 25/28/33, and 70/107/154. PartiDesign’s joint Top1 is 50.00%, while DynamicBind’s RMSD-only Oracle is 82.14%. Method ordering thus depends on whether evaluation emphasizes the first ranked physically plausible pose or coverage across sampled poses, together with the differing input protocols.

### G.3 Cyclic-peptide docking examples

Fixed-identity cyclic-peptide docking tests both bound-pose recovery and accommodation of covalent closures. Four PartiDesign examples illustrate distinct topologies drawn from CPSet^57^: 6D40 combines head-to-tail cyclization and a disulfide; 4X1Q has disulfide cyclization; 2YQ6 has an MK8 staple; and 5C3F has an NLE staple. Inputs preserve the peptide sequence, nonstandard residue identities, and specified covalent connections.

Table S9 reports Top1 and the best of forty saved refined candidates for each example. The displayed candidate is chosen retrospectively by the lowest peptide-backbone RMSD among forty, with its PartiScore rank stated explicitly. RMSD is measured over corresponding peptide N/C*α*/C/O atoms after fitting receptor C*α* atoms alone. The peptide receives no independent fit, so the metric retains both conformation and binding-site placement errors. These selected structures demonstrate individual geometries and do not estimate success across a larger cyclic-peptide set.

**Table S9.** Four cyclic-peptide docking examples. RMSDs are in Å for the peptide N/C*α*/C/O backbone after receptor-C*α* fitting. Each example has forty saved refined candidates. The last column identifies the PartiScore rank of the displayed best-of-forty candidate.

| PDB | Covalent topology | Top1 | Best of 40 | Displayed rank |
|---|---|---|---|---|
| 6D40 | Head-to-tail + disulfide | 0.401 | 0.303 | 3 |
| 4X1Q | Disulfide | 0.565 | 0.479 | 8 |
| 2YQ6 | MK8 staple | 1.222 | 1.222 | 1 |
| 5C3F | NLE staple | 0.730 | 0.620 | 26 |

**Figure S16.**
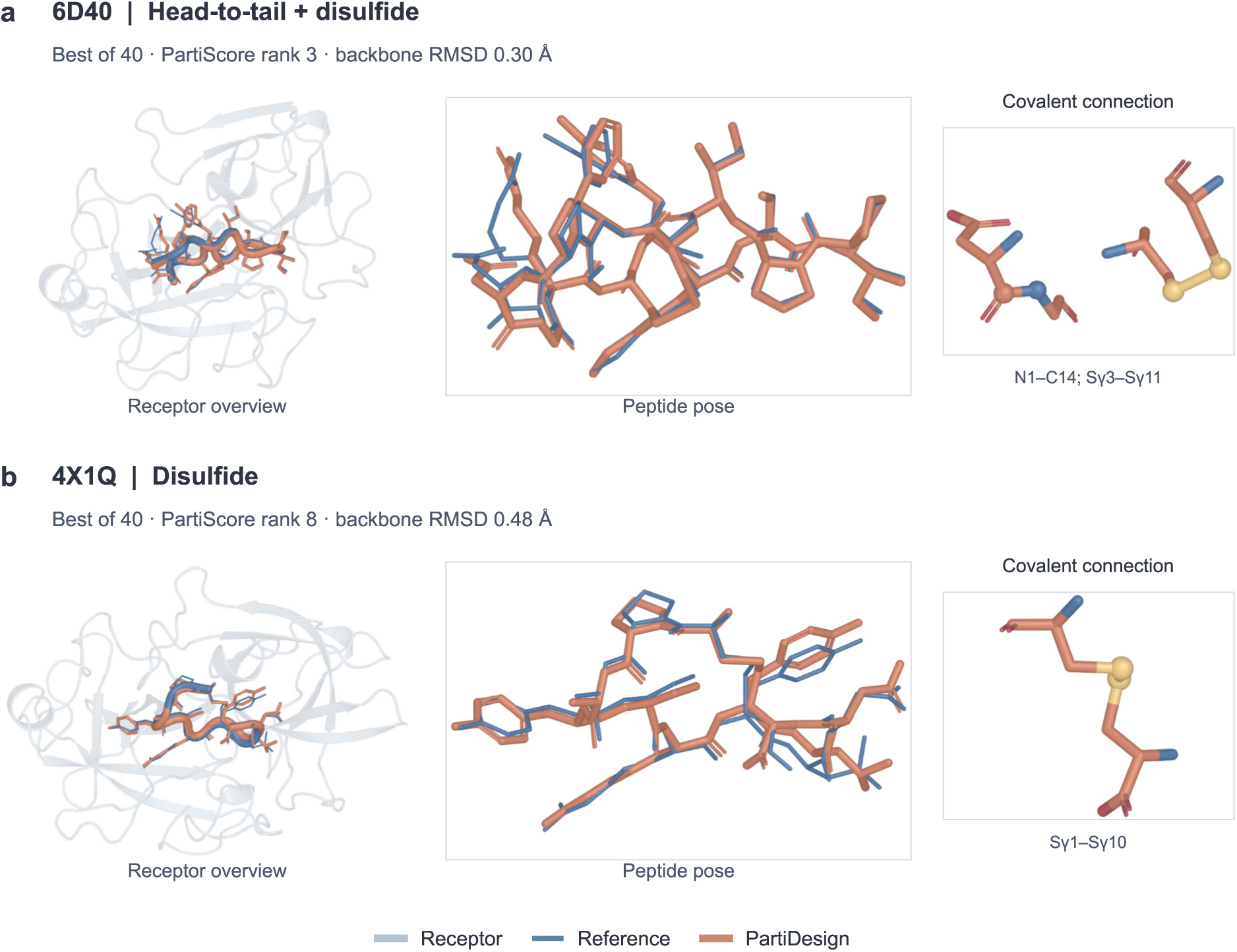
Head-to-tail and disulfide docking examples. Receptor overviews and peptide overlays compare the experimental reference (blue) with the saved refined PartiDesign candidate (orange). The displayed 6D40 and 4X1Q candidates are selected by best backbone RMSD among forty, with PartiScore ranks 3 and 8. Connection insets show the saved N1–C14 head-to-tail and Cys3–Cys11 disulfide in 6D40, and Cys1–Cys10 disulfide in 4X1Q. Labels report receptor-aligned peptide-backbone RMSD.

**Figure S17.**
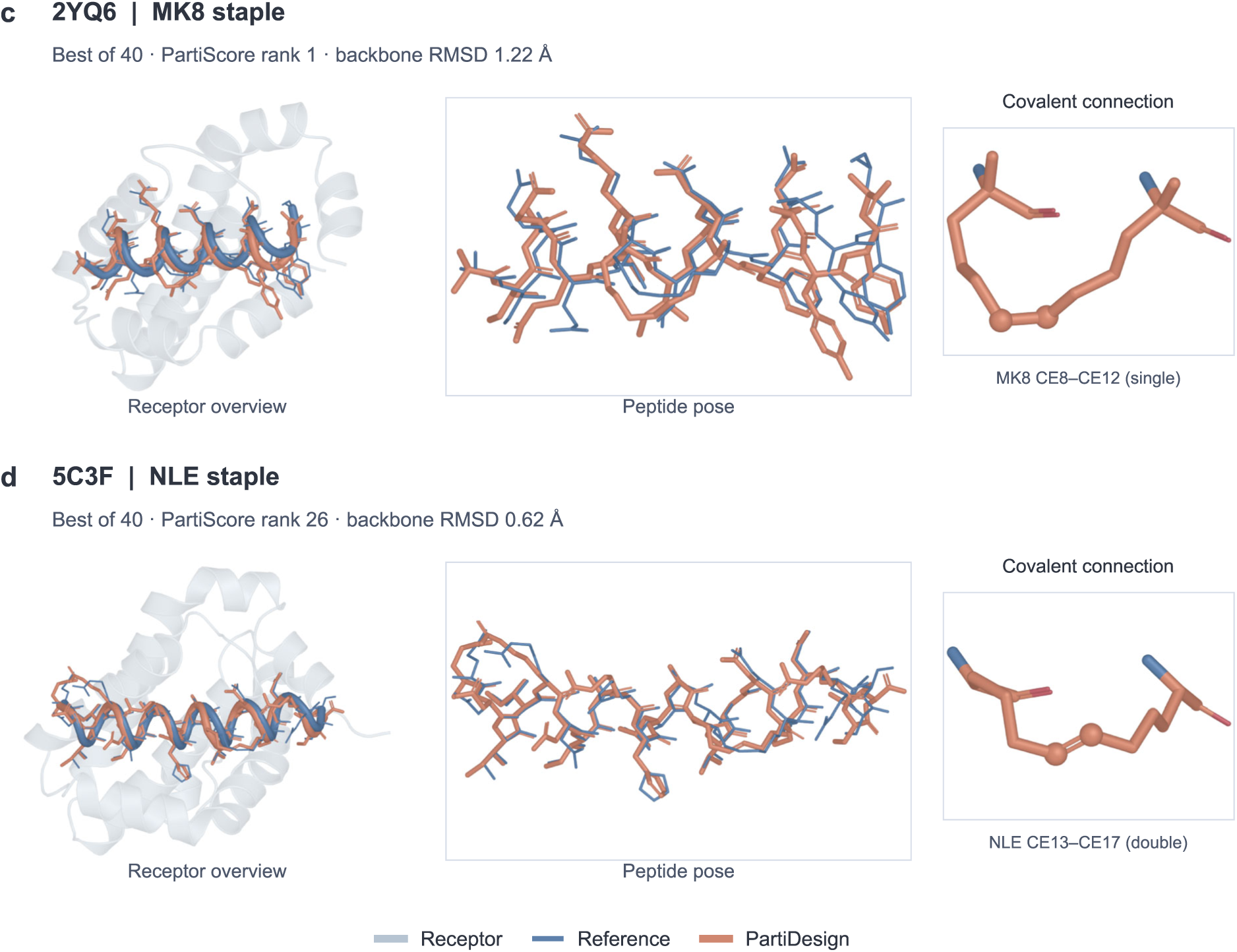
Stapled-peptide docking examples. Reference (blue) and saved refined PartiDesign (orange) poses are shown with receptor overviews, peptide overlays, and enlarged covalent connections. The best-of-forty 2YQ6 and 5C3F candidates have PartiScore ranks 1 and 26. The saved 2YQ6 MK8 CE8–CE12 connection is a single bond; the 5C3F NLE CE13–CE17 connection is a double bond. Labels report peptide N/C*α*/C/O backbone RMSD after receptor-C*α* fitting, without a peptide self-fit.

